# Probing small ribosomal subunit RNA helix 45 acetylation across eukaryotic evolution

**DOI:** 10.1101/2021.11.30.470322

**Authors:** Bortolin-Cavaillé Marie-Line, Quillien Aurélie, Thalalla Gamage Supuni, Justin M. Thomas, Sas-Chen Aldema, Sharma Sunny, Plisson-Chastang Célia, Vandel Laurence, Blader Patrick, Denis L.J. Lafontaine, Schwartz Schraga, Jordan L. Meier, Cavaillé Jérôme

## Abstract

NAT10 is an essential enzyme that catalyzes the formation of N^4^-acetylcytidine (ac^4^C) in eukaryotic transfer RNA (tRNA) and 18S ribosomal RNA (rRNA). Recent studies in human cells suggested that rRNA acetylation is dependent on SNORD13, a non-canonical box C/D small nucleolar RNA (SNORD) predicted to base-pair with 18S rRNA via two antisense elements. However, the selectivity of SNORD13-dependent cytidine acetylation and its relationship to NAT10’s essential function in pre-rRNA processing remain to be defined. Here, we used CRISPR-Cas9 genome editing to formally demonstrate that SNORD13 is required for acetylation of a single cytidine residue of human and zebrafish 18S rRNA. In-depth characterization revealed that SNORD13-dependent ac^4^C is dispensable for yeast or human cell growth, ribosome biogenesis, translation, and the development of multicellular metazoan model organisms. This loss of function analysis inspired a cross-evolutionary survey of the eukaryotic rRNA acetylation ‘machinery’ that led to the characterization of many novel SNORD13 genes in phylogenetically-distant metazoans and more deeply rooted photosynthetic organisms. This includes an atypical SNORD13-like RNA in *D. melanogaster* which appears to guide ac^4^C to 18S rRNA helix 45 despite lacking one of the two rRNA antisense elements. Finally, we discover that *C. elegans* 18S rRNA is not acetylated despite the presence of an essential NAT10 homolog. Altogether, our findings shed light on the molecular mechanisms underlying SNORD13-mediated rRNA acetylation across the eukaryotic tree of life and raise new questions regarding the biological function and evolutionary persistence of this highly conserved rRNA base modification.

## Introduction

N4-acetylcytidine (ac^4^C) in eukaryotic and archaeal rRNA was first documented in the 1970’s, but the catalytic machinery and specific sites of this base modification have been defined only recently (1, 2). In eukaryotes, ac^4^C occurs at two positions within rRNA (helix 34 and 45) where it is catalyzed by the nucleolar GCN5-related RNA acetyltransferase NAT10, also referred to as Kre33 or Rra1 in yeast (3–5). In addition, recent studies showed that certain archaeal species harbour >100 ac^4^C residues in their rRNA, including at helix 45, placing cytidine acetylation on par with prevalent nucleoside modifications such as pseudouridines and 2’-O-methylations in these organisms (6, 7). Beside its involvement in rRNA modification, NAT10/Kre33 (assisted by its specific co-factor THUMPD1/Tan1) can modify C12 of serine and leucine tRNAs (5, 8) and has been suggested to modify eukaryotic mRNAs (9, 10). Orthogonal validation has, however, called into question the involvement of NAT10 in mRNA modification (6). In addition to modifying RNA, NAT10 may also be implicated in protein acetylation (11–13). Moreover, NAT10 is essential for pre-rRNA processing reactions leading to synthesis of 18S rRNA (5), and unsurprisingly it was associated with human diseases (14, 15). The importance of helix 45 ac^4^C for cell homeostasis has not been addressed.

In yeast, the ability of Kre33 to specifically acetylate 18S rRNA was demonstrated to rely on two box C/D small nucleolar RNAs (SNORD), namely snR4 and snR45, which form imperfect base pairing interactions surrounding the targeted cytidine. Indeed, knockout of snR4 and snR45 led to the specific disappearance of 18S rRNA acetylation at SSU-C1280 (helix 34) and SSU-C1773 (helix 45), respectively, while leaving tRNA acetylation unaffected (16). SNORDs are associated with four core proteins (SNU13, NOP56, NOP58, and Fibrillarin) and form nucleolar ribonucleoprotein (RNP) complexes known to mediate site-specific ribose methylation of rRNA, U6 spliceosomal snRNA and tRNAs; some of them are also involved in early nucleolar pre-rRNA processing (17). The possible contribution of snoRNA-associated proteins in rRNA acetylation remains undefined. Of note, SNORD13 (formerly U13) displays two rRNA complementarity tracts reminiscent of those present on snR45 (**Figure 1**), strongly implicating it as a human counterpart (5, 18). Consistent with this notion, inhibiting SNORD13 expression in a cancer cell line using antisense oligonucleotides causes a ∼50% decrease in levels of ac^4^C in 18S rRNA (5). However, the nucleotide-specificity of this inhibition and the residual level of acetylation did not afford to easily assess the impact of the modification on cell growth or ribosome biogenesis. Finally, three SNORD13-like RNA species were also reported in *A. thaliana* (snoR105, snoR108 and snoR146) but their roles in rRNA acetylation still awaits investigation (19).

**Figure 1.**
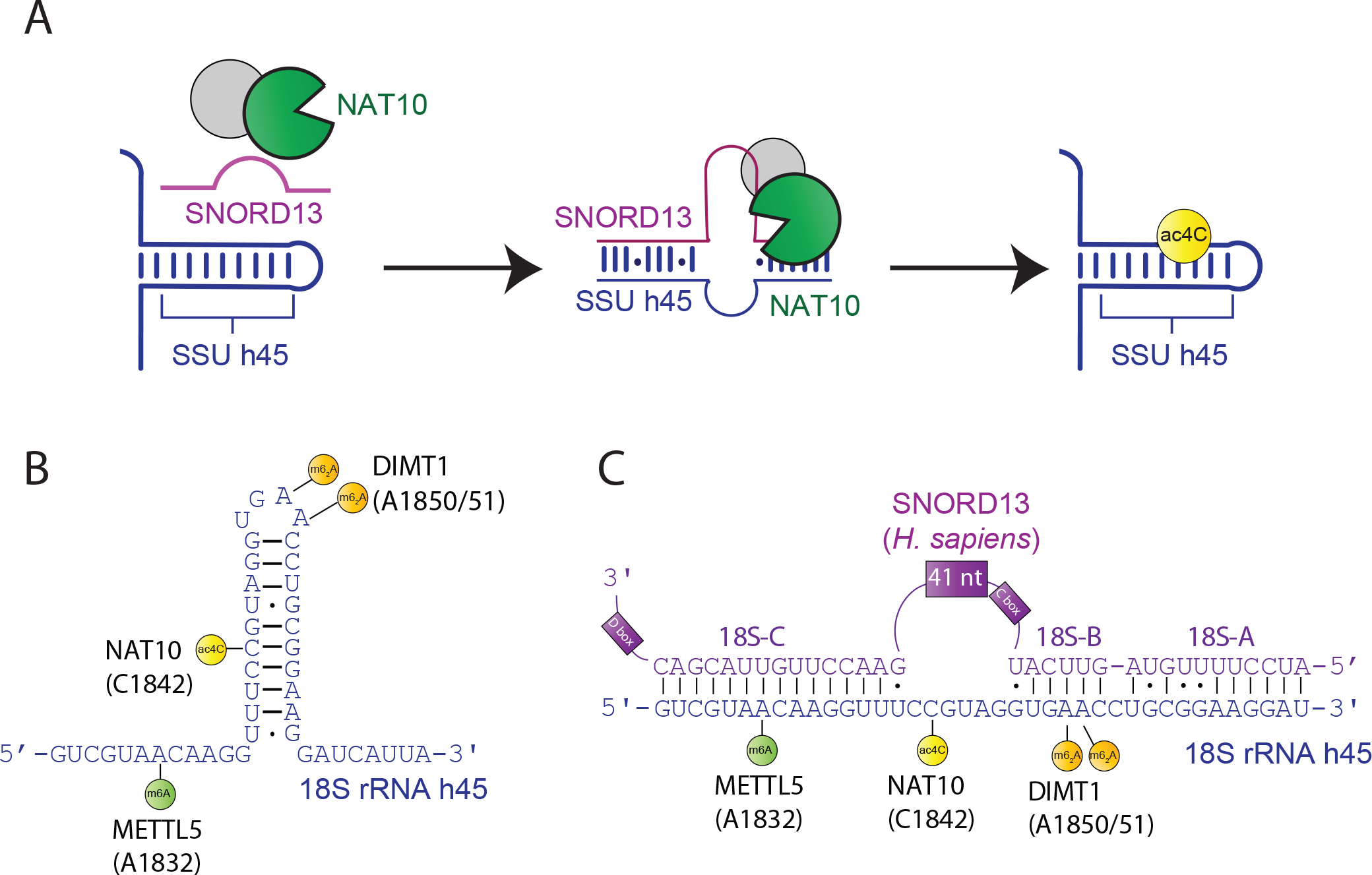
Schematic of SNORD13-dependent ribosomal RNA acetylation. **A)** Through an incompletely defined antisense-mediated mechanism, human SNORD13 assists the RNA cytidine acetyltransferase NAT10 in modifying C1842 in the universally conserved helix 45 at the 3’ extremity of 18S rRNA. **B)** Helix 45 is enriched with post-transcriptional RNA modifications. RNA modifying enzymes and their modified nucleotides (lollipops) are indicated: SSU-ac^4^C1842 (yellow), SSU-m^6^A1832 (green) and SSU-m_2_ A1850 and SSU-m_2_ A1851 (orange). **C)** Human SNORD13 base-pairs with 18S rRNA sequences on each side of the cytidine to be acetylated. Note that SNORD13-rRNA duplexes tolerate G-U wobble base-pairs, mismatches or even bulged nucleotides and the length of the intervening loops, from either rRNA or SNORD side, also differ from one organism to another (**Supplementary figure S1**).

Outside their potential to form two imperfect base-pairing interactions with 18S rRNA, vertebrate SNORD13 and its *S. cerevisiae* and *A. thaliana* homologs are not highly conserved. This explains why, although SNORD13 was discovered thirty years ago (18, 20), its relationship with snR45 was only recognized recently (16). This suggests two distinct evolutionary hypotheses: either SNORD-mediated rRNA acetylation emerged early in a common ancestor of plants, fungi and animals and diverged thereafter, or alternatively, SNORD13 arose independently in these different lineages via convergent evolution. Irrespective of the scenario, the possibility also exists that SNORD13 has been lost in some lineages and, accordingly, helix 45 acetylation may rely on a stand-alone enzyme, or no longer be present in rRNA in certain species. This poses the question as to how SNORD-guided cytidine acetylation is conserved across the eukaryotic tree of life, and what cross-organismal commonalities in the nucleic acid acetylation machinery and its rRNA substrates govern this process.

Here, we use CRISPR-Cas9 genome editing to delete SNORD13 in human cells, confirming the hypothesis that this snoRNA is specifically required for acetylation of a single cytidine residue in small subunit ribosomal RNAs (SSU-ac^4^C1842 located in rRNA helix45). Using this cell line and related model systems, we further establish that loss of SSU-ac^4^C1842 does not lead to obvious defects in terms of cell growth, protein synthesis, ribosome biogenesis, or the development of metazoan model organisms (zebrafish, flies). Finally, considering that Kre33/NAT10 is essential for pre-rRNA processing (5) while ac^4^C in helix 45 appears dispensable, we surveyed the conservation of the ribosomal RNA acetylation machinery across the eukaryotic tree of life. This cross-evolutionary analysis revealed that SNORD-mediated rRNA acetylation is widely conserved in Metazoans, but defined *C. elegans* as a multicellular organism in which rRNA acetylation has been lost. Our findings illuminate the molecular mechanisms underlying SNORD13-mediated rRNA acetylation across evolution and raise new questions regarding the role of highly conserved SSU-ac^4^C1842.

## Results

### Genetic deletion of human SNORD13 gene causes loss of SSU-ac^4^C1842

As a first step to probe directly the functional role of SNORD13 in rRNA modification, the human SNORD13 gene was inactivated in the haploid human chronic myelogenous leukemia-derived HAP1 cell line (21). In order to avoid potential transcriptional interference with the adjacent TTI2 gene, the 3’-end region of SNORD13 was targeted for CRISPR-Cas9-mediated deletion (**Figure 2A**). PCR analysis identified three independent SNORD13-deficient clones (#12/#20/#30) harboring the expected deletion (not shown), which rendered SNORD13 undetectable by Northern blot **(Figure 2B)**. Three additional clones (#13/#19/#22) were identified that were exposed to CRISPR/Cas9 without any deletion at SNORD13, providing a set of matched control cell lines subjected to identical clonal selection procedures and/or putative off-target effects. Unless otherwise specified, in subsequent experiments SNORD13-KO and WT are used to refer to a stoichiometric mixture of these three SNORD13-deficient and SNORD13-expressing clones, respectively.

**Figure 2.**
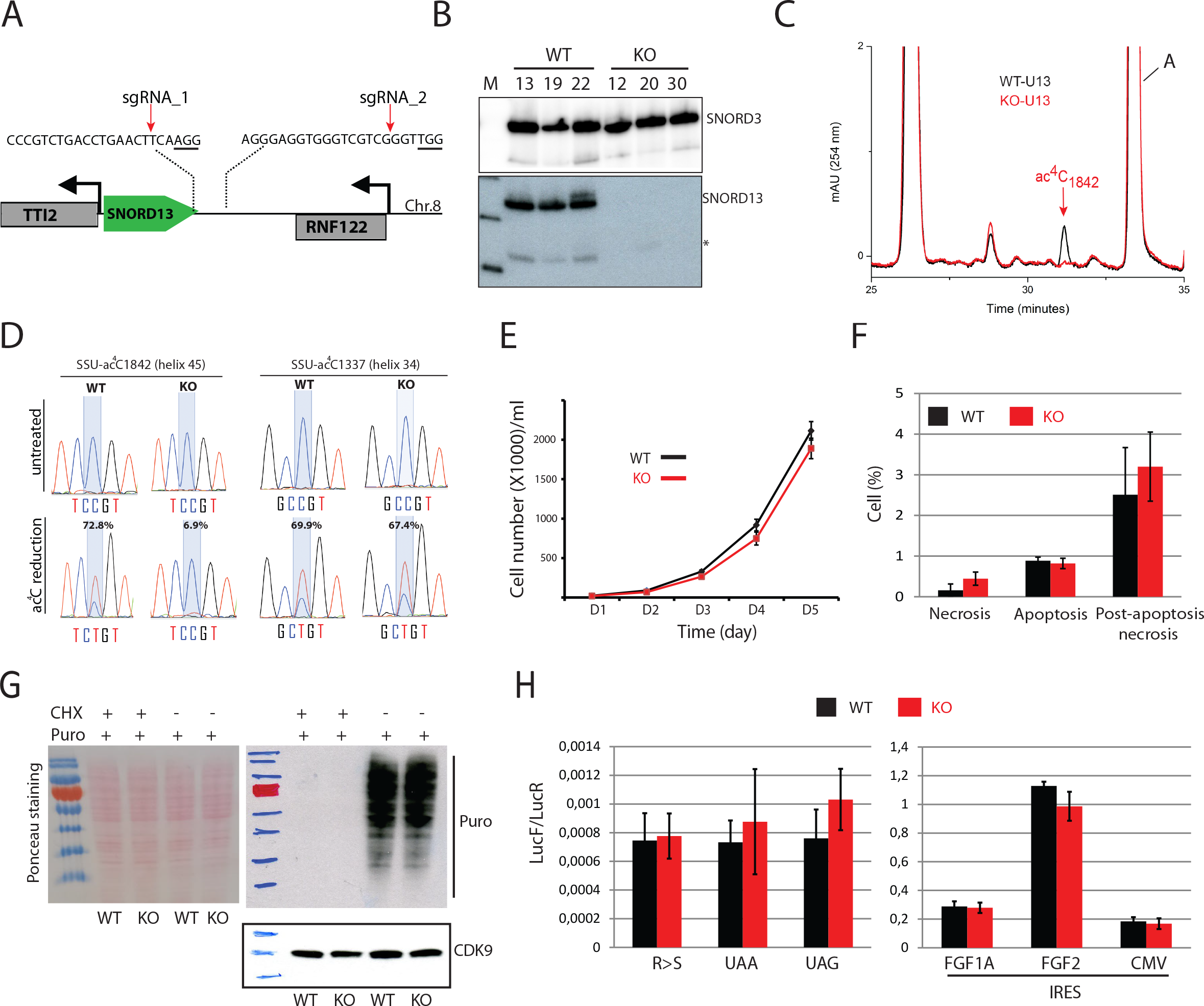
The specific lack of SNORD13-mediated acetylation in helix 45 does not alter cell growth or translation. **A)** Schematic representation of SNORD13 locus at human chromosome 8. The relative location of the two DNA sequences targeted by sgRNAs is shown (PAM sequence is underlined). **B)** Northern blot showing that SNORD13 is no longer detected in clones bearing deletions. * indicate truncated SNORD13 form which is routinely detected and corresponding to RNA species whose 5’-end is positioned 5-6 nucleotides upstream of the C-box (not shown). **C)** RP-HPLC chromatograms of nucleosides obtained from helix 45 of WT (black) and SNORD13 (red) cells. Peak corresponding to SSU-ac^4^c1842 is indicated. **D)** Sanger DNA sequencing of RT-PCR products obtained after borohydride treatments of total RNA extracted from WT and SNORD13-KO cells. % of misincorporation (C-to-U) at SSU-C1842 (helix 45) and SSU-C1337 (helix 34) is indicated below each electropherogram. **E)** Growth curves of WT and SNORD13-KO cells as judged by cell counting (n=6 for each genotype). **F)** Apoptosis levels determined by flow cytometry using propidium and Annexin V staining of WT and SNORD13-KO cells (n=2 for each genotype). **G)** SUnSET assay. Incorporation of puromycin into the elongating peptide was assayed in WT and SNORD13-KO cells (western blotting with anti-puromycin antibodies). Ponceau S staining of the membrane and detection of CDK9 (western blotting) were used as gel loading controls. Cells treated with the translation inhibitor cycloheximide (CHX) were also used as negative controls. **H)** Luciferase reporter gene assays. WT and SNORD13-KO cells were transiently co-transfected by two plasmids expressing Renilla luciferase gene (LucR; internal control) or Firefly luciferase gene (lucF; translation reporter) carrying either an in-frame stop-codons (UAA, UAG), a detrimental mutation (R-to-S) or an IRES from FGF1A, FGF2 and CMV. Histograms show the normalized luciferase activity (lucF-to-lucR ratio). Data are expressed as mean +/-s.e.m. and represent 5-6 independent experiments with triplicate measurements.

To define the site-specific activity of SNORD13, we took a dual approach to analyze our mutant and WT cell lines. First, we directly measured ac^4^C levels in helix 45 using a mung bean nuclease protection assay coupled to reversed phase high performance liquid chromatography (RP-HPLC, (5)). Briefly, total RNA extracted from SNORD13-KO and WT cells were hybridized with an antisense DNA oligonucleotide overlapping helix 45 and its flanking sequences and were treated with mung bean nuclease. Protected rRNA fragments were then gel-purified and nucleoside hydrolysates obtained after nuclease P1 digestions were subjected to RP-HPLC analysis. As shown in **Figure 2C**, cytosine acetylation at helix 45 was completely abolished in SNORD13-KO cells as compared to WT. As orthogonal validation, RNA isolated from SNORD13-KO and WT cells was analyzed using ac^4^C sequencing chemistry (22). In this approach total RNA is treated with sodium cyanoborohydride under acidic conditions, causing chemical reduction of the acetylated residue. This reduced nucleobase causes misincorporation of non-cognate NTPs during reverse transcription (RT), enabling ac^4^C sites to be detected as C-to-T mutations upon cDNA sequencing. Applying this approach to the two cell lines, we detected SNORD13-dependent misincorporation at SSU-ac^4^C1842 but not at SSU-ac^4^C1337 **(Figure 2D).** These experiments validated our knockout strategy and unambiguously defined SNORD13 as a human counterpart of yeast snR45 required for guiding formation of ac^4^C at the terminal stem-loop of human 18S rRNA.

### SNORD13 deletion does not disrupt cell growth, pre-rRNA processing or protein synthesis in human cells

Next, we assessed the phenotypic consequences of SNORD13 deletion and/or SSU-ac^4^C1842 loss. Analysis of proliferation by cell counting revealed that loss of SNORD13 did not impact cell growth significantly (**Figure 2E)** and that cell death rates and modalities were within normal range (**Figure 2F**). We next examined the levels of pre-rRNA precursors and mature rRNAs in SNORD13-KO cells by Northern blot **(supplementary Figure 2A-B),** RNase H mapping **(supplementary Figure 2C)** and pulse chase experiments **(supplementary Figure 2D).** These analyses revealed that loss of SNORD13 and/or SSU-ac^4^C1842 does not interfere with normal pre-rRNA processing. An involvement of SNORD13 and/or SSU-ac^4^C1842 on ribosomal subunit assembly and global translation was also investigated by performing polysomal analysis. In this assay, whole cytoplasmic extracts from SNORD13-KO and WT cells were subjected to sucrose density gradient fractionation (10-50%). As shown in **supplementary figure 2E**, the distribution and amplitude of peaks corresponding to 40S, 60S, 80S monosomes and polysomes were virtually identical in SNORD13-KO and WT cells. Moreover, detection of nascent protein production by puromycin labelling (SUnSET assay) revealed identical, translation-dependent incorporation patterns in SNORD13-KO and WT cells, again indicating that overall protein synthesis is unlikely to be severely impaired in SNORD13-deficient cells (**Figure 2G)**. To test whether SSU-ac^4^C1782 plays a more subtle role in fine-tuning aspects of ribosome function such as translational fidelity, we transfected WT and SNORD13-KO cells with firefly luciferase reporter genes bearing in-frame stop codons (UAA, UAG) or point mutations (R218S) that abrogate enzymatic luciferase activity. These reporters enable any decrease in translational accuracy to be detected via a corresponding increase in firefly luciferase activity. Our data indicate that stop codon read-through and amino acid misincorporation in SNORD13-KO cells was similar between WT and KO cells (**Figure 2H-left panel**). Finally, when using a set of Renilla-IRES-Firefly bicistronic vectors, we also showed that the efficiency of internal ribosome entry site (IRES)-mediated translation was apparently unaffected in SNORD13-KO cells **(Figure 2H-right panel)**. Overall, these findings suggest that SNORD13 and SSU-ac^4^C1842 are not essential for ribosome biogenesis or function; at least in standard laboratory growth conditions.

### SNORD13-deficient zebrafish embryos develop normally

In order to understand the biological relevance of SNORD13 at the whole organism level, a 189 bp-long, CRISPR-Cas9 mediated deletion overlapping the *Danio rerio* (zebrafish) *snord13* gene locus on chromosome 8 was generated (**Figure 3A**). Crossing of individual heterozygotes led to the generation of homozygous adult snord13 knockout fish of normal appearance (not shown). To exclude any maternal contribution, three different homozygous females were crossed with heterozygous males. As expected, *snord13* and ac^4^C in helix 45 were no longer detectable in RNA isolated from homozygous knockout fish, as judged by Northern blotting (**Figure 3B**) and ac^4^C sequencing (**Figure 3C**), respectively. Homozygous *snord13* individuals in the progeny did not present any apparent morphological defects at 30 hours post fertilization (30 hpf) and developed normally until adulthood (**Supplementary Figure 3A)**. To explore the possible occurrence of more subtle defects, such as activation of the nucleolar stress pathway in response to impaired ribosome biogenesis (23), we performed immunostaining against activated Caspase-3. No increase in apoptotic cells was observed in the developing brain **(Figure 3D-E)** or spinal cord **(Figure 3F-G)** of *snord13*-deficient embryos, as compared to WT. Accordingly, the level of expression of the pro-apoptotic *bcl2*, *bax* and *puma* genes, as judged by RT-qPCR, was not significantly altered in *snord13* mutant embryos (**supplementary Figure 3B**). Finally, we performed immunostaining in WT and mutant embryos for the pan neural marker HuC/D which labels newly differentiated neurons in different structures including olfactory epithelium, pineal gland or optic tectum and the spinal cord. Again, no obvious difference could be detected between WT and mutant embryos (**Figure 3D-G**). Beside central nervous system development, we also sought to explore whether snord13 mutant displayed defects in another process such as vasculogenesis. We found that the vasculature of the trunk which is composed of the dorsal aorta (DA), the cardinal vein (CV) and intersegmental vessels was also normal in 30 hpf snord13-deficient embryos, as revealed by *in situ* hybridization against the key vascular transcription factor *fli1a* (**Figures 3H-I**). Similarly, the expression levels of various markers of the neural system (*lhx1a*, *pax6a* and *otx5*) and of the dorsal aorta trunk vasculature (*cdh5* and *fli1a*) were unchanged between WT and snord13-KO embryos, validating the lack of phenotypes observed **(Supplementary Figure 3B**). In conclusion, neither SNORD13 nor 18S rRNA ac^4^C in helix 45 appear to play any essential roles in embryonic zebrafish development.

**Figure 3.**
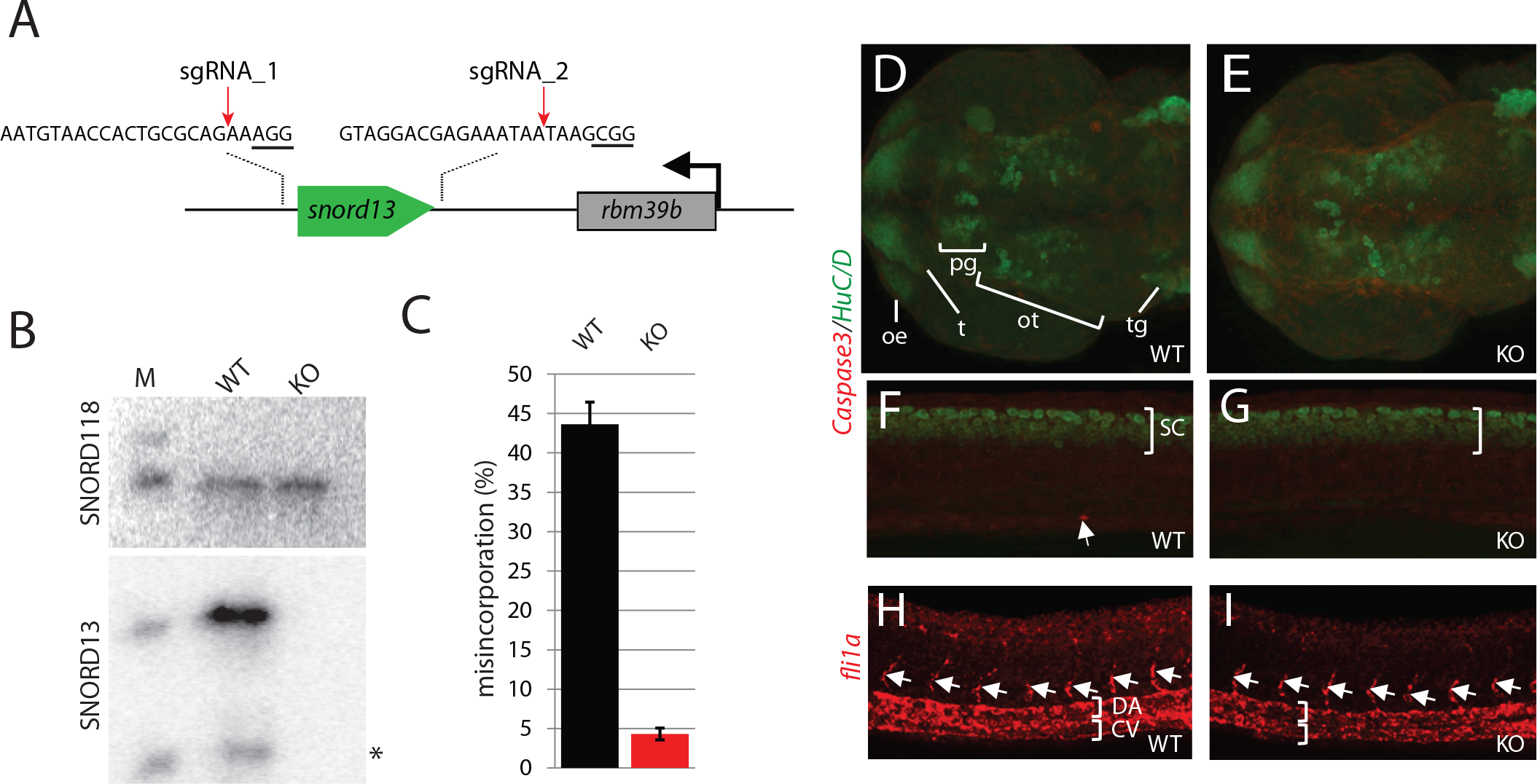
Phenotypic analyses of SNORD13-deficient zebrafish embryos. **A)** Schematic representation of the zebrafish *snord13* locus. The relative location of the two DNA sequences targeted by sgRNAs is shown (PAM sequence is underlined). **B)** Northern blotting showing that SNORD13 is no longer detected in embryos bearing deletion events. *: truncated SNORD13 form which is routinely detected. SNORD118 was used as a gel loading control. **C)** Histograms show percentage of misincorporation (C-to-U ratio) measured in borohydride-treated RNA samples prepared from WT and SNORD13-KO embryos (n= 3). **D-G** Confocal projections of WT (**D**, **F**) and SNORD13-KO (**E**, **G**) embryos after immunostaining against cleaved Caspase-3 (red) and the pan neural marker HuC/D (green). **D-E**: Dorsal view of the zebrafish brain at 30 hpf, neurons from the olfactory epithelium (oe), tectum (t), pineal gland (pg), optic tectum (ot) and trigeminal ganglion (tg) can be detected. **F-G:** Lateral view of the spinal cord (sc) at 30 hpf. **H-I** Confocal projections of WT (**H**) and SNORD13-KO (**I**) embryos after *in situ* RNA hybridization using antisense riboprobes directed against *fli1a* transcripts. White brackets indicate the dorsal aorta (DA) and the cardinal vein (CV) and the white arrows point to intersegmental vessels. Pictures are representative of three independent experiments (at least 6 embryos imaged per genotype).

### A cross-evolutionary survey of eukaryotic ribosomal RNA acetylation machinery

To more fully apprehend the evolutionary origin of SNORD-mediated rRNA acetylation, we revisited the phylogenetic distribution of SNORD13 throughout eukaryotes, focusing primarily on Metazoan (animals) and Archaeplastida (organisms capable of photosynthesis). Using BLAST searches with low stringency algorithm parameters together with manual inspection, we identified many novel candidate SNORD13 homologs outside the vertebrate lineage, including Urochordata, Cephalochordata, Echinodermata, Arthropoda, Mollusca, Brachiopoda, Cnidaria and even Porifera (**Figure 4A/B**). As depicted in **Figure 4B**, many of the newly-identified SNORD13 genes are located within introns, both in the sense or antisense orientation with respect to transcription of their host-gene transcript. In Archaeplastida, SNORD13 was also uncovered in phylogenetically distant land plants, including in the freshwater green alga *C. braunii* (**Figure 4C**). A full list of newly-identified SNORD13 and proposed snoRNA:rRNA duplexes is provided in **Supplementary Data 1** and **Supplementary Figure 1,** respectively. These sequence analyses were accompanied by experimental profiling of ac^4^C at helix 45 in representative eukaryotic species. As listed in **Figure 4A**, acetylated cytidine was readily detected in the vast majority of organisms we examined, including unicellular eukaryotes such as slime molds (*P. polycephalum*), ciliates *(P. tetraurelia*, *P. falciparum*) or microalgaes (*P. tricornutum*). We found, therefore, that SNORD13 is much more broadly distributed than initially thought and SNORD-mediated RNA acetylation is a widespread principle in eukaryotes. Consistent with this conclusion, NAT10 homologs were found in almost all organisms we queried, with the notable exception of *E. gracilis* **(Figure 4A, Supplementary data 2)**.

**Figure 4.**
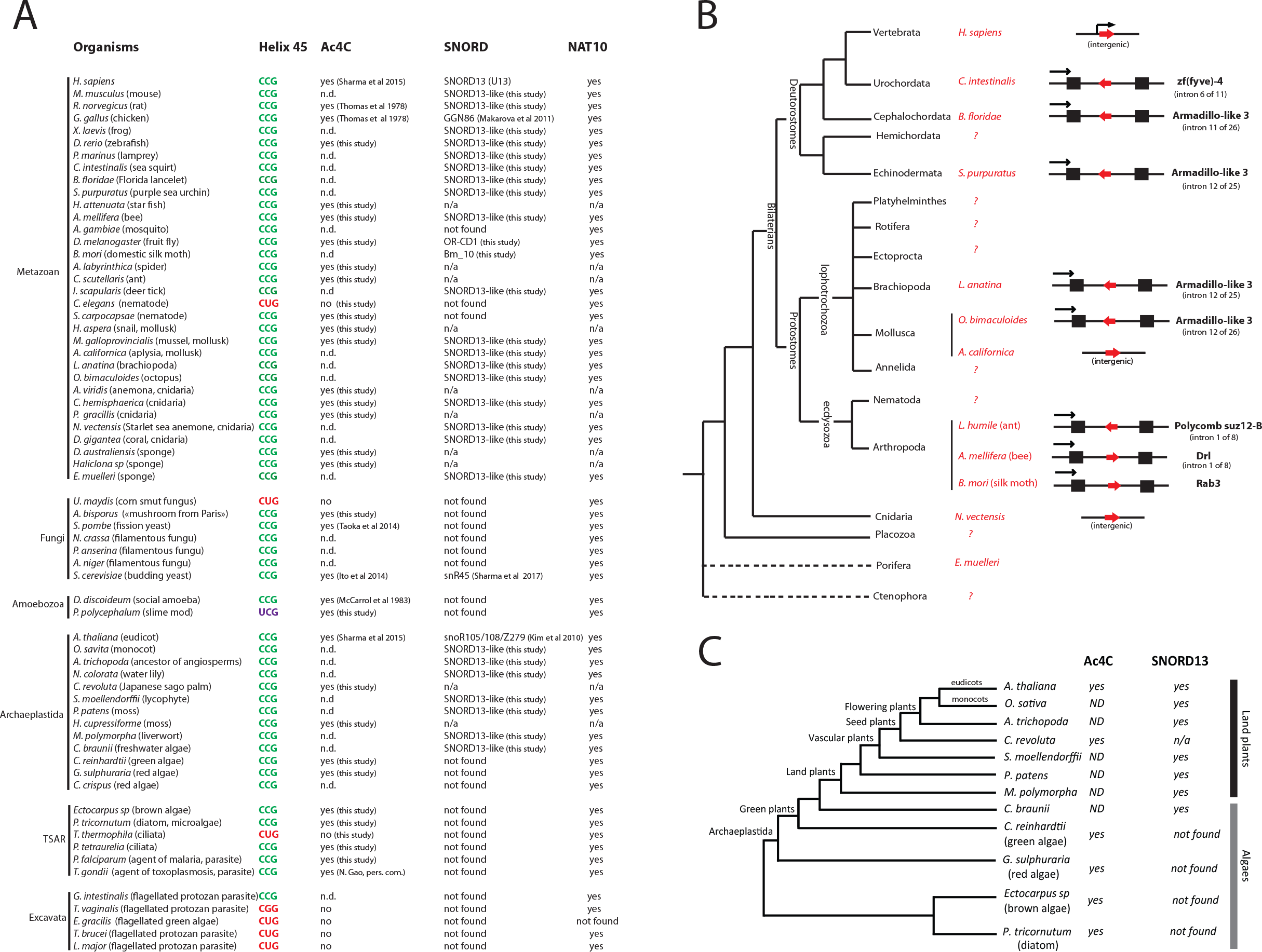
Cross-evolutionary survey of eukaryotic rRNA acetylation machinery. **A)** This table summarizes the conservation of the 5’-CCG-3’ motif, the acetylation status of helix 45 and the detection of SNORD13 and NAT10 genes in commonly used models or organisms for which we experimentally assayed the presence of ac^4^C at helix 45. n.d.: not determined. n/a: not applicable (genome is not available). **B)** Simplified phylogenetic tree of Metazoan. The genomic organization (intronic vs intergenic) of newly-identified SNORD13 genes (red arrows) is shown. Note that intronic SNORD13 can be positioned in the sense or antisense orientation with respect to their host-genes (black arrow). **C)** Simplified phylogenetic tree of Archaeplastida. Acetylation status of helix 45 and the presence of SNORD13 in the genome of representative species are indicated. Note that while rRNA in algal species is acetylated, we failed to identify obvious SNORD13 counterparts. A full list of newly-identified SNORD13 sequences can be found in **Supplementary data S1**.

### Atypical SNORD13-like homologs in Diptera

SNORD13-like genes are present in many groups of Arthropoda (**Figure 5A, Supplementary data 1**), including *D. melanogaster* and *B. mori* where they had previously been identified as SNORDs of unknown functions (named Or-CD1 and Bm-10, respectively; (24, 25)). In Diptera for which a genome annotation is available, Or-CD1 can be found either in large introns positioned in the 5’-UTR of two transcript isoforms generated from two alternative transcription sites (e.g. Swim, Tinagl1_2), or within short introns embedded in the open reading frame of the host-gene (e.g. Zfp15, Rps11) (**Figure 5B)**. To our surprise, the published sequence of Or-CD1, as well as that of all newly-identified SNORD13-like in Diptera, lack conserved antisense rRNA sequence at their 5’-end (**Figure 5C**). One potential explanation would be that the sequence upstream of the C-box is removed by exonucleolytic trimming of the spliced-out host-intron, very likely due to the absence of a 5’-capped structure which normally plays a protective role against degradation (26). Although plausible for small introns, such as those in Zfp15 or Rps11 genes, this hypothesis cannot account for all observations. For example, many Or-CD1-like in Arthropoda are located in introns (**Figure 4B**), yet they still retain a conserved rRNA complementarity in their 5’-end segment **(Figure 5C, Supplementary Figure 1)**. As an alternative, we reasoned that being intronic may not necessarily rule out independent transcription from distinct promoter regions. This is supported by the fact that SNORD13 can be found positioned either in the sense or antisense orientation relative to transcription of their host-gene (**Figure 4B/5B),** a scenario that is not compatible with post-transcriptional processing from introns.

**Figure 5.**
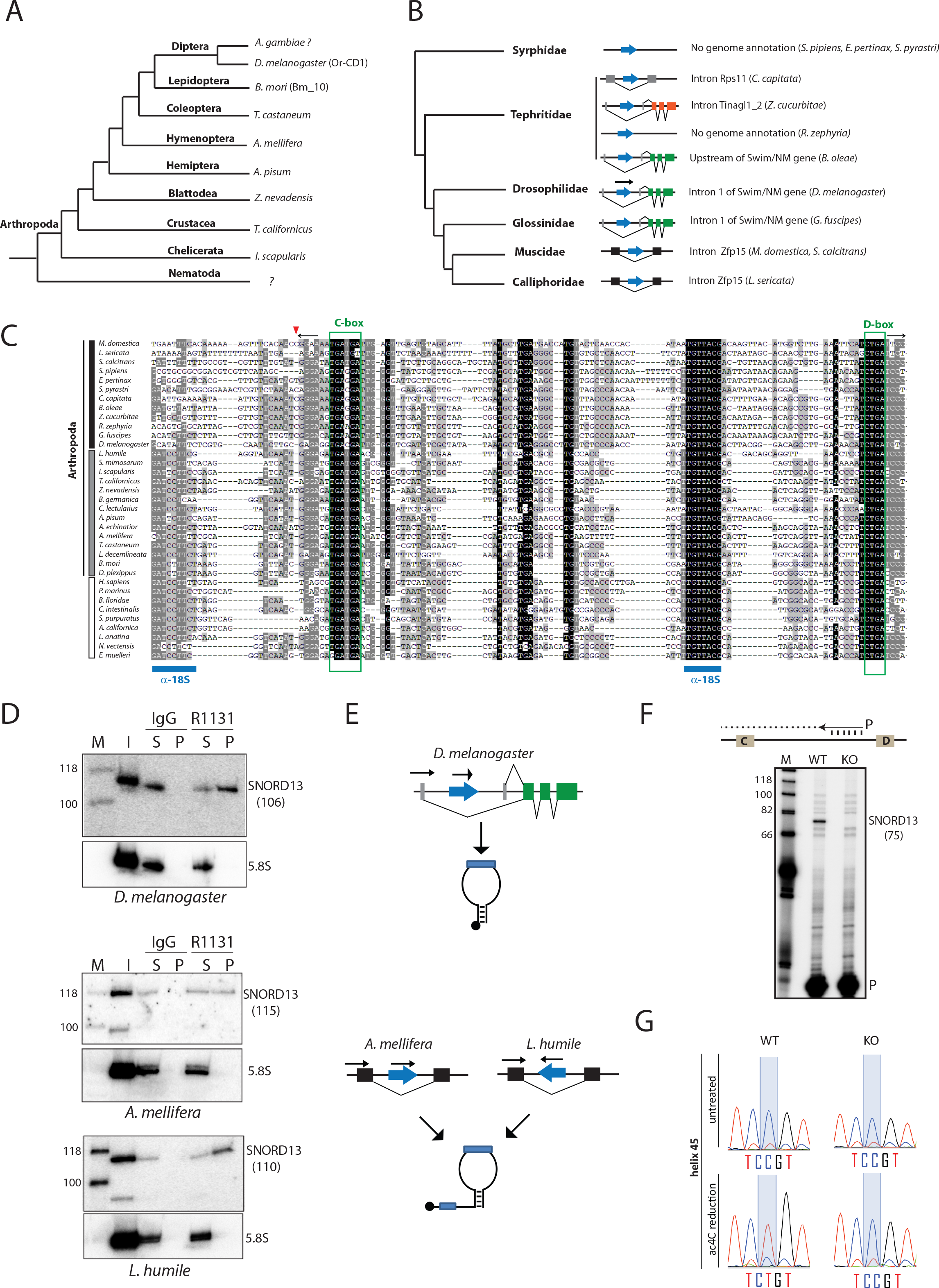
Atypical SNORD13-related RNA guides rRNA acetylation in *D. melanogaster*. **A)** Simplified phylogenetic tree of Arthropoda. Representative species where SNORD13-like were identified are indicated. **B)** Simplified phylogenetic tree of Diptera. The genomic arrangements of newly-identified SNORD13 genes (blue arrow) are shown. **C)** Multiple sequence alignment of representative eukaryotic SNORD13 sequences. The small red arrow indicates the relative position of the mature 5’-end of Or-CD1 in Diptera (see also panel F) while the two horizontal black arrows in opposite orientation depict the terminal 5’-3’ stem structure. Note that SNORD13 sequences in Diptera - but not in other species - lack one of the two conserved antisense rRNA elements (denoted by horizontal blue bars). Vertical black, grey and white bars highlight Diptera, other Arthropoda and non-Arthropoda species, respectively. **D)** Immunoprecipitation by anti-trimethyl cap (R1131) antibodies. OR-CD1 in various insects (as indicated above the panels) was detected by Northern blots. The same membrane was hybridized with an antisense oligo probes that recognized uncapped 5.8S rRNA used as negative controls. Input RNA (I); RNA recovered from the pellet (P), RNA recovered from the supernatant (S). The theoretical size of SNORD13 (nt) in each of the species studied is indicated in parentheses. **E)** Schematic representation of two distinct modes of expression: independent transcription gives rise to short (top) or long (bottom) forms of 5’ capped SNORD13. **F)** A mutant KO fly strain harboring an inserted P-element in the Or-CD1 gene (Dmel\P{EP}snoRNA:Or-CD1G9117) does not express OR-CD1 as assayed by primer extension. Note that the 5’-end of the cDNA product confirms that fly Or-CD1 lacks one 18S rRNA complementarity. The expected size (nt) of OR-CD1 with ∼ 4-5 nucleotides upstream of the C-box (as shown on the panel) is indicated in parentheses. ^32^P labeled primer (P). **G)** Misincorporation (C-to-U) at SSU-C1968 (helix 45) as judged by Sanger DNA sequencing of RT-PCR products obtained after borohydride treatments of total RNA extracted from WT and Or-CD1-KO adult flies.

To probe the mode of expression of Or-CD1 in Arthropoda, total RNAs extracted from *D. melanogaster*, *L. humile* (ants) and *A. mellifera* (bees) were subjected to immunoprecipitation with R1131 antibodies that recognize the 2,2,7 trimethylguanosine (TMG) cap structure, enabling its use as a proxy for independent transcription. Consistent with the expected size for small RNAs lacking 5’-end extension in *D. melanogaster*, Northern blot analyses detected a radioactive signal migrating at about 110 nucleotides **(Figure 5D, see also Figure 5F)**. These RNA species were specifically immunoprecipitated with the R1131 antibodies indicating that, despite being located within an intron, Or-CD1 gene is independently transcribed. This conclusion is in line with prior studies showing that Or-CD1 represents one of the very few SNORD genes that recruit RNA polymerase II and the Little Elongation Complex (LEC;(27)). In agreement with phylogenetic comparison that identifies two conserved antisense rRNA elements (**Figure 5C**), Or-CD1 displays a 5’-capped extension in both *L. humile* and *A. mellifera* (**Figure 5D**), as evidenced by the sizes of RNAs recovered in the immunoprecipitation pellet fraction. This therefore identifies a second expression strategy by which independent transcription from introns can also yield “regular” extended forms of Or-CD1 **(Figure 5E)**.

Thus, Or-CD1 in Diptera - but not in Arthropoda - displays only one antisense rRNA sequence **(Figure 5C)**. In order to test if this remarkable fly SNORD13 was capable of guiding rRNA acetylation, a strain bearing a P-element inserted in the Or-CD1 gene was retrieved from the Bloomington Drosophila Stock Center (Dmel\PsnoRNA:Or-CD1G9117). After demonstrating that Or-CD1 was no longer detected in the mutant adult flies (**Figure 5F)**, we showed that acetylation in helix45 was abolished (**Figure 5G).** Thus, SSU-C1968 in *D. melanogaster* can be targeted for acetylation by a SNORD13 homolog displaying a single antisense element. Of note, Or-CD1 adult knockout flies are viable, fertile and do not manifest gross abnormalities (not shown), thus strengthening the notion that N4-acetylcytidine at helix 45 is largely dispensable for viability, at least under standard laboratory conditions. Based on our findings, we propose to rename Or-CD1 as SNORD13.

### Life without 18S rRNA acetylation: the cases of *C. elegans* and genetically modified ΔsnR4ΔsnR45 mutant yeast

A key role in ribosome biology of 18S rRNA helix 45, which in mature small ribosomal subunits lies directly adjacent to the decoding site, is implied by its extreme sequence conservation in many phylogenetically-distant organisms belonging to all known eukaryotic supergroups. Furthermore, when mutations in helix 45 arise, they are often accompanied by compensatory base changes (**Supplementary Figure 4A**). The 5’-CCG-3’ motif - wherein the second C is targeted for acetylation - was established as a key specificity sequence determinant for NAT10 (3, 6). Remarkably, a few well-studied organisms lack this conserved motif or even the targeted substrate cytidine, including flagellated protozoan parasites (*T. vaginalis, T. brucei, L. major*), protozoan algae (*E. gracilis*), ciliate protozoa (*T. thermophila*), slime mold (*P. polycephalum*), corn smut fungus (*U. maydis*) or even the nematode (*C. elegans*) (**Figure 4A).** Other examples can be found in **Supplementary Figure 4B**. Only in the slime mold *P. polycephalum* did our ac4C sequencing reaction provide evidence for helix 45 acetylation within a non-consensus 5’-UCG-3’ motif, suggesting that NAT10 may have a distinct biochemical specificity in this organism **(Supplementary Figure 5)**. Of note, helix 34 of *P. polycephalum* is apparently not acetylated despite the presence of a similar 5’-UCG-3’. Given its importance as animal model and the prior determination that it carries an essential NAT10 gene (named nath-10)(28), we next focused on *C. elegans*. As shown in **Figure 6A-6B**, helix 45 of *C. elegans* 18S rRNA harbors a 5’-CUG-3’ motif also found in other nematodes, notably those belonging to the Clades IV and V. By definition, this motif variant that lacks a middle cytidine cannot be acetylated canonically. Nonetheless, since NAT10 is present in the worm genome, we verified the absence of any other ac^4^C residue within helix 45 (**Figure 6C,** not shown). More surprisingly, we found that the 5’-CCG-3’ motif in helix 34 was also devoid of acetylation (**Figure 6C**). The lack of rRNA acetylation in *C. elegans* elsewhere than in helices 34 and 45 was further examined using anti-ac^4^C immuno-northern blotting, in which signals for acetylated RNA were found to be limited to tRNAs (**Figure 6D)**. It is important to note that ac^4^C-free ribosomes are not necessarily a universal feature of nematodes. Indeed, helix 45 was found to be acetylated in *S. carpocapsae,* an entomopathogenic nematode of the *Steinernematidae* family **(Supplementary Figure 5)**. The evolutionary forces that drove disappearance of rRNA acetylation in *C. elegans* remain unknown. Nonetheless, our results indicate that the changes in robustness of vulval cell-fate specification observed when the essential nath10 gene was mutated (28, 29) did not reflect loss of rRNA modification but more likely the involvement of nath10 in pre-rRNA processing. A similar conclusion was reached recently when it was shown that the involvement of Fibrillarin in neural crest cell maturation depends on its role in pre-rRNA processing rather than in methylation (doi.org/10.1101/2021.11.25.469989). This is similar to prior observations in *S. cerevisiae* where Kre33 was shown to be essential but snR45 and/or SSU-ac^4^C1773 were not (16). As further validation of ac^4^C’s dispensability in rRNA we interrogated a double mutant yeast strain in which snR4 and snR45 genes were simultaneously deleted, producing rRNA completely devoid of ac^4^C. As observed in **Figure 6E**, growth of double ΔsnR45ΔsnR4 mutants, as assayed in liquid or solid media, was undistinguishable from that of its wild-type counterpart whether at 30°C or higher sub-optimal temperatures. The lack of obvious phenotypes for ΔsnR4ΔsnR45 mutants at 30°C was also previously reported (29). Collectively, these data are in agreement with previous findings showing that ΔKre33 yeasts re-expressing a catalytic-dead form of Kre33 (H545A or R637A) grow normally, as compared to ΔKre33 yeasts re-expressing a WT form of Kre33 (16). Thus, we confirm that the absence of rRNA acetylation is unlikely to contribute to the essential function of Kre33. This latter lies very likely with its modification-independent requirement for pre-rRNA processing events leading to the production of mature 18S rRNA (5). This conclusion does not, however, rule out some roles for tRNA acetylation, particularly under suboptimal growth conditions. Indeed, tan1 mutant strains show reduced growth rates at higher temperature (39°C), presumably due to decreased stability of unmodified serine tRNAs (30).

**Figure 6.**
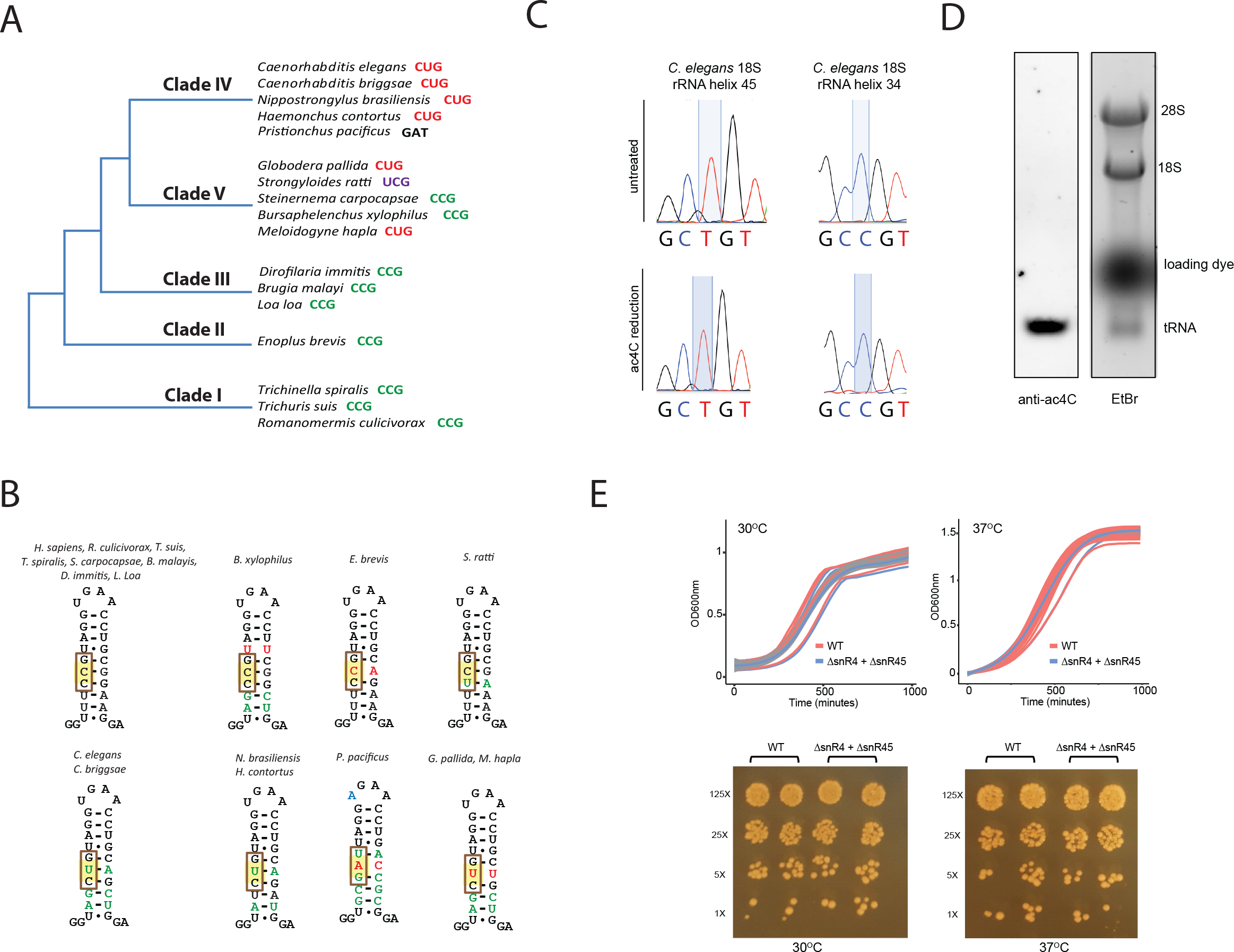
Life without rRNA acetylation. **A)** Simplified phylogenetic tree of Nematoda. Sequence contexts of the acetylated cytidine in representative species of each clade are indicated. **B)** Schematic representations of helix 45. Green bases represent compensatory base changes, as compared to human helix 45, while red bases provoke irregularity in the geometry of helix 45. The triplet sequence motif is framed. **C)** Sanger DNA sequencing of RT-PCR products obtained after borohydride treatments of total RNA extracted from *C. elegans*. **D)** Ac^4^C content in total RNA extracted from adult *C. elegans* was visualized by Northern blotting-based assay using anti-Ac^4^C antibodies. Ethidium bromide (etBr) staining indicates the relative position of rRNA and tRNA species. **E)** Top-panels: WT and ΔsnR4ΔsnR45 yeast strains were grown at 30°C (left) or 37°C (right) in liquid YPD. Optical density at 600 nm (OD_600_) was measured every 30 minutes. Growth curves of individual biological samples (thin lines; n=3 or 12 for 30 °C and 37 °C samples, respectively) and the mean of replicates (thick lines) are presented along with a confidence interval (gray area). Bottom-panels: WT and ΔsnR4ΔsnR45 yeast strains were spotted on a YPD-agar plate and incubated for 48 hours at 30°C or 37°C. Cells were grown in duplicates with a 5-fold dilution between distinct concentrations.

## Discussion

In order to interrogate the function of a single SNORD13-dependent cytidine acetylation in helix 45 of 18S rRNA, we have combined a cross-evolutionary survey of rRNA acetylation machinery across the eukaryotic tree of life and loss-of-function studies in four distinct models: budding yeasts, human cells, zebrafish and fly. This study represents one of the very few examples addressing the physiological roles of a SNORD in multicellular organisms bearing a germline-derived knockout (31–35). Although this peculiar rRNA ac^4^C is highly conserved, its loss surprisingly does not appear to grossly impact ribosome biogenesis or function, or even embryonic development. Consistent with this observation, we identify *C. elegans* as an evolutionary example of dispensable metazoan rRNA acetylation. Finally, the discovery of a 5’ truncated form of SNORD13 in *D. melanogaster* suggests that the mechanism by which SNORDs assist NAT10 in catalyzing rRNA ac^4^C is likely more diverse than previously anticipated and in some case may rely on a single antisense element.

An important question raised by our study concerns the specific role of ac^4^C in 18S rRNA helix 45. In humans, the ribosomal protein eL41 (formerly RpL41) is positioned at the interface between the 40S and 60S subunits (36, 37) and directly contacts ac^4^C1842 (38). This led to the proposal that acetylation may mediate functionally important crosstalk between the two subunits (38). However, our experimental settings failed to uncover any robust defects in ribosomal subunit assembly or protein synthesis in human SNORD13-KO cells. This is perhaps not surprising since eL41 knockout in yeast does not affect translation unless formation of another inter-subunit contact is compromised (39). A prominent role of ac^4^C in eL41 assembly and/or RNA::protein interaction is also quite unlikely since eL41 is correctly incorporated into ribosomes of *T. vaginalis* (40), *E. gracilis* (41), *L. donovani* (42) and *T. brucei* (42), all of which lack acetylation in helix 45 **(Supplementary Figure 6)**. Even more tellingly, the relative positioning of eL41 in ribosome and the 3-D architecture of helix 45 in the SSU of the archaea *T. kodakarensis* are not affected when TkNAT10 gene is deleted (6). Altogether, these observations point to a subtle molecular role for ac^4^C, possibly in fine tuning ribosome structure and/or function in sub-optimal environmental contexts and/or sensitized genetic backgrounds. Unravelling the precise physiological relevance of 18S rRNA acetylation in multicellular organisms, if any, represents a conceptually and technically formidable, but important, task for future studies.

Our bioinformatics search has considerably expanded the inventory of SNORD13 sequences in metazoans, identifying homologs well beyond vertebrates where they were previously thought to be restricted. However, we were unable to identify SNORD13 counterparts in many major phyla including Hemichordata, Annelida, Nematoda, Fungi (other than budding yeasts), algae or unicellular eukaryotes. Although the most parsimonious hypothesis would be that rapidly-evolving SNORD13 sequences make them difficult to detect, one cannot exclude the possibility that SNORD13 may have been lost in these lineages, which would imply that in these organisms rRNA acetylation may be catalyzed by NAT10 alone. More sophisticated structural and biochemical analyses will be critical to solve this question. An additional related question is whether acetylation of helix 34 is also strictly dependent on an antisense SNORD, as shown only in *S. cerevisiae* to date (16). The characterization of RNA-guided rRNA acetylation systems in Archaea, should they exist, represents an additional avenue that may lend invaluable evolutionary and mechanistic insights into this process (6, 7).

Single nucleotide resolution cross-linking analysis has previously described how SNORDs guide the acetyltransferase to its rRNA substrate by use of two short base-pairing interactions on each side of the substrate cytosine, forming a “modification pocket” akin to H/ACA snoRNAs (SNORA) acting in pseudouridylation (16). During the course of this study, we unexpectedly identified independently-transcribed SNORD13 gene in *D. melanogaster* that produces RNA species lacking 5’-sequence complementary to rRNA (**Figure 7**). The same may also hold true in other Diptera for which SNORD13 is embedded within a very short intron (e.g. *L. sericata, M. domestica*). In this case, it is indeed more than likely that exonucleolytic processing of the host-intron produces uncapped RNAs with the C- and D-boxes brought together by a short terminal 5’-3’ terminal stem, as commonly found for other intron-located SNORD (26). The genomic constraints and selection pressure that led to changes in SNORD13 expression strategies across evolution are unknown, yet they have inherent mechanistic repercussions since the deposition of ac^4^C in helix 45 in *D. melanogaster* does not strictly require two antisense elements flanking the rRNA ac^4^C site. This finding contrasts with conclusions drawn from mutagenesis of yeast snR45 where the 5’ antisense element appears essential for acetylation (16). It therefore implies an alternative mode of action of SNORD13 in *D. melanogaster* and, also more broadly, in Diptera. Finally, the overall structure of SNORD13 in Diptera is indistinguishable from that of the canonical intron-located SNORDs which mediate rRNA ribose methylation. It thus raises the intriguing question as to whether other intronic SNORDs in these organisms, particularly those of still elusive function, might also mediate acetylation on still unidentified RNA targets.

**Figure 7.**
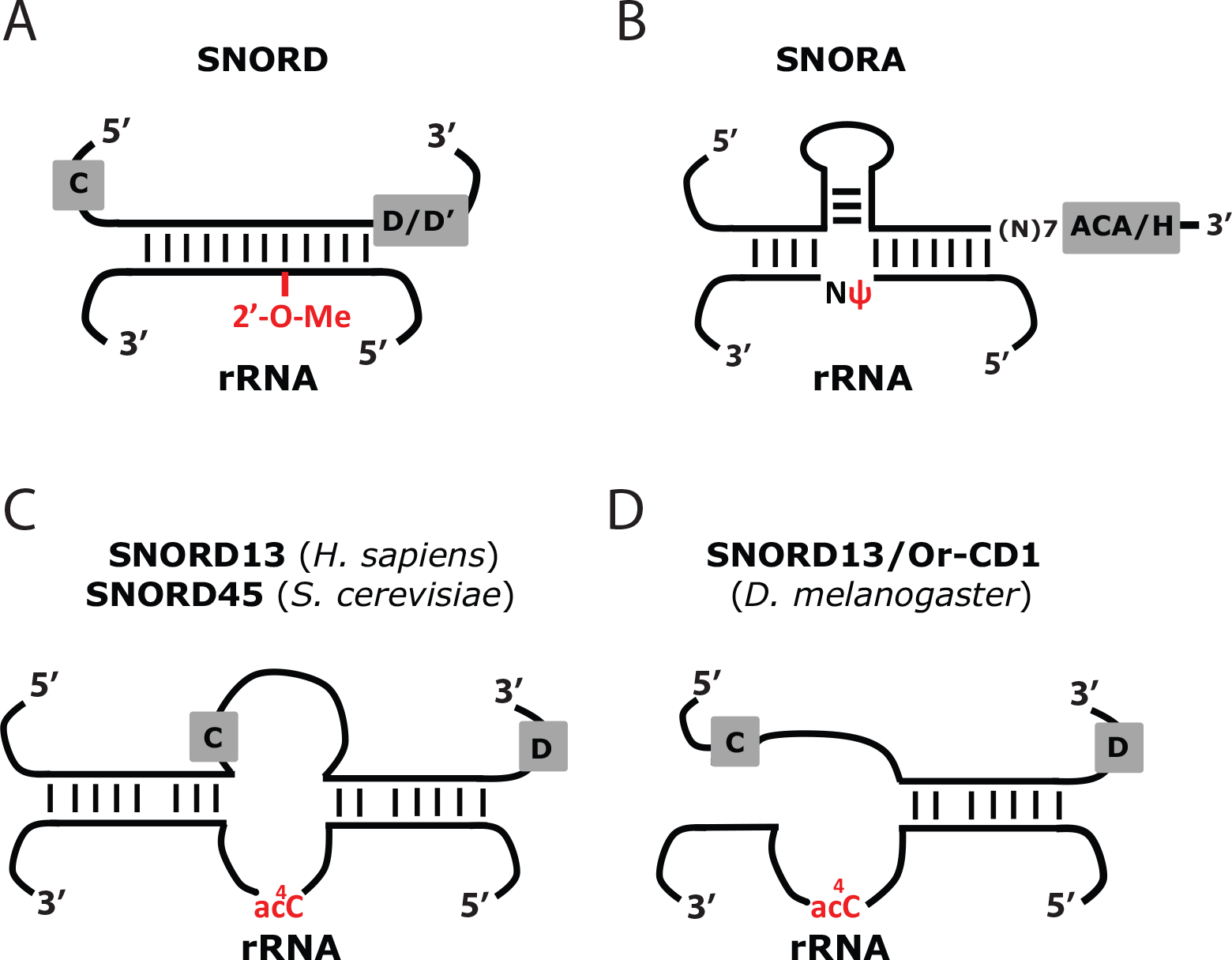
Comparative schematic representation of rRNA modification systems. **A)** Through the formation of a perfect (or near perfect) RNA duplex (∼10-20 bp), SNORDs guide Fibrillarin/Nop1 to their target rRNA nucleotides, i.e. the nucleotide to be 2’-O-methylated (2’-O-Me) is paired to the fifth position upstream of the conserved D (or D’) box. **B)** Through the formation of two perfect (or near perfect) RNA duplexes (∼4-10 bp each), SNORAs guide Dyskerin/Cbf5 to their target rRNA nucleotides, i.e. the uridine to be isomerized into pseudouridine (ψ) remains unpaired and it is located at ∼ 14-16 nt from the conserved H (or ACA) box. **C)** snR45, snR4 and SNORD13 contain two imperfect antisense elements matching either side of the cytidine to be acetylated (ac^4^C) and, through still unknown mechanisms, guide cytosine acetylation by NAT10/Kre33. **D)** In *D. melanogaster*, SNORD13 targets rRNA acetylation at helix 45 through the use of a single antisense rRNA sequence.

Altogether, our data point to interlaced questions regarding SNORD13 sequence diversity across evolution, the molecular mechanisms underlying site-specific rRNA acetylation and the physiological importance of ac^4^C in rRNA in cell homeostasis, development and physiology. In our view, tackling these complex issues through an evolutionary perspective will be undoubtedly fruitful and may produce more surprises since, paradoxically not to mention ironically, observations made here with two widely-studied models - *C. elegans* and *D. melanogaster -* depart considerably from those previously obtained from vertebrates and budding yeasts.

## Materials and Methods

Unless otherwise noted, all techniques for cloning and manipulating nucleic acids were performed according to standard protocols.

### Generation and characterization of SNORD13-deficient HAP1 cells

HAP1 cells were grown at 37°C with 5% CO_2_ in IMDM (Gibco; 4.5 g/L glucose) supplemented with 10% fetal bovine serum (PAN biotech), 1 mM Sodium Pyruvate (Gibco) and 1% penicillin-streptomycin (Sigma-Aldrich). The human SNORD13 gene was disrupted via CRISPR/Cas9-mediated deletion. Two sgRNAs selected using the CRISPR design tool at https://zlab.bio/guide-design-resources were cloned into pX459 V2.0 (Addgene #62988) plasmid. HAP1 cells seeded in six-well dishes (70-80% confluency) were co-transfected using lipofectamine 2000 (Thermo Fisher Scientific) with pX459_sgRNA1 and pX459_sgRNA2 (500 ng each). 48 hours post-transfection, cells were transiently treated with puromycin (2 µg/mL; ∼ 36 hours) before being subjected to clonal selection by limiting dilution. Deletion events were validated by PCR (primer sequences are listed in **supplementary data S3)** and three clones per genotype were randomly chosen for further analyses. The rate of cell growth was assayed using Beckman Coulter Z1 particle counter. Cells were seeded into a 24-well plate and counted in quadruplicate at each time points. Six independent proliferation assays were performed. Apoptosis was detected using Annexin V-FITC/propidium iodide staining according to the manufacturer’s instructions (BioLegend). Cells were analyzed using FacsVerse (Becton Dickinson) with acquisition of 20,000 total events (BD FacsSuite). Two independent experiments were performed.

### RNA extraction and rRNA analyses

Total RNA was prepared using TRI reagent (MRC) according to the manufacturer’s instructions and treated with RQ1 RNAse-free DNase (Promega) and proteinase K (Sigma). For Northern blot analyses of low molecular-weight species, total RNA (10 µg) was fractionated by electrophoresis on a 6% acrylamide, 7 M urea denaturing gel and electrotransferred onto a Hybond N membrane (Amersham GE Helthcare) as described previously (43). For Northern blot analyses of higher molecular-weight species, 4 µl of total RNA (2.5µg/µl) were added to 20 µl of “glyoxal mix” (60% DMSO, BTPE 1.2X, 5% glycerol, 8% glyoxal solution (Sigma) and 40 µg/ml Ethidium bromide), heat-denaturated at 55°C for 60 min, chilled on ice for 10 min and separated by electrophoresis on a 1.2% agarose gel (Pipes 10 mM, Bis-tris 300 mM, EDTA 10 mM). Capillarity transfer was performed onto Hybond N+ membrane. In all cases, transfer was followed by UV light irradiation and membranes were pre-hybridized for 1 hour in 5x SSC, 0.1% SDS, 5x Denhardt’s, 150 mg/ml yeast tRNA. Hybridization was carried out overnight at 50°C with 5’-[^32^P]-labeled-DNA oligonucleotide probes (100 000 cpm/ml). Membranes were washed twice with 0.1XSSC, 0.1% SDS at room temperature (2x10 min) before autoradiography. Pulse chase experiments were performed as described in (44). The day before, 12-well plates are seeded with 500,000 cells. Cells are then incubated in methionine-free DMEM (Invitrogen) for 30 min at 37°C, labeled for exactly 15 min with L-methyl ^3^H methionine (50mCi/ml) and rinsed twice with IMDM containing unlabeled methionine. They were then incubated for 0 min (rinsed immediately twice with cold PBS), 30 min, 1h, 1h30, 2h, 2h30 in regular IMDM. After the corresponding chase time, cells were rinsed twice with cold PBS and lysed with Tri-Reagent (MRC). RNA samples were glyoxal denatured and separated on a 1.2% agarose gel and passively transferred to a nylon membrane as described above. The membrane was exposed 5 weeks to Biomax KODAK MS films with a KODAK BioMax Transcreen LE. RNAse H mapping was performed as described in (44). A water solution containing 4 µg of total RNA was heat-denatured (95°C for 3 min) in the presence of an antisense oligonucleotide matching the 3’-end of 18S rRNA (RNase H_2, 100µM). After allowing annealing by cooling down to room temperature, the RNA/oligonucleotide mixture was diluted to 30 µl final with a reaction mix (1X RNase H reaction buffer, 65 µM DTT, 0.5U/µl RNasin (Promega), 50 U RNase H (New England Biolabs) and incubated at 37°C for 30 min. The reaction was stopped by adding 0.3 M sodium acetate pH 5.2/0.2 mM EDTA. RNA was phenol extracted, ethanol precipitated, loaded onto a 15% acrylamide gel and analyzed by Northern blot (^32^P-labeled 3’18S oligo-probe) as described above. For primer extension, two µg of total RNA extracted from adult flies were mixed with a 5′-^32^P-labeled oligo (∼100000 cpm) and cDNA synthesis, generated according to the manufacturer’s instructions, were resolved onto a 12 % denaturing acrylamide gel.

### Sucrose gradient sedimentation and Sunset assay

Exponentially growing WT and SNORD13-Ko cells (∼ 20 million per genotype) were treated with 100 µg/ml of cycloheximide (CHX, SIGMA) for 10 min at 37°C, rinsed twice with cold PBS/CHX (100µg/ml), collected by centrifugation (400 g, 5 min) and resuspended in 5 ml of cold Buffer A (10 mM Hepes, 1,5 mM MgCl2, 100mM KCl, CHX 100µg/ml, 1mM DTT). Cells were then centrifuged (400g, 10 min at 4°C), suspended in 700 µl of Buffer A and incubated 10 min on ice. Cells were then mechanically disrupted with a cold Dounce homogenizer (B) and centrifuged for 10 min (1000 g at 4°C). The cytoplasmic fraction (supernatant) was collected and centrifuged twice at 10 000 g (10 min, 4°C). About 1,5 mg of protein was loaded on a 10–50% sucrose gradient and centrifuged for 2h45min at 39 000 rpm in an Optima L-100XP Ultracentrifuge (Beckman-Coulter) using a SW41 rotor (Beckman). Gradients were then monitored at 260 nm and fractions were collected from the top using a Foxy Jr. fraction collector (Teledyne ISCO). Sunset assay was performed as described in (45). Exponentially growing cells were treated with 1 µg/ml of puromycin for 20 min. As a control, cycloheximide (100 µg/ml) was also added 15 min before puromycin treatment. Total cell extracts were then processed for western blot using an anti-puromycine antibody (Millipore, clone 12D10 #MABE343) as well as Anti-CDK9 (C12F7) mAb (cellsignal.com) as gel loading control.

### Immunoprecipitations

Thirty microliters of rabbit R1131 anti-trimethyguanosine (3 mg/ml, a gift of Dr. R. Luhrmann) were incubated with gentle agitation for 120 min. at 4°C with 80µl of Proteine A Sepharose® 4B, Fast Flow from *S. aureus* (PAS, Sigma #P9424) in 1 ml of NET-150 buffer (50 mM Tris-HCl (pH 7.4), 150 mM NaCl, 0.05% Igepal® CA-630, SIGMA). Mouse IgG (Jackson Immunoresearch #315-005-008) was used as negative controls. PAS-Ig pellets were washed three times with 1 ml of NET-150 buffer. Thirty micrograms of total RNAs was then added to PAS-R1131 (or PAS-IgG) in 0.5 ml of NET-150 and incubated with gentle agitation for 60 min. at 4 °C. Pellets were then collected by centrifugation and washed seven times in 1 ml of NET-150. RNAs from pellet and supernatant were extracted by SDS/phenol extraction and analyzed by Northern blot as described below.

### Searching for SNORD13-like and 18S rRNA sequences in eukaryotic genomes

Using vertebrate SNORD13 sequences as queries, iterative BLAST searches with low stringency algorithm parameters, together with manual inspection of some hits, were conducted using eukaryotic genome databases found at https://blast.ncbi.nlm.nih.gov/Blast.cgi, https://rnacentral.org/, https://www.echinobase.org/entry/, http://mgbase.qnlm.ac/home, http://www.insect-genome.com/, https://bipaa.genouest.org/is/, https://i5k.nal.usda.gov/ webapp/blast/, https://spaces.facsci.ualberta.ca/ephybase/. Eukaryotic 18S rRNA sequences were retrieved either at https://www.arb-silva.de/ or https://www.ncbi.nlm.nih.gov/nucleotide/.

### Mung bean nuclease protection assay and RP-HPLC

Mung bean nuclease (MBN) protection assay was performed exactly as described before (46). 500 pmol of the synthetic deoxy oligonucleotide (5’-TAATGATCCTTCCGCAGGTTCACCTACGGAAACCTTGTTACGAC TTTTAC*-3’*) was incubated with 50 pmol of 18S rRNA and 5% of DMSO in 0.3 volume of hybridization buffer (250 mM of HEPES, 500 mM of KCl at pH 7). The mixture was incubated at 90°C for 5 minutes and then slowly cooled down to 45°C over 2 h. After hybridization, 35 units of mung bean nuclease (New England Bio Labs (NEB)) and 0.02 mg/ml RNase A (Sigma-Aldrich) along with appropriate amount of 10x MBN buffer (NEB) were added to start digestion. The digestion was carried out at 35°C for 1 h. The protected fragment (RNA-DNA hybrid) was extracted from the reaction mixture by phenol/chloroform extraction, followed by overnight ethanol precipitation. The protected rRNA fragment was separated from the complementary DNA oligonucleotides on a denaturing 7 M Urea 15% PAGE gel. Bands were visualized by ethidium bromide staining and the rRNA band was excised and eluted using the D-Tube TM Dialyzers according to the manufacturer’s protocol for electro elution (Novagen). Eluted rRNA fragment was digested to nucleosides using P1 nuclease and alkaline phosphatase as described before (46). Nucleosides were next analyzed by RP-HPLC on a Supelcosil LC-18-S HPLC column (25 cm x 4.6 mm, 5 μm) equipped with a pre-column (4.6 × 20 mm) at 30°C on an Agilent 1200 HPLC system, using a protocol described previously in (16).

### Targeted ac4C sequencing

Nucleotide resolution sequencing of ac4C in 18S rRNA (helix 45 and helix 34) was performed as previously described (6). RNA samples were treated with sodium cyanoborohydride (100 mM in H_2_O) or vehicle (H_2_O) in a final reaction volume of 100 μL. Reactions were initiated by addition of 1 M HCl to a final concentration of 100 mM and incubated for 20 minutes at room temperature. Reactions were stopped by neutralizing the pH by the addition of 30 μL 1 M Tris-HCl pH 8.0. After incubation, reactions were adjusted to 200 μL with H_2_O, ethanol precipitated, desalted with 70% ice-cold ethanol, briefly dried on Speedvac, resuspended in H_2_O, and quantified by using a Nanodrop 2000 spectrophotometer. Cellular total RNA from individual reactions (200 pg) was incubated with specific h45/h34 rev primer (4.0 pmol) in a final volume of 20 μL. Individual reactions were heated to 65 °C for 5 min and transferred to ice for 3 min to facilitate annealing in SuperScript III reaction buffer (Invitrogen). After annealing, reverse transcriptions were performed by adding 5 mM DTT, 200 units SuperScript III RT, 500 μM dNTPs (use 250 μM dGTP) and incubating for 60 min at 55 °C.

Reaction was quenched by increasing the temperature to 70 °C for 15 min and store at 4 °C. cDNA (2 μL) was used as template in 50 μL PCR reaction with Phusion Hot start flex (New England Biolabs). Reaction conditions: 1X supplied HF buffer, 2.5 pmole each forward and reverse primer, 200 μM each dNTPs, 2 units Phusion hot start enzyme, 2 μL cDNA template. PCR products were run on a 2% agarose gel, stained with SYBR safe and visualized on UV transilluminator at 302 nm. Bands of the desired size were excised from the gel and DNA extracted using QIA-quick gel extraction kit from Qiagen and submitted for Sanger sequencing (GeneWiz) using the forward PCR primer. Processed sequencing traces were viewed using 4Peaks software. Peak height for each base was measured and the percent misincorporation was determined using the equation: “Percent misincorporation = (peak intensity of T)/(sum of C and T base peaks)*100%”.

### Dual luciferase reporter assay

100,000 WT and SNORD13-KO cells were seeded into a 24-well plate for 24 hours. Cells were transfected using lipofectamine 2000 (Invitrogen) using 0.06 ng Renilla vector (internal control) and 500 ng of reporter Firefly vectors. 24 hours post-transfection, cells were harvested, lysed and luciferase activity was monitored using the Dual-Luciferase Reporter Assay kit (Promega) according to the manufacturer’s recommendations. Luciferase detection was assayed on Centro LB 960 Microplate Luminometer (Mikrowin 2000 Software). Five-Six independent transfections were performed with each measurement made in triplicate.

### Yeast cells and media conditions

*S.cerevisiae* cells deleted of snR4 and snR45 or wild-type cells were grown at 30°C in standard YEP medium (1% yeast extract, 2% Bacto Peptone) supplemented with 2% dextrose (YPD). For Liquid growth assay, yeast cells were inoculated from a fresh YPD-agar plate into liquid YPD and grown for 48-72 hours at 30°C or 37°C. Optical density at 600 nm (OD_600_) of cells was then measured and used to dilute cells into a 96 well plate (Cat # 167008, Thermo Scientific) at equal ODs of ∼0.02OD (n=3 or 12 for 30°C and 37°C samples, respectively). Cells were then grown for 90 hours in an Epoch 2 microplate spectrophotometer (BID EPOCH2 Microplate Spectrometer from BioTek) at 30°C and 37°C and OD was measured every 30 minutes. Measured OD values were analyzed in R^2^. For presentation of growth curves, a regression line representing the mean of biological replicates was calculated using a generalized additive model. For Spot assay, yeast cells were inoculated from a fresh YPD-agar plate into liquid YPD and grown for 72 hours at 30°C and 37°C. Optical density at 600 nm (OD_600_) was subsequently measured and cells were serially diluted in 2% YPD, such that OD of cells varied from 0.02 OD to 0.00016 OD (5 fold dilutions). 3 µl of each diluted sample was used to spot cells on a YPD-agar plate, which was transferred to 30°C and 37°C. After 48 hours of incubation pictures were taken of the colonies. Each strain was grown in duplicates.

### Ethics statement and Zebrafish care

Fish were handled in a facility certified by the French Ministry of Agriculture (approval number A3155510). The project has received an agreement number APAFIS#7124-20161 00517263944 v3. Anesthesia and euthanasia procedures were performed in Tricaine Methanesulfonate (MS222) solutions as recommended for zebrafish (0.16 mg/ml for anesthesia, 0.30 mg/ml for euthanasia). All efforts were made to minimize the number of animals used and their suffering, in accordance with the guidelines from the European directive on the protection of animals used for scientific purposes (2010/63/UE) and the guiding principles from the French Decret 2013–118. Embryos were raised and staged according to standard protocols and the Recommended Guidelines for Zebrafish Husbandry Conditions (47, 48)

### Generation of Zebrafish snord13 mutants

The two guide RNAs (gRNA) were designed using CHOPCHOP CRISPR Design *website* (http://chopchop.cbu.uib.no). The designed oligos were annealed and ligated into the gRNA plasmid pX459 digested by *BbsI* (Thermo Scientific). The gRNAs were generated using the MEGAshortscript T7 transcription kit (Ambion) with PCR fragments containing T7 promoter. Transcripts were purified by phenol-chloroform extraction and Ethanol precipitation. 1 nl of a solution containing 10µM EnGen Cas9 NLS (NEB) and 100 ng/µl of gRNAs was injected at the one-cell stage. WT, heterozygous, and homozygous SNORD13 animals were then identified by PCR. Sequence primers can be found in **(supplementary data S3)**.

### RNA extraction, Reverse transcription and real-time PCR

Total RNAs from 15 wild type and 15 mutant zebrafish embryos were extracted using TRI reagent (MRC) and treated with RQ1-RNase free DNase I (Promega) and Proteinase K (Sigma) as recommended by manufacturer’s instructions. Total RNAs were converted into cDNA using GoScript RTase (Promega) random hexamer primers for 60 min at 42 °C according to manufacturer’s instructions. cDNAs were then diluted 20-fold and quantified by qPCR using SYBER green (Bio-rad) and specific primers. Data were acquired on CFX96 Real-Time PCR detection System (Bio-rad). Samples were analysed in triplicates and the expression level was calculated relative to zebrafish housekeeping gene *EF1α*. Sequence primers can be found in **(supplementary data S3)**.

### Immunostaining and *in situ* hybridization

Embryos were fixed overnight at 4°C in BT-FIX, after which they were immediately processed or dehydrated and stored at −20°C until use. After fixation or rehydration, embryos were washed twice with Phosphate Buffered Saline/1% Triton X-100 (PBST), permeabilized with PBST/0.5% Trypsin for 30 sec and washed twice again with PBST. After blocking with PBST/10% Fetal Calf Serum (FCS)/1% bovine serum albumin (BSA) (hereafter termed ‘blocking solution’) for at least 1 h, embryos were incubated with antibodies directed against either cleaved Caspase-3 (Asp175) (Cell Signaling Technology), or HuC/D (Molecular Probes), in blocking solution overnight at 4 °C followed by 5 washing steps with PBST. Embryos were then incubated with the appropriate Alexa Fluor-conjugated secondary antibodies (Molecular Probes) for at least 2 h at room temperature and washed three times. Fluorescent in situ hybridization was carried out as previously described (49). *fli1a* riboprobe preparation has been described elsewhere (50). Embryos were dissected, flat-mounted in glycerol and images were recorded on the confocal microscope TCS SP8 (Leica Microsystems) with an L 25 × /0.95 W FLUOSTAR VIZIR objective (zoom X1.25) using the scanner resonant mode.

### Comparison of Ribosome 3D structures

Atomic models of ribosomes from the following species: *H. sapiens*, PDB: 6EK0 (Natchiar et al, 2017); *P. falciparum*, pdb: 6OKK and 3J79 (51) ;*T. gondii*, PDB: 5XXU (40); *T. vaginalis*, PDB: 5XYI (40), *L. donovani*, PDB: 6AZ1, 6AZ3 (42); *E. gracilis*, PDB: 6ZJ3 (41) were displayed and aligned in UCSF Chimera (52) using the “Matchmaker” option, with the human 18S rRNA chain as reference. Prior to this structural alignment, cryo-EM maps of *P. falciparum* and *T. gondii* (EMD-2660 and EMD-6780, respectively) were visually inspected. Using UCSF Chimera and Coot (53), unattributed densities above C2065 in *P. falciparum* and C1764 in *T. gondii* cryo-EM map allowed to unambiguously replace unmodified residues by their ac^4^C counterparts in the corresponding atomic models.

### Data availability statement

The data that support the findings of this study are available from the corresponding author [JC] upon reasonable request.

## Acknowledgments

We thank the many colleagues who kindly provided us with biological samples, particularly EMBRC-France (N. Turque, R. Lasbleiz), the Botanical garden of Toulouse, Insectosphere, Botanic-Blagnac, A. Mattout, S.Tournier, I. Massou, R. Jeanson, L. Distefano, P.Valenti, A. Dussutour, M. Betermier, J. Santos, K. Mochizuki, G. Allorent and P.M. Delaux. We are also grateful to Aurore Laire for excellent zebrafish care. This work was supported by Agence Nationale de la Recherche (ANR-18-CE12-0008-01). Research in the lab of JLM was supported by the Intramural Research Program of the NIH, National Cancer Institute, Center for Cancer Research (ZIA-BC011488). Research in the lab of D.L.J.L. is supported by the Belgian Fonds de la Recherche Scientifique (F.R.S./FNRS), the Université Libre de Bruxelles (ULB), the European Joint Programme on Rare Diseases (EJP-RD ‘RiboEurope’ and ‘DBAGencure’), the Région Wallonne (SPW EER ‘RIBOcancer’), the Internationale Brachet Stiftüng, and the Epitran COST action (CA16120).

---

**Supplementary figure S1.**
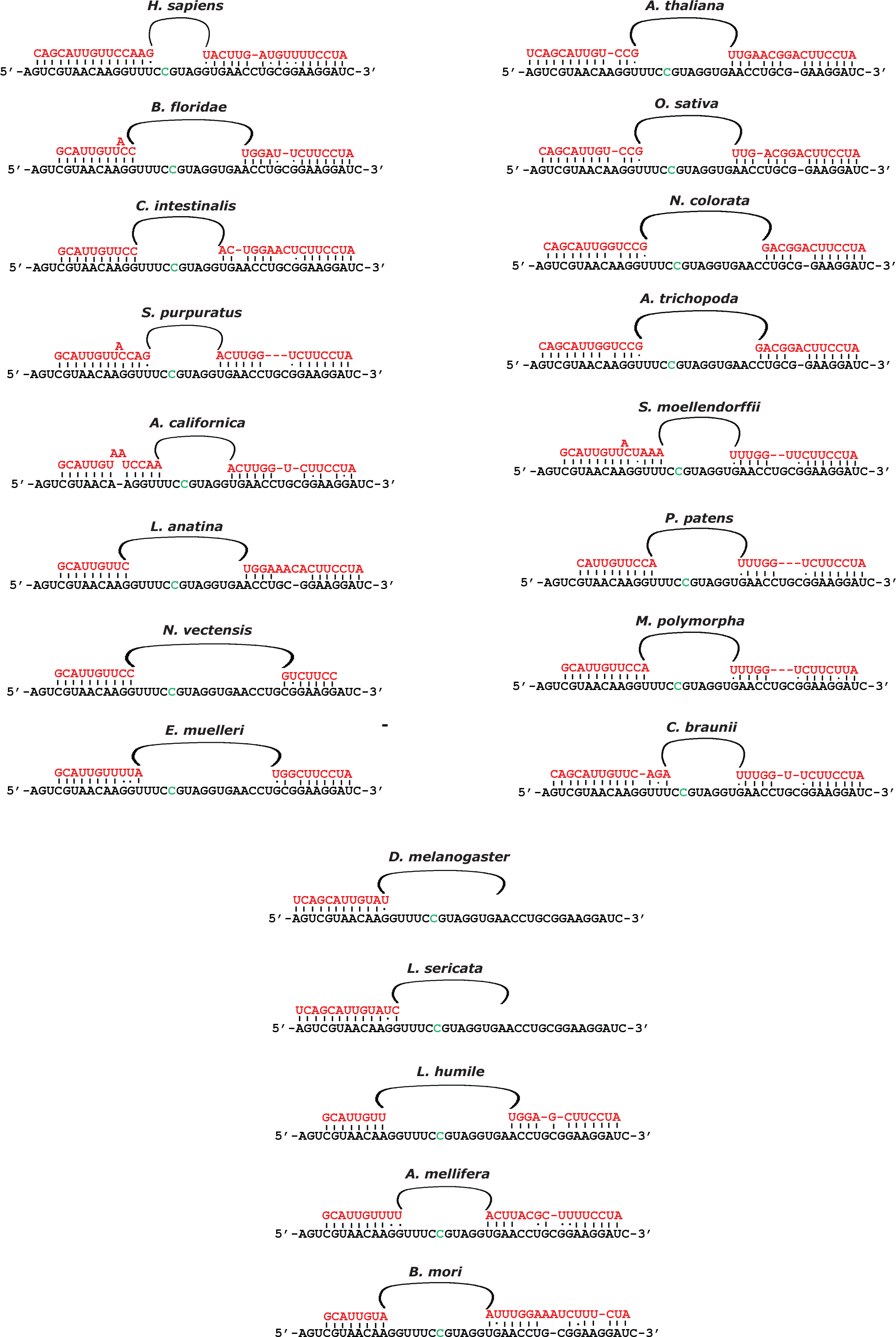
Examples of SNORD13-rRNA duplexes across evolution. Schematic representations of base-pairing interactions between SNORD13 and 18S rRNA (only a few representative examples are shown). The targeted C for acetylation is written in green.

**Supplementary figure S2.**
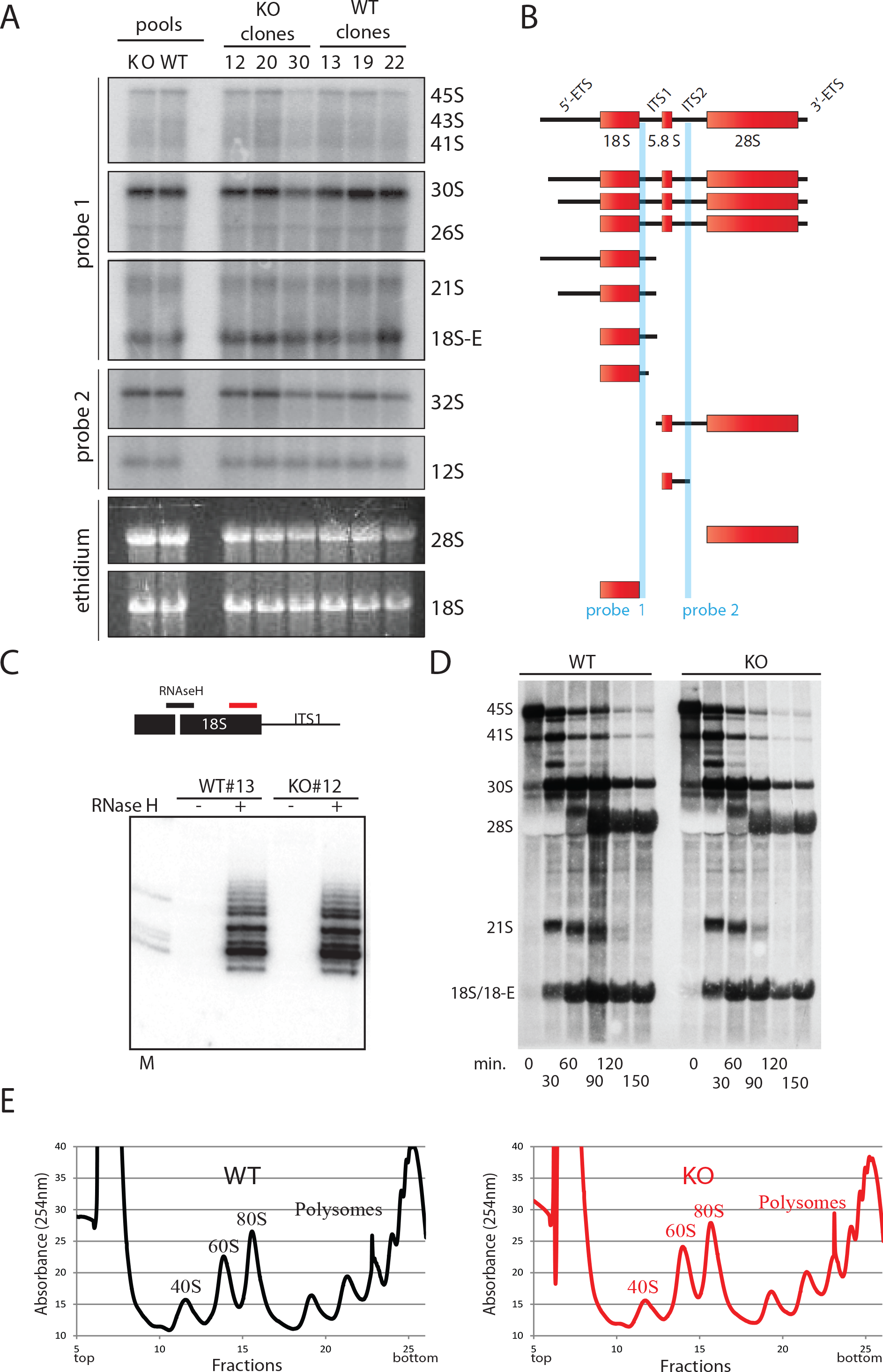
Loss of SNORD13 does not affect pre-rRNA synthesis and processing. **A-B)** Northern blot - Total RNA prepared from WT and SNORD13-KO clones, as well as from a pool of the three WT and KO clones, were analysed by Northern blot using probe 1 (5-ITS1) and probe 2 (ITS2) that detect main rRNA precursors intermediates as depicted in panel B). Cytoplasmic 18S and 28S rRNAs were also visualized by ethidium bromide staining of a denaturing 1.2% agarose gel. Relative quantification of rRNA intermediates based on three independent Northern blots did not reveal any obvious abnormalities (not shown). **C)** RNAse H assay - Total RNA extracted from WT or SNORD13-KO cells were annealed with antisense DNA oligonucleotide (black line), RNAse H digestion products were then resolved onto a 12% acrylamide gels and visualized by Northern blot with an antisense oligo-probe matching the 3’- end of 18S (red line). The ladder-like pattern reflecting heterogeneity of the 3’-end at 18S in SNORD13-KO cells was indistinguishable from that of observed in WT cells, indicating that the accuracy of cleavages that delineates the 3’ extremity of 18S rRNA is not impaired in SNORD13-KO cells. **D)** Pulse chase experiments - WT and SNORD13-KO cells were pulse-labelled with L-[methyl-^3^H]-methionine and harvested by chase times as indicated above the panel. Freshly-synthesized rRNA intermediates were detected by autoradiography after being separated by electrophoresis on denaturing 1.2% gel and transferred to a nylon membrane. Dynamics of rRNA synthesis in the SNORD13-KO cells was in the normal range, as shown by the timing of appearance and lifespan of the detected methyl-^3^H labelled rRNA precursors. **E)** Polysome profiling - Whole cytoplasmic extracts prepared from WT and (left) SNORD13-KO (right) cells were fractioned by ultracentrifugation on a 10-50% sucrose gradient. Absorbance (254 nm) of each fraction was recorded and the peaks corresponding to free 40S and 60S subunits, 80S ribosomes (monosomes) and polysomes are indicated.

**Supplementary figure S3.**
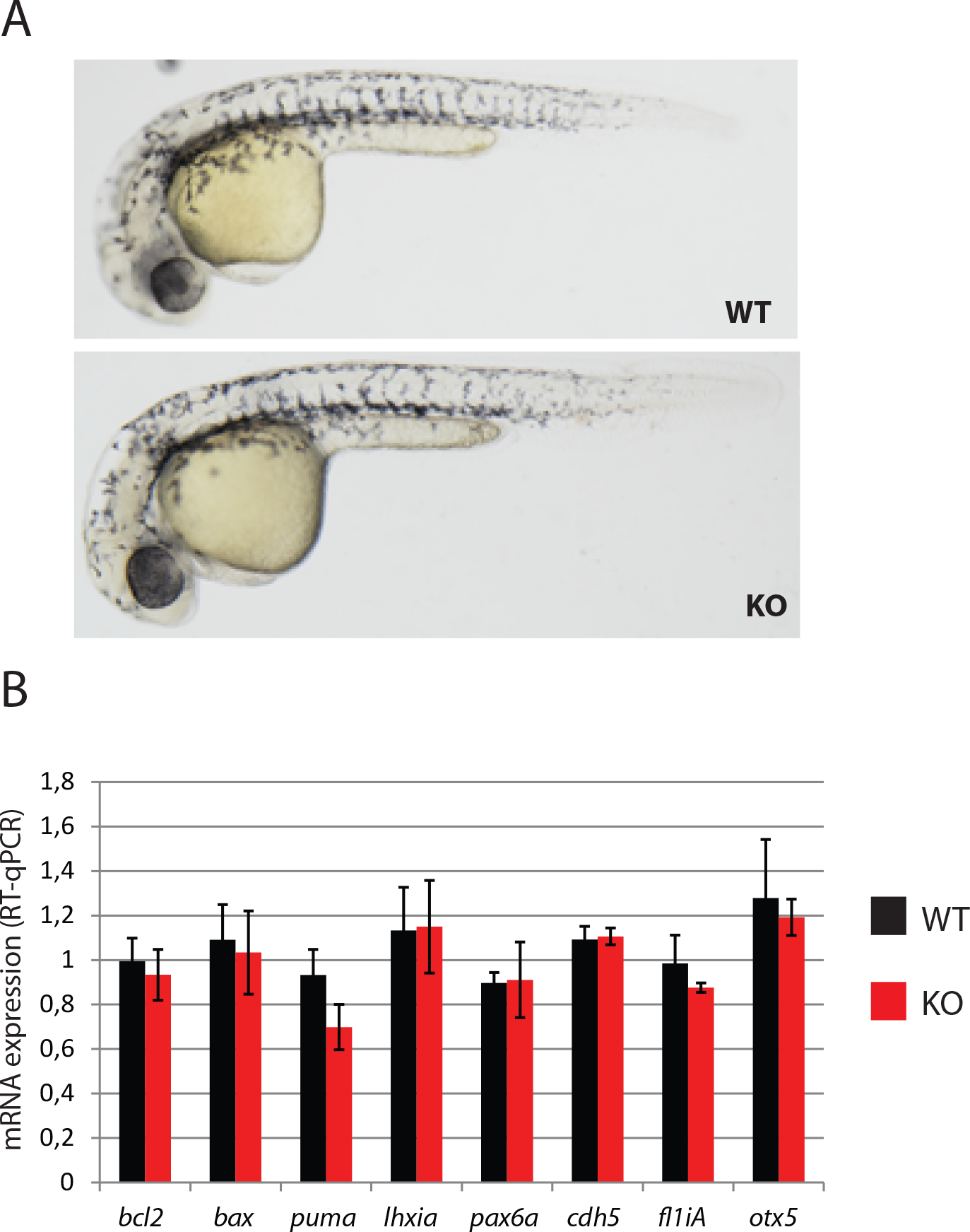
Deletion of *snord13* gene does not perturb zebrafish development. **A)** Bright field images of WT (up) and homozygous (down) *snord13* mutant embryos derived from the crossing of homozygous *snord13* females with heterozygous males. Pictures are representative of three independent crosses. **B)** Relative mRNA expressions determined by RT-qPCR in 30 hpf WT and SNORD13*-*KO embryos (3 independent experiments with at least 15 animals per condition). The tested genes are listed below the histogram (black and red histograms correspond to WT and KO, respectively).

**Supplementary figure S4.**
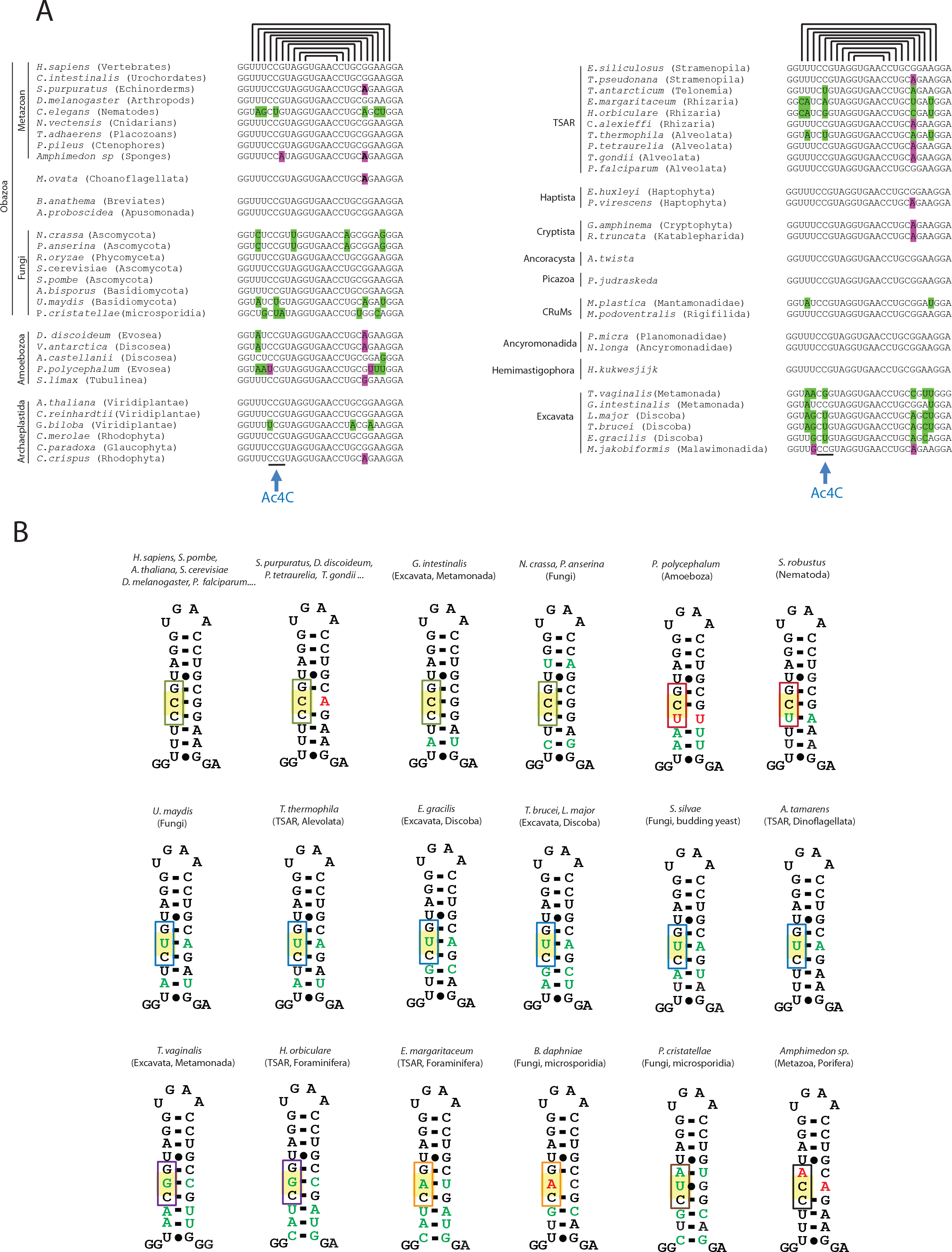
Phylogenetic conservation of helix 45 across evolution. Sequence **(A)** and schematic representations **(B)** of helix 45 in representative species belonging to each eukaryotic supergroup. Green bases represent compensatory base changes, as compared to human helix 45, while red bases alter the geometry of helix 45. The triplet sequence motif is framed.

**Supplementary figure S5.**
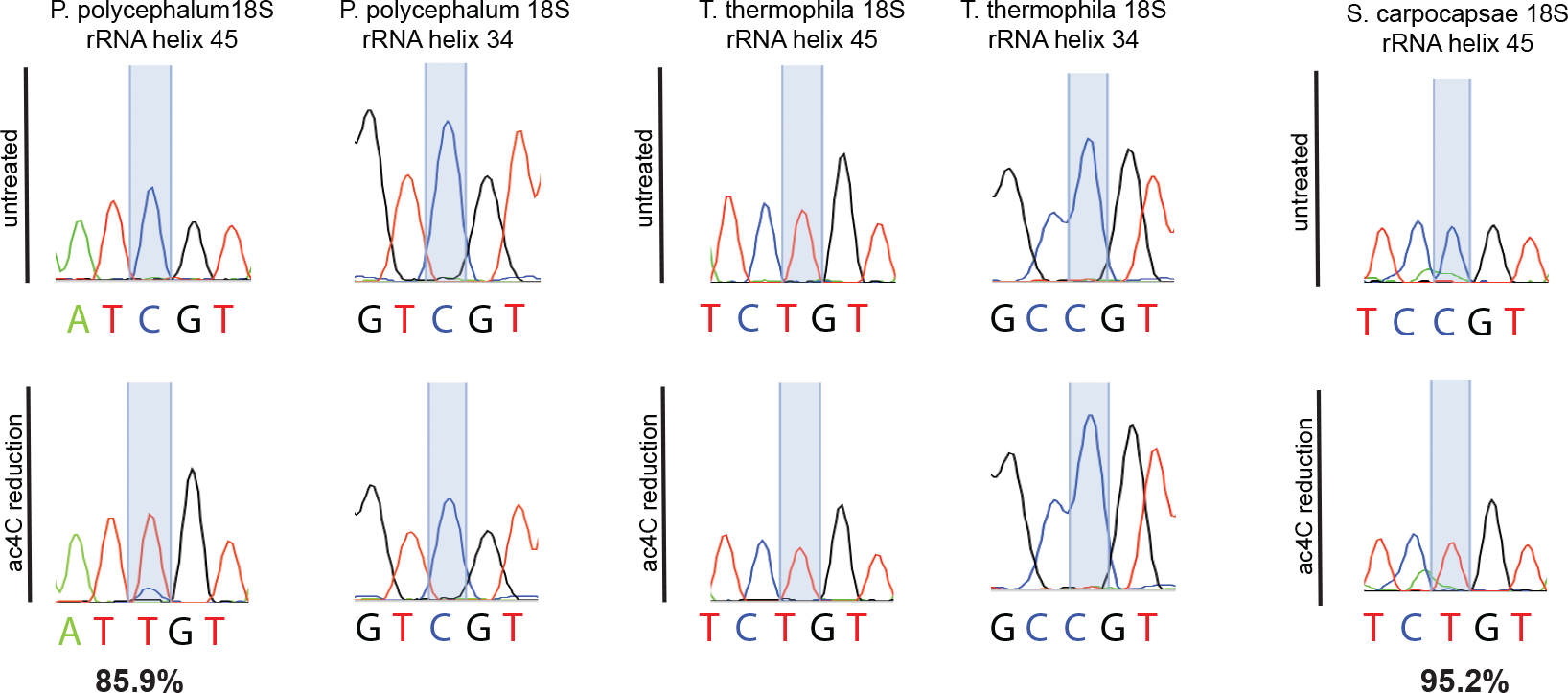
Probing rRNA acetylation in *T. thermophila, S. carpocapsae* and *P. polycephalum*. Sanger DNA sequencing of RT-PCR products obtained after borohydride treatments of total RNA extracted from *P. polycephalum* (left) *T. thermophila* (middle), *S. carpocapsae* (right). % of misincorporation at helix 45 is indicated below each electropherogram.

**Supplementary figure S6.**
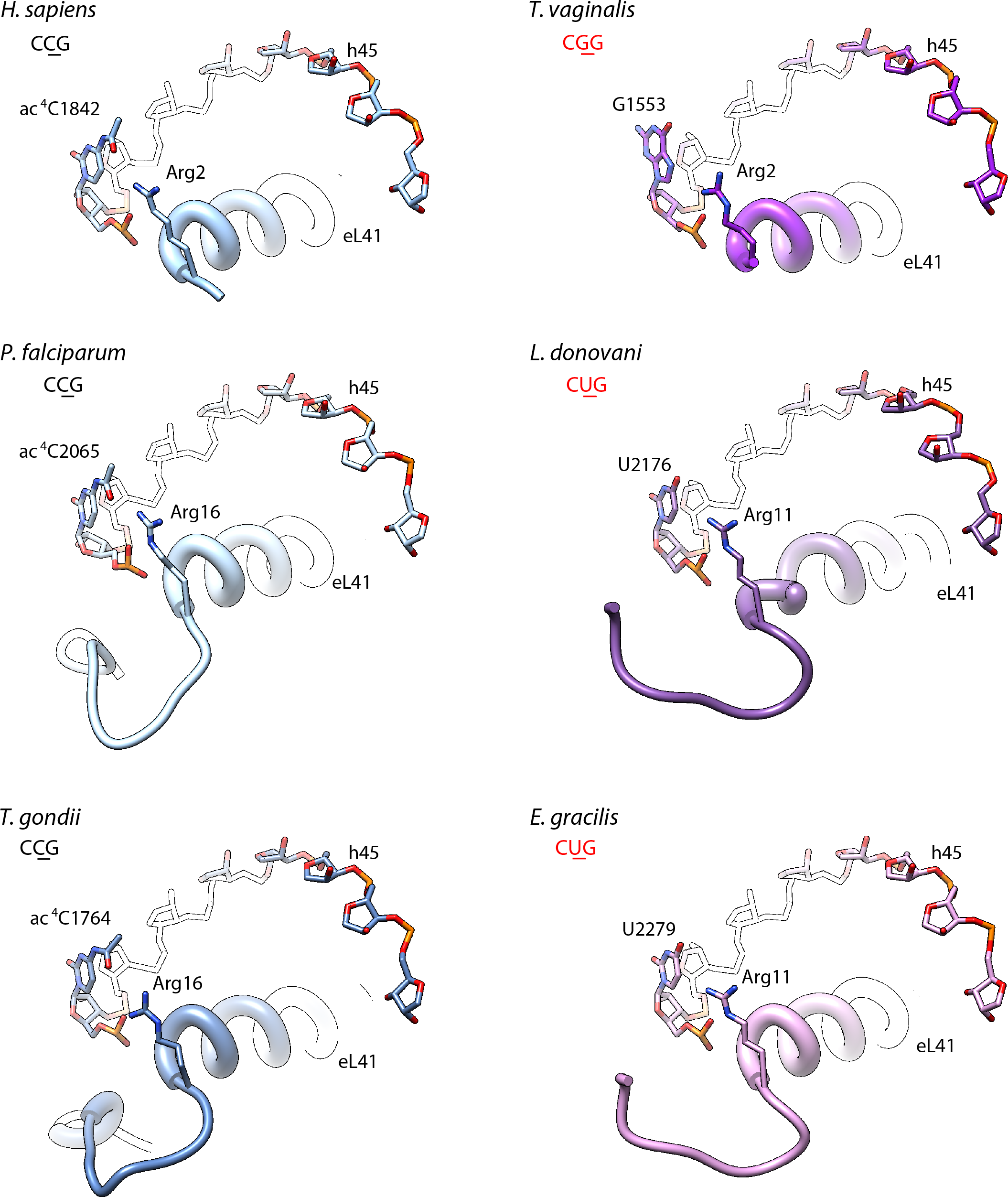
rRNA h45 acetylation does not seem to affect Rpl41 positioning. Spatial alignment of Ribosomal 3D structures of various species, carrying either acetylated C (left panel) or unmodified U or G (right panel) on rRNA helix 45 (h45) revealed no change in the location of ribosomal protein Rpl41 (eL41). For each aligned species (indicated on the top left of each vignette), the modified/unmodified nucleotide of interest is shown both on the structure as well as on the 3-nucleotide sequence by an underlined character. For spatial cueing, lateral chain of an N-terminal Arginine of Rpl41, one of the closest aminoacids of the nucleotide of interest, is fully displayed.

**Supplementary data S1.**
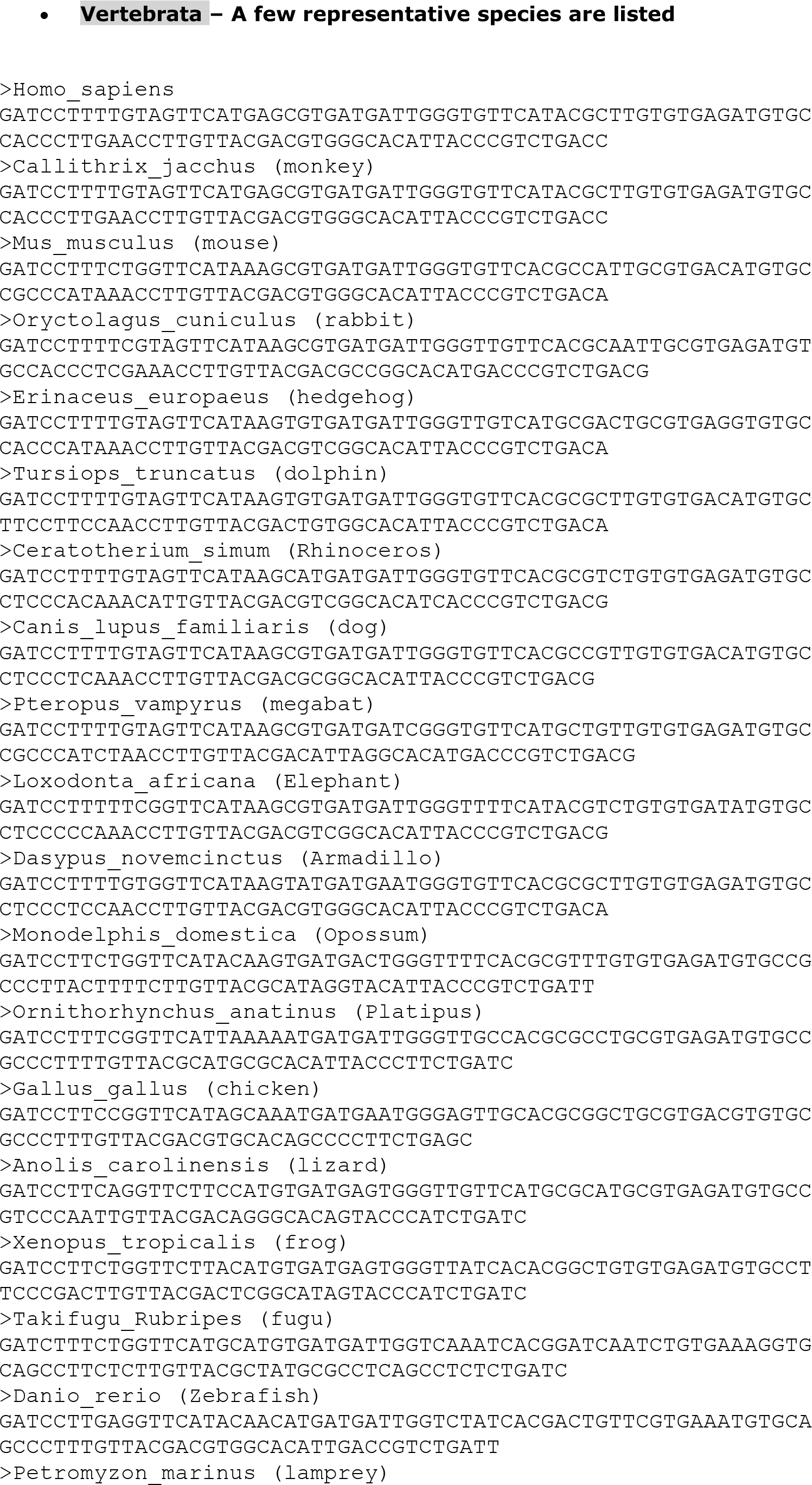
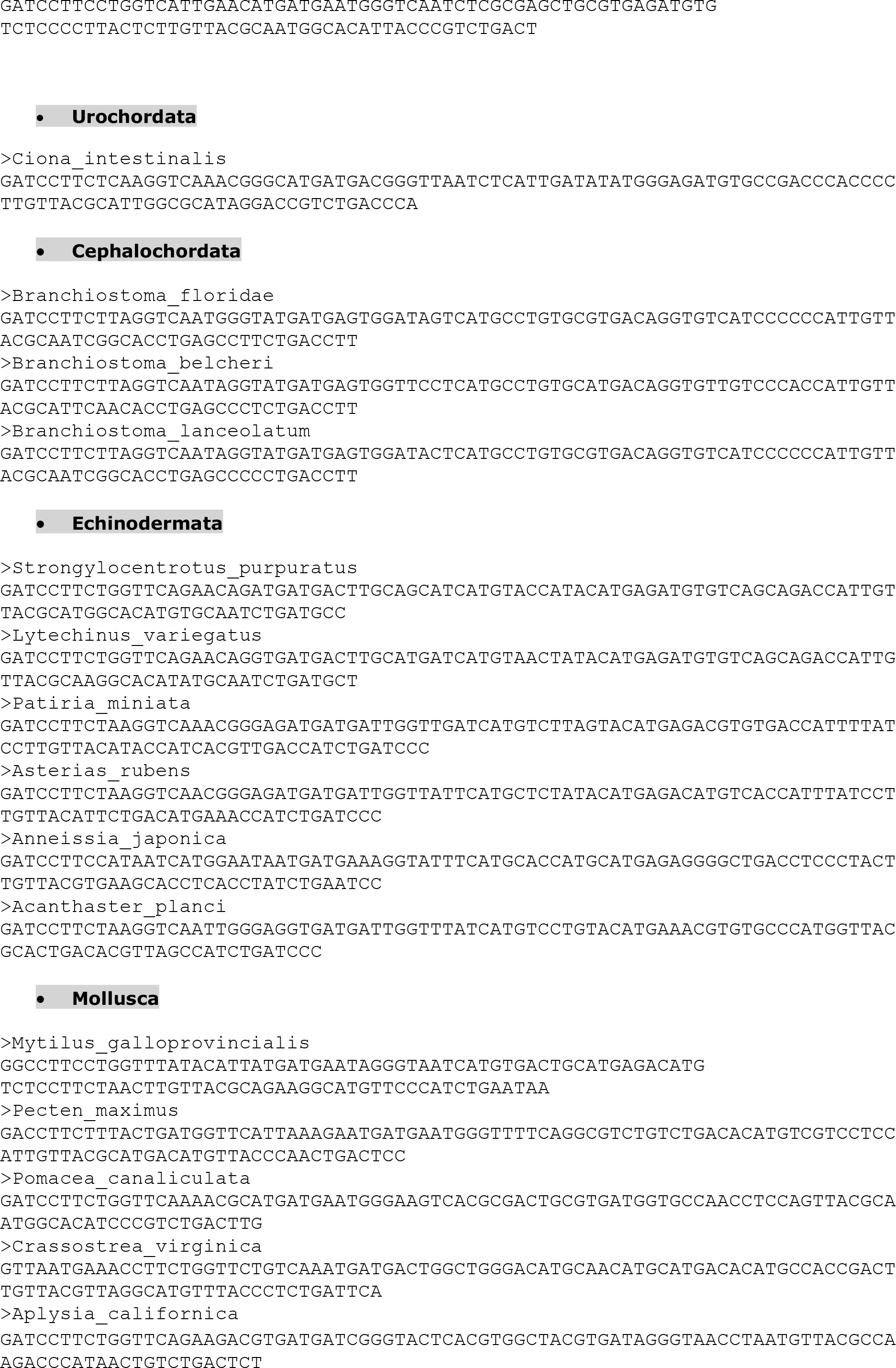
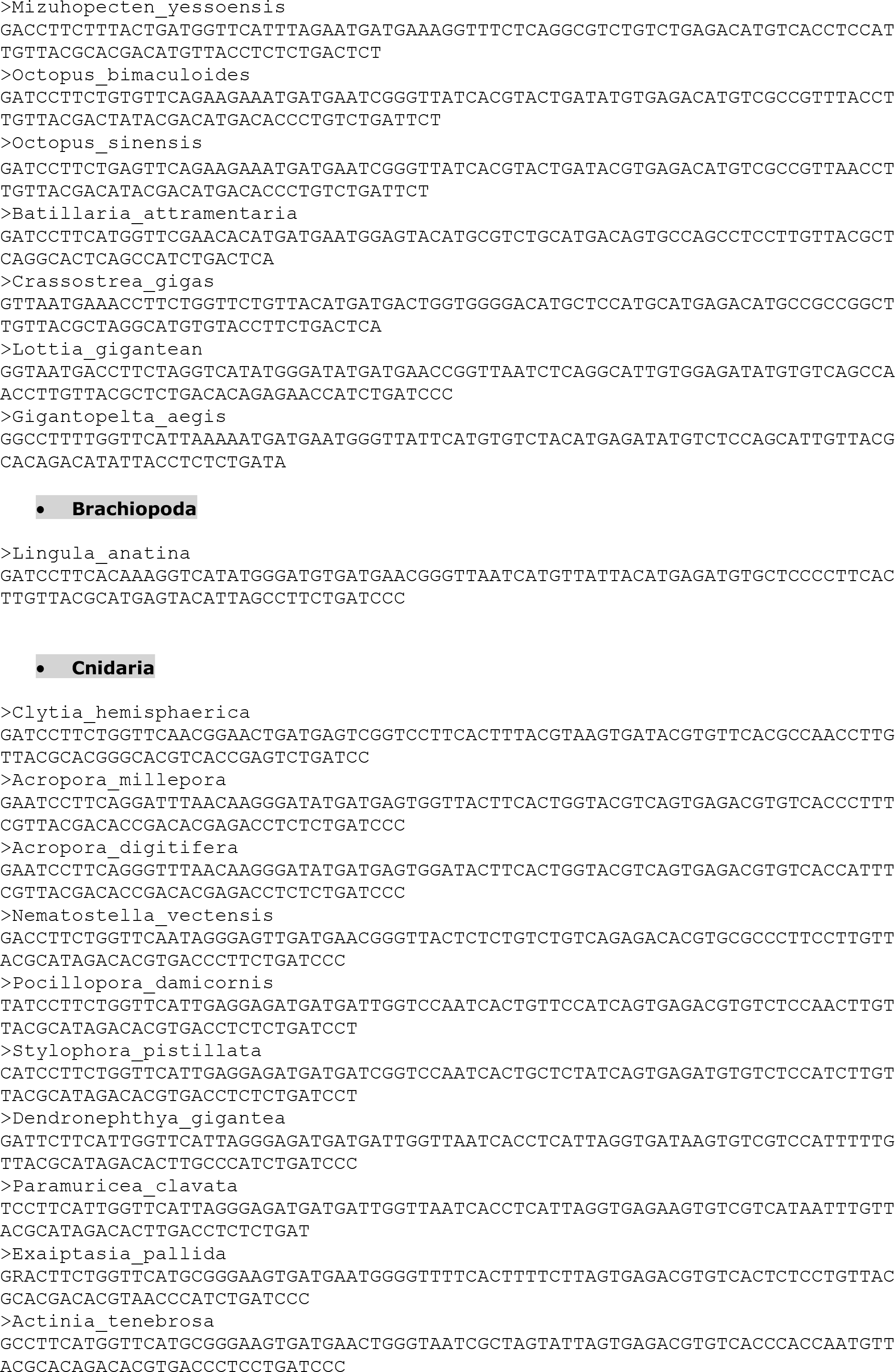
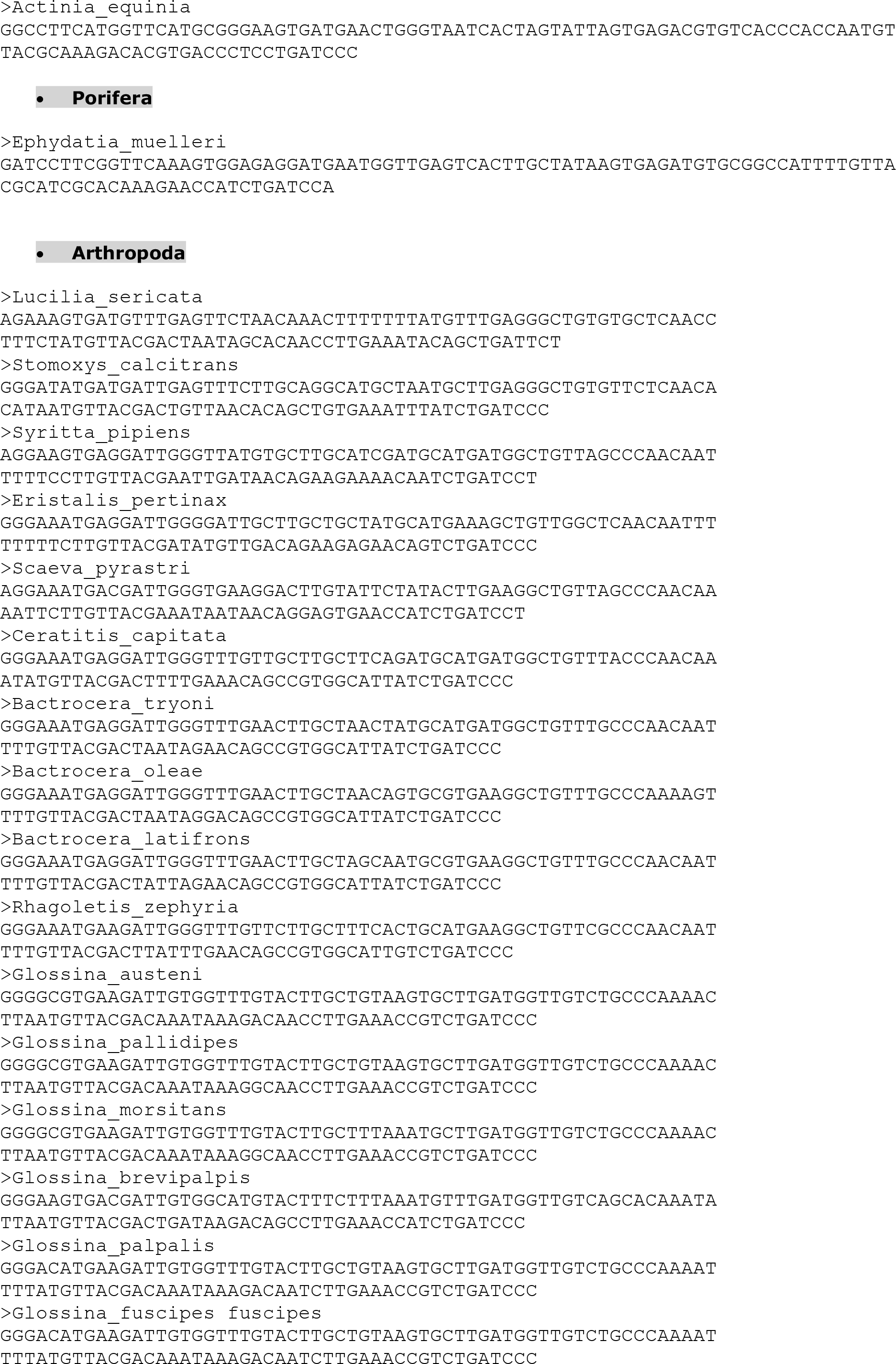
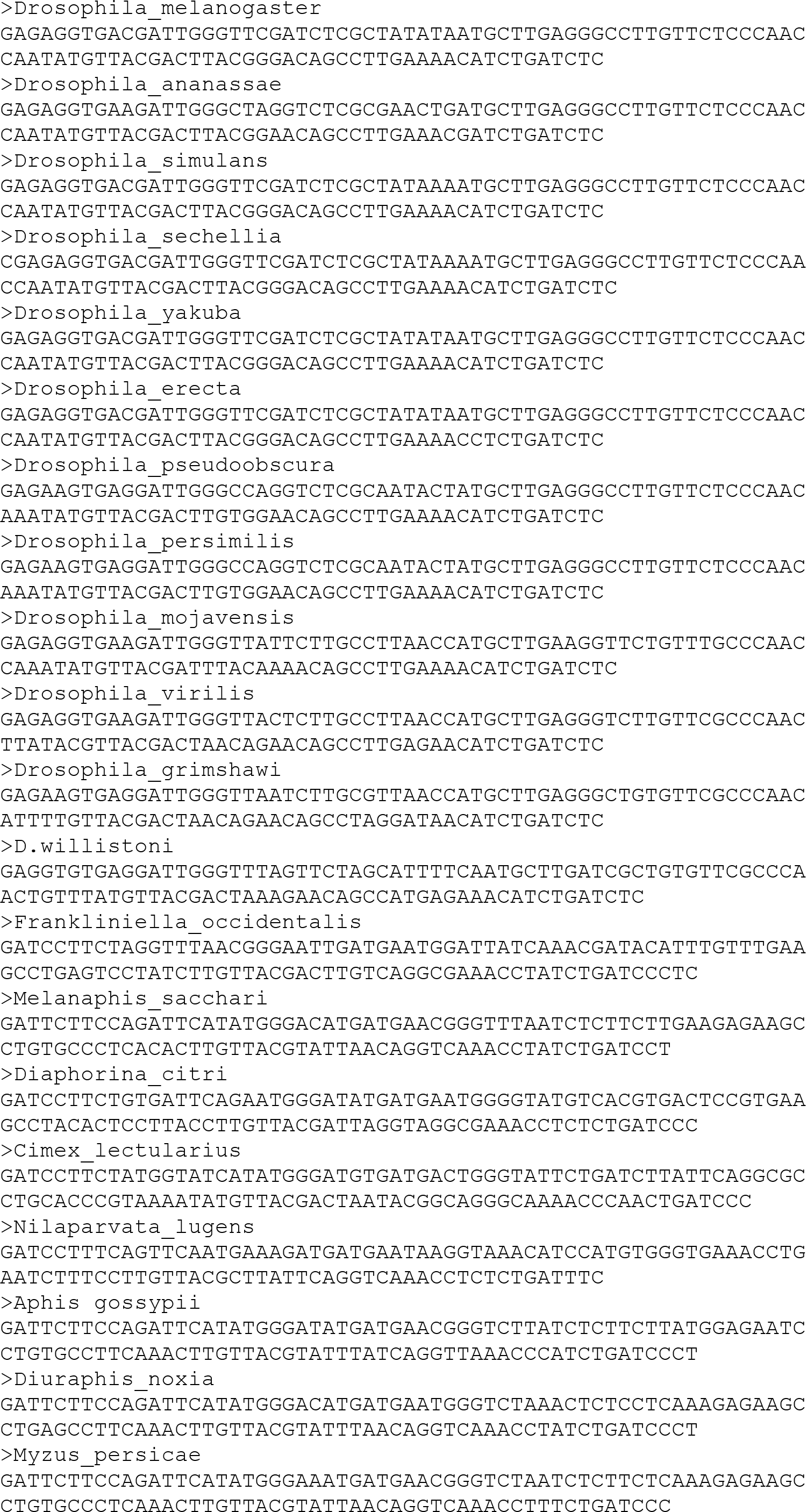
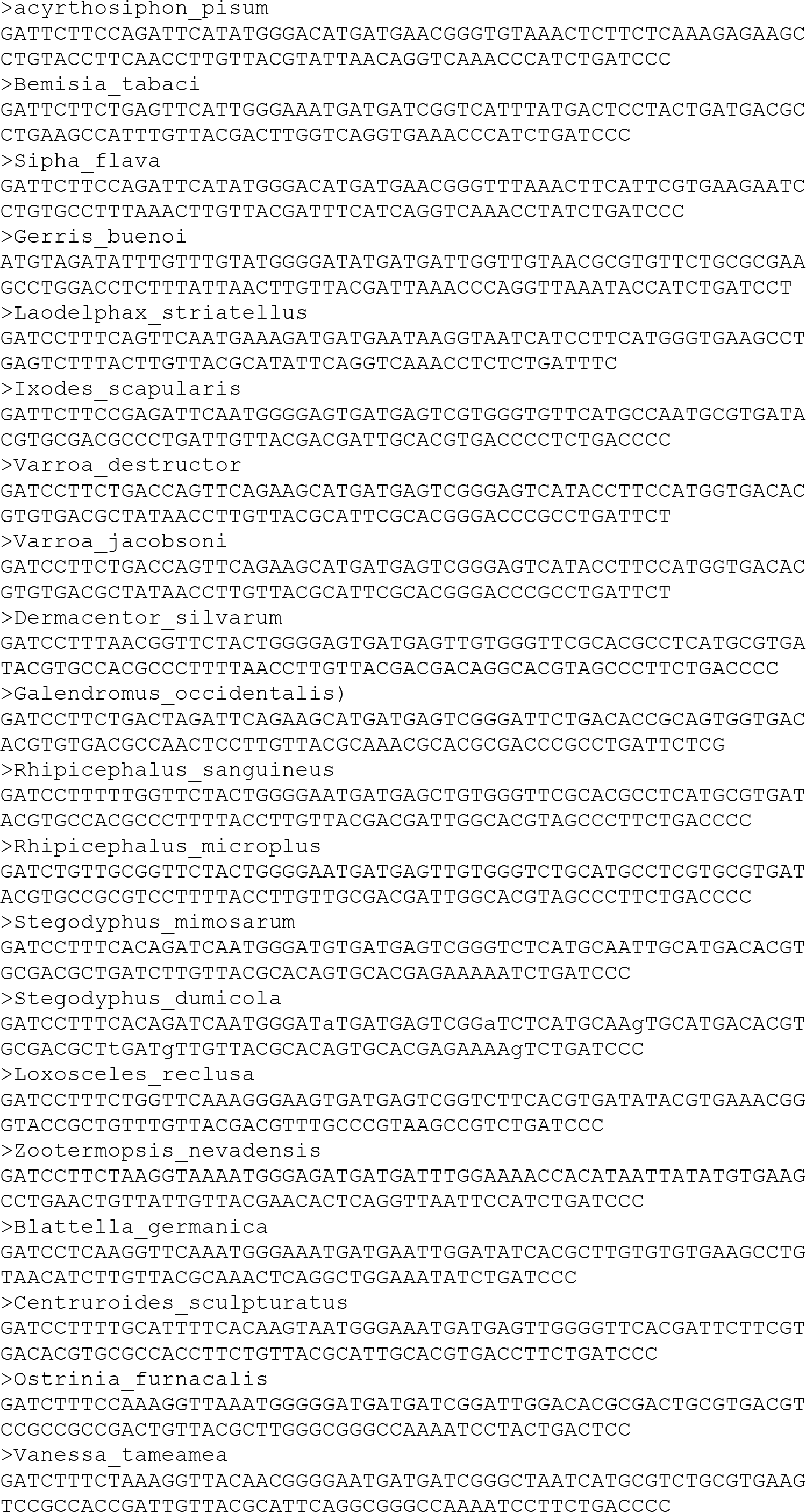
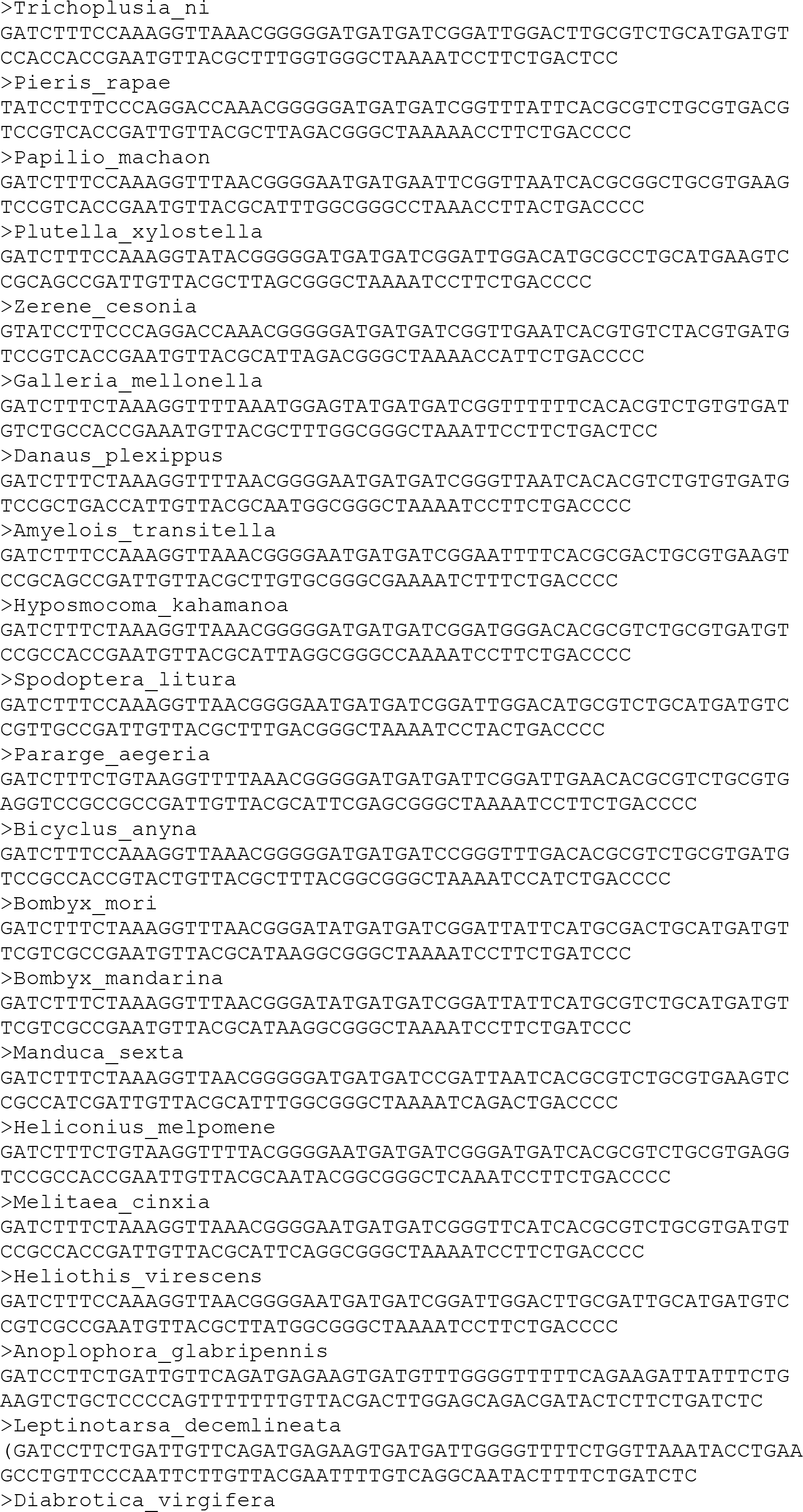
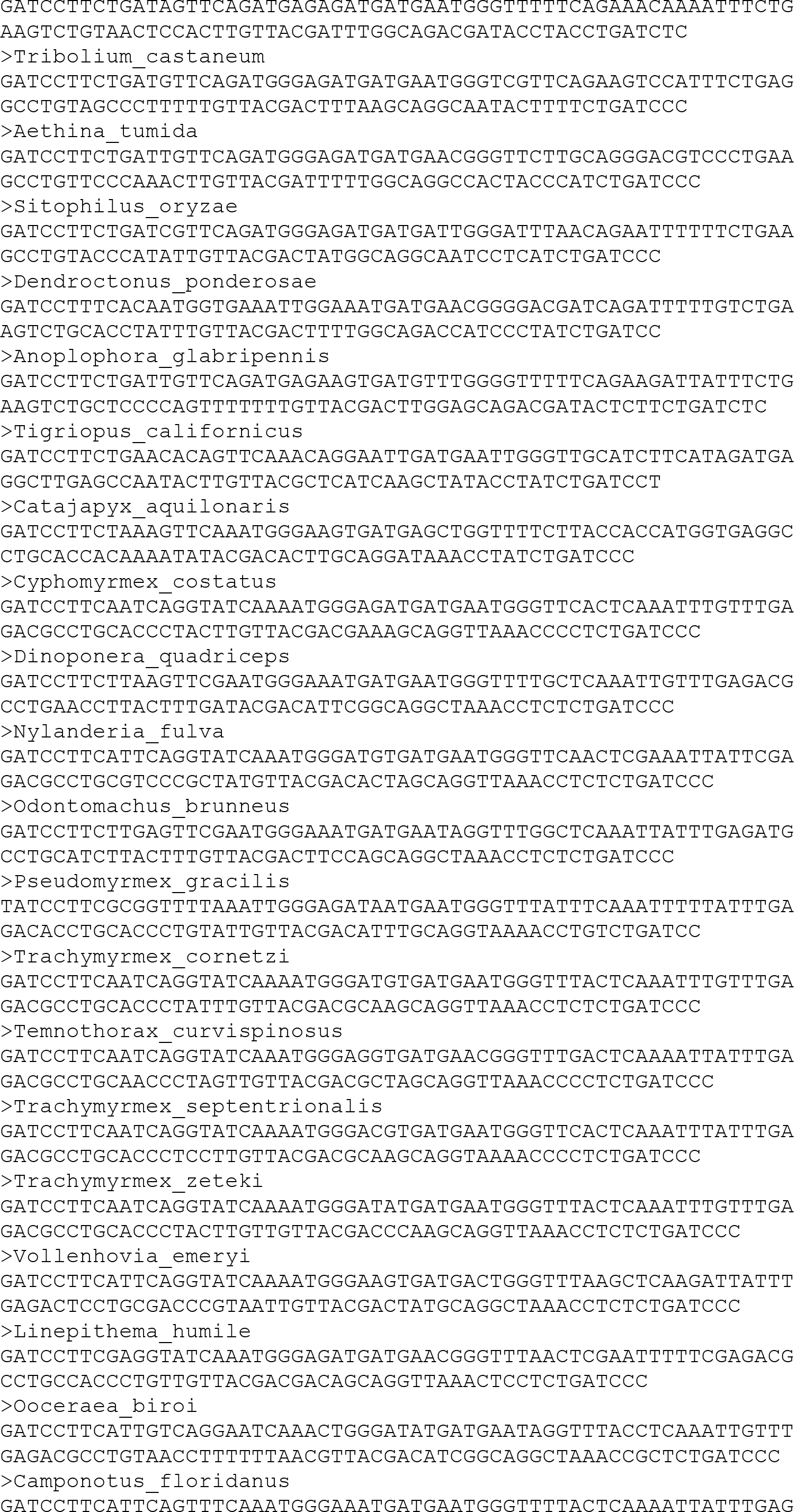
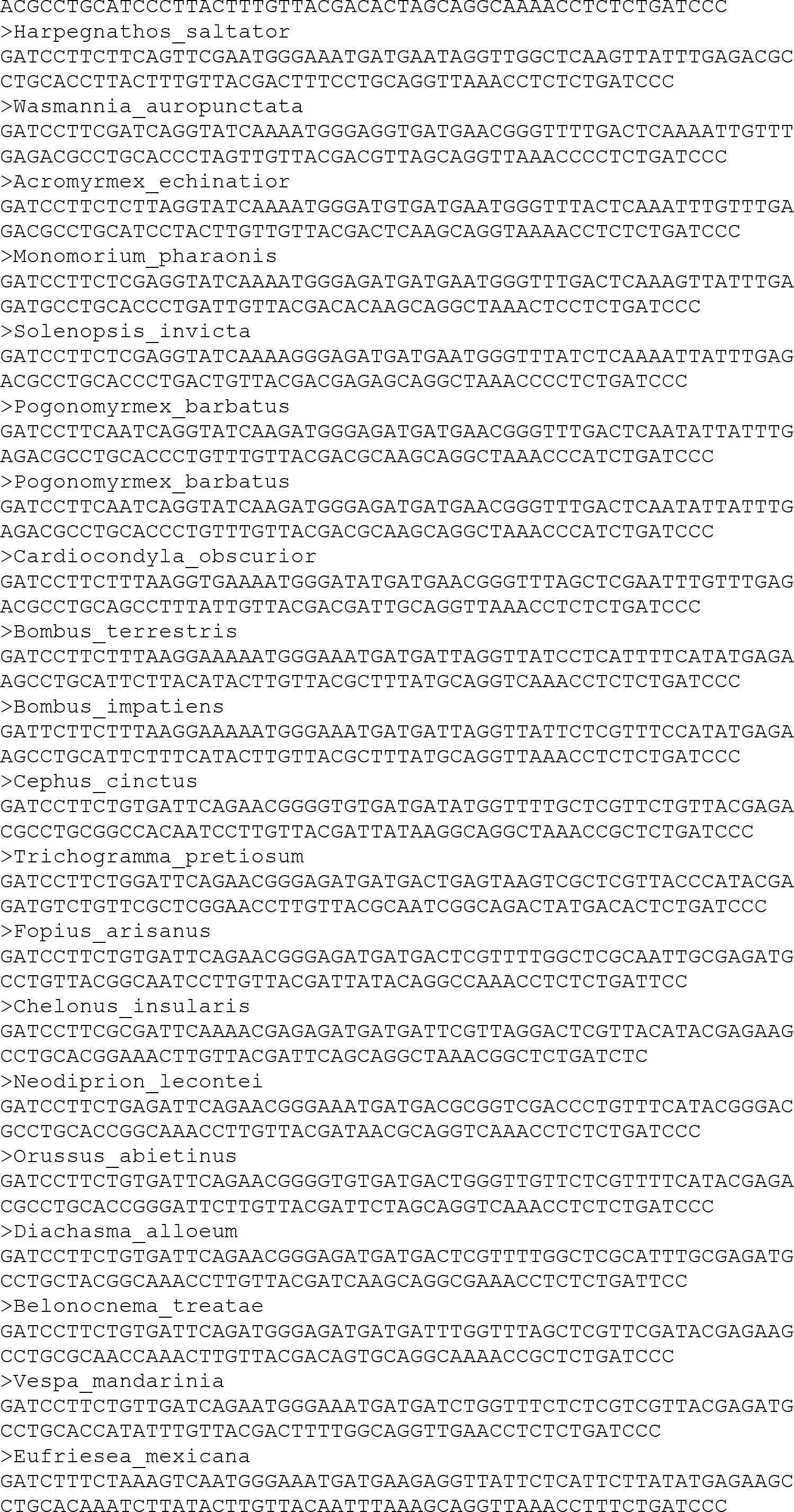
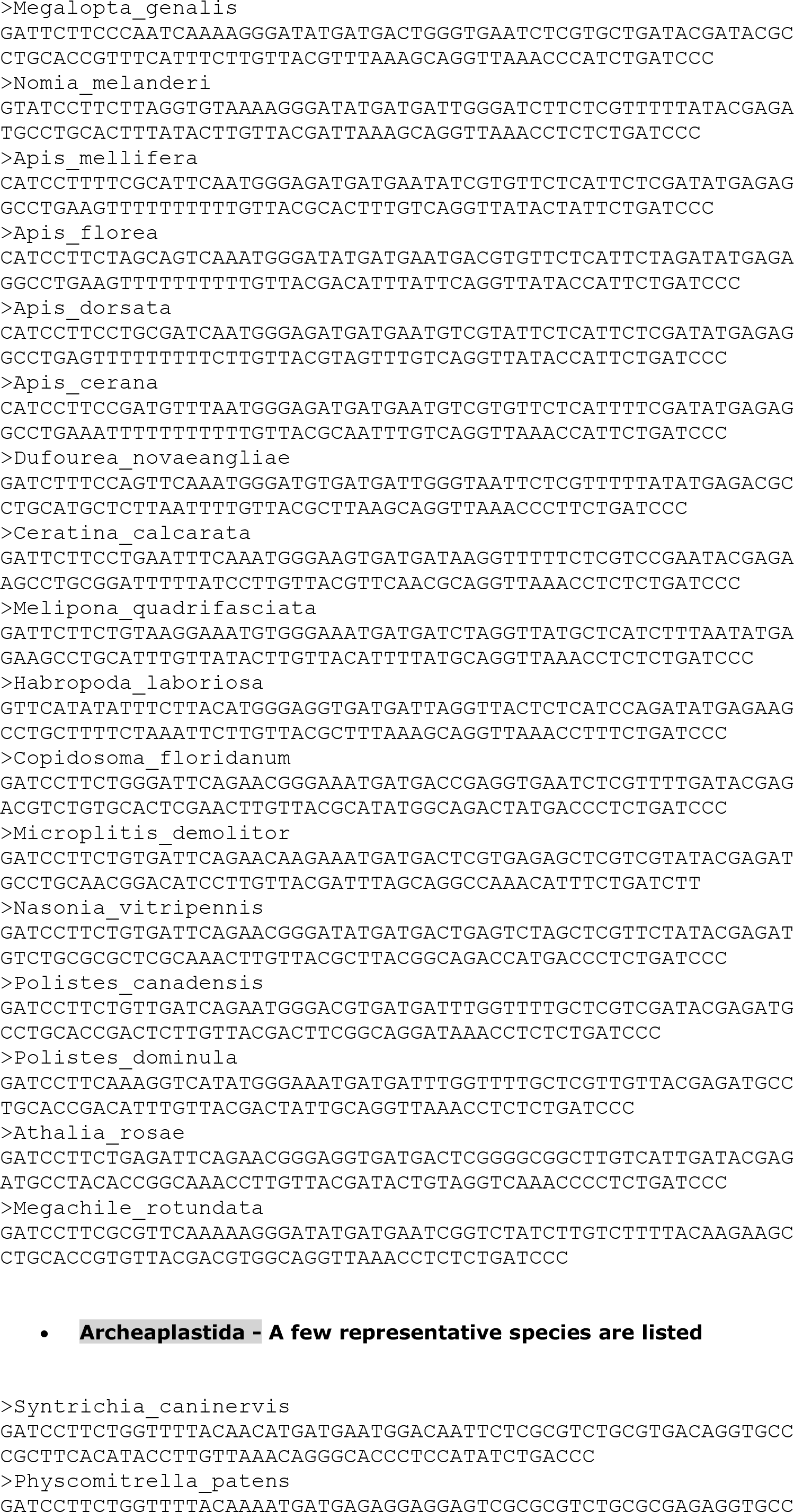
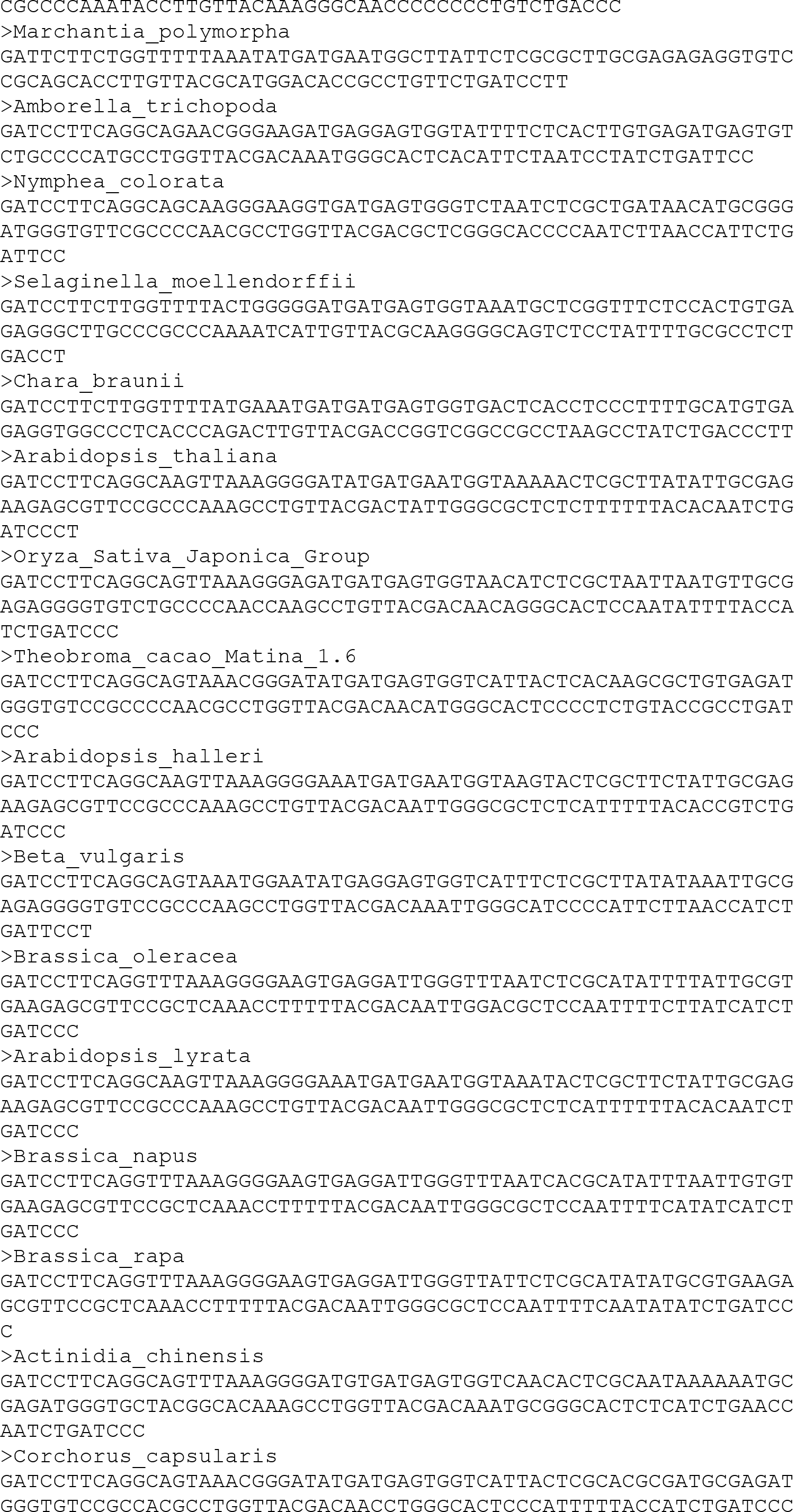
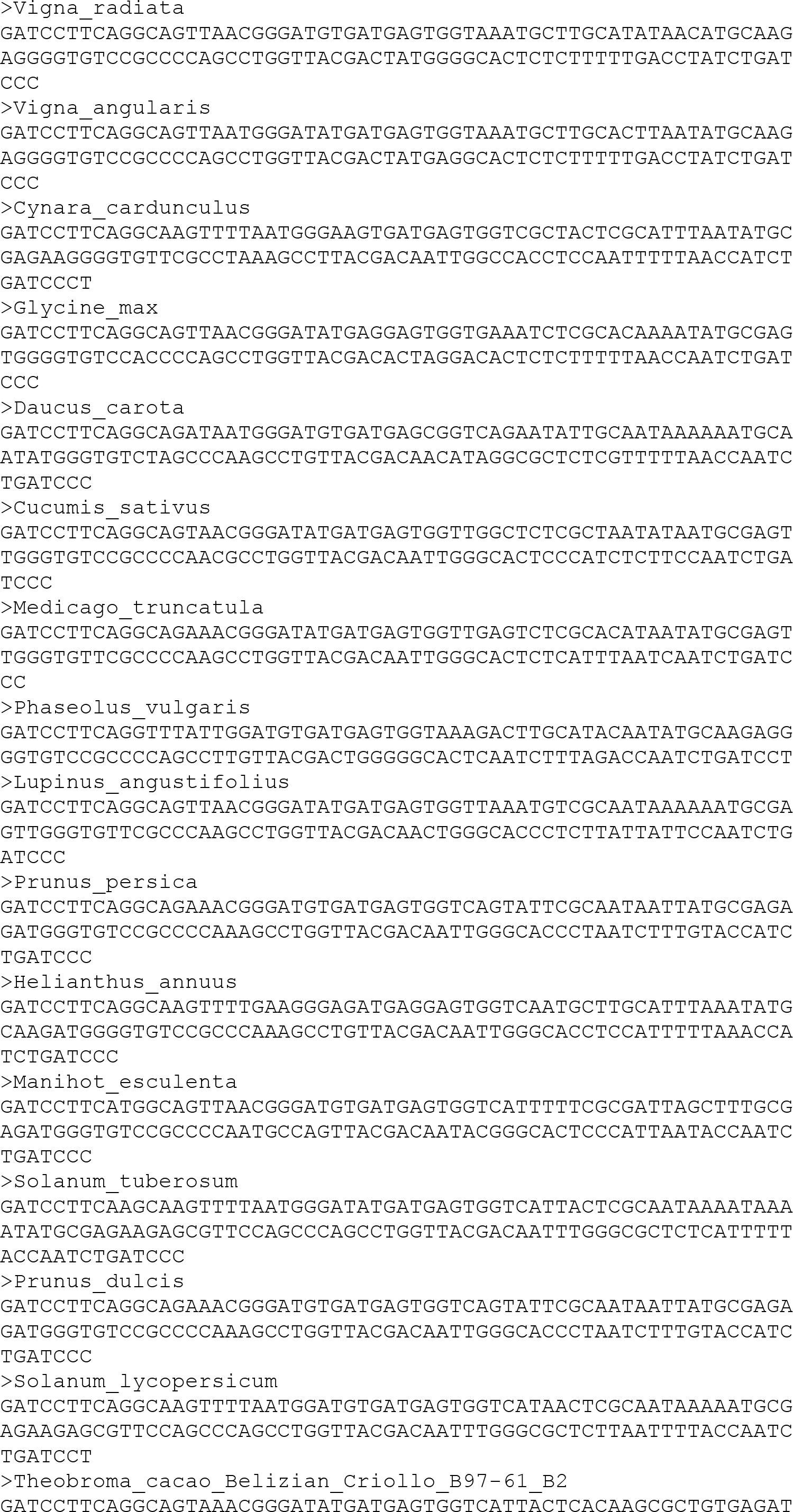
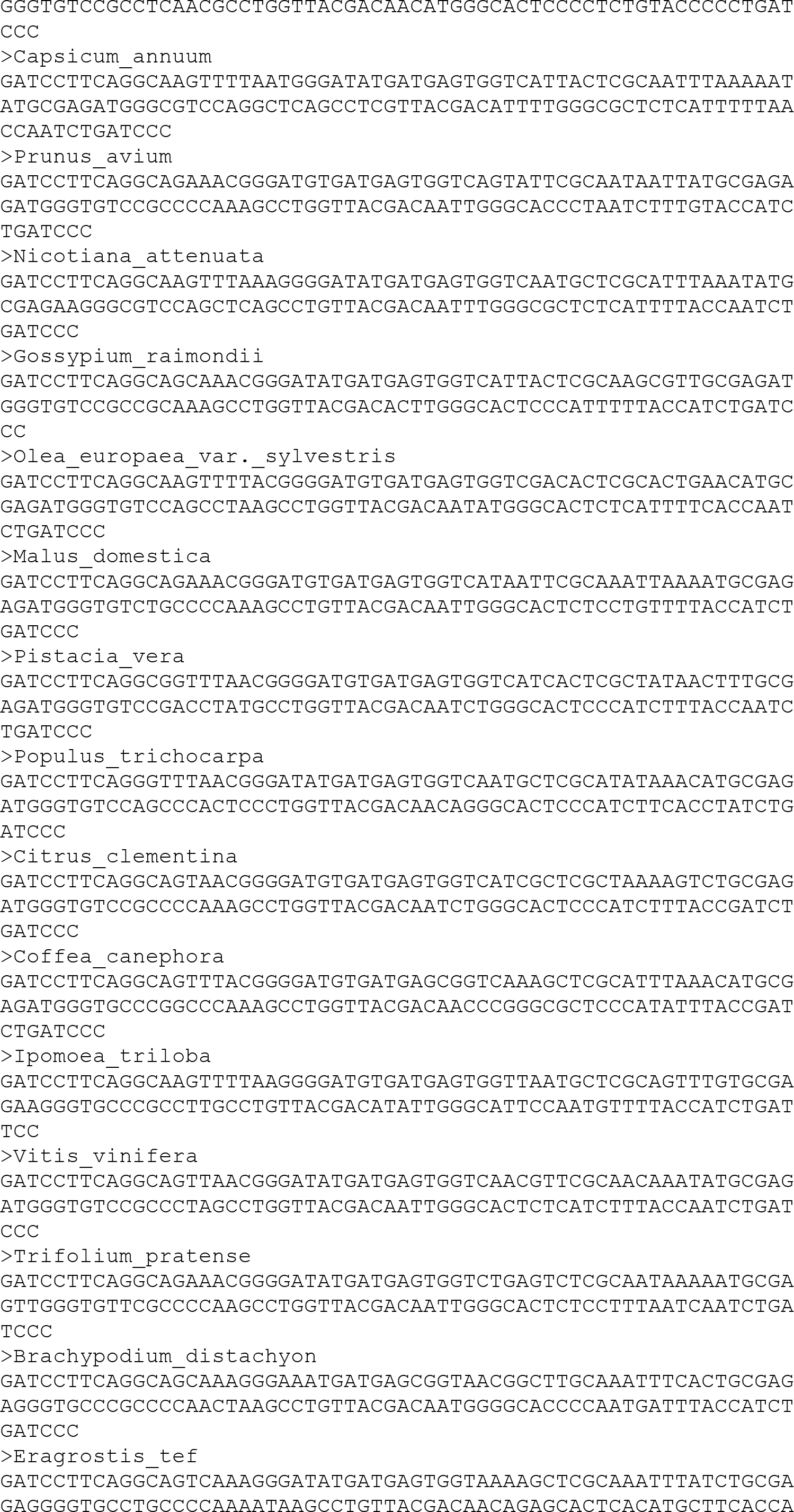
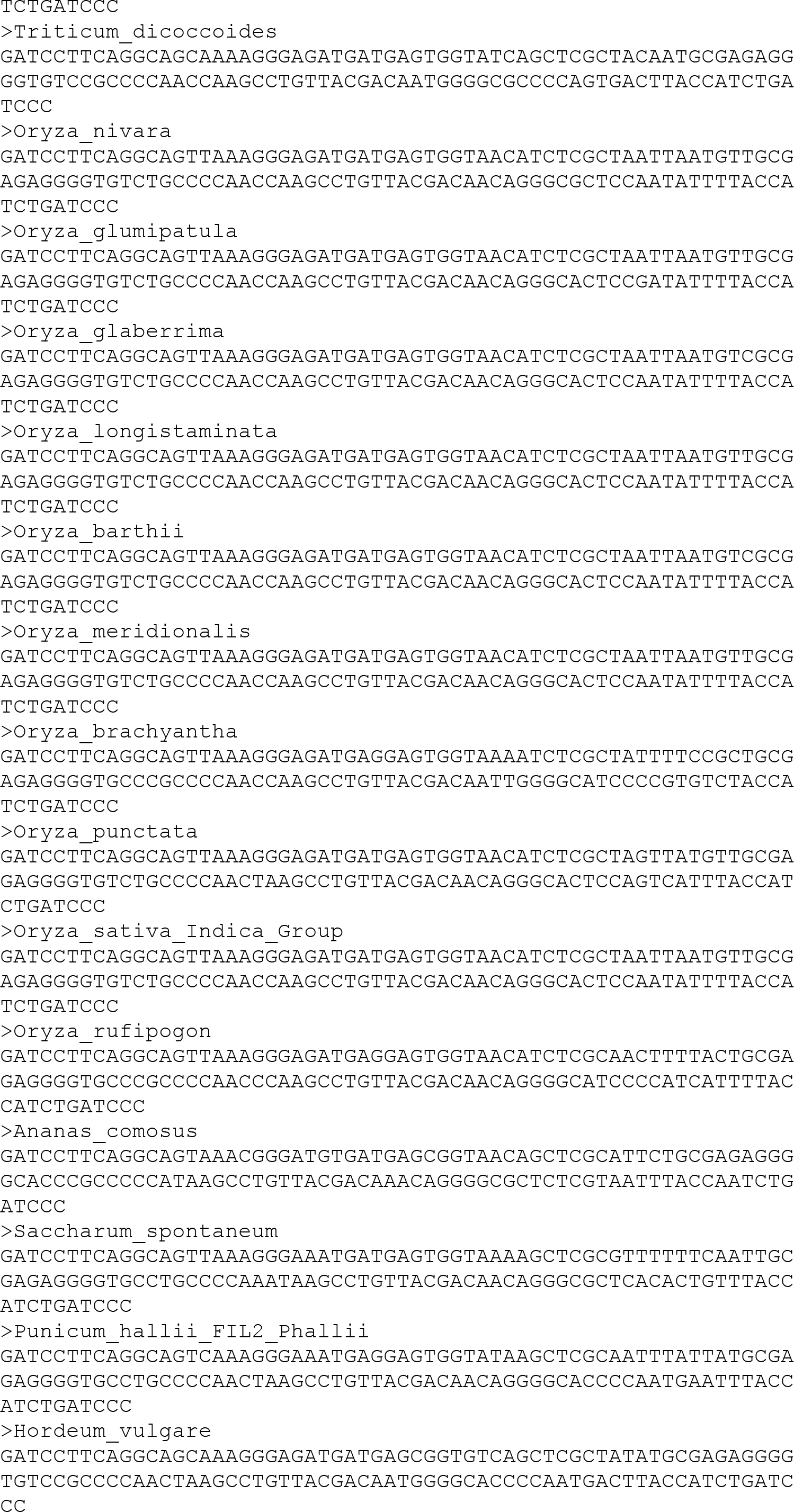
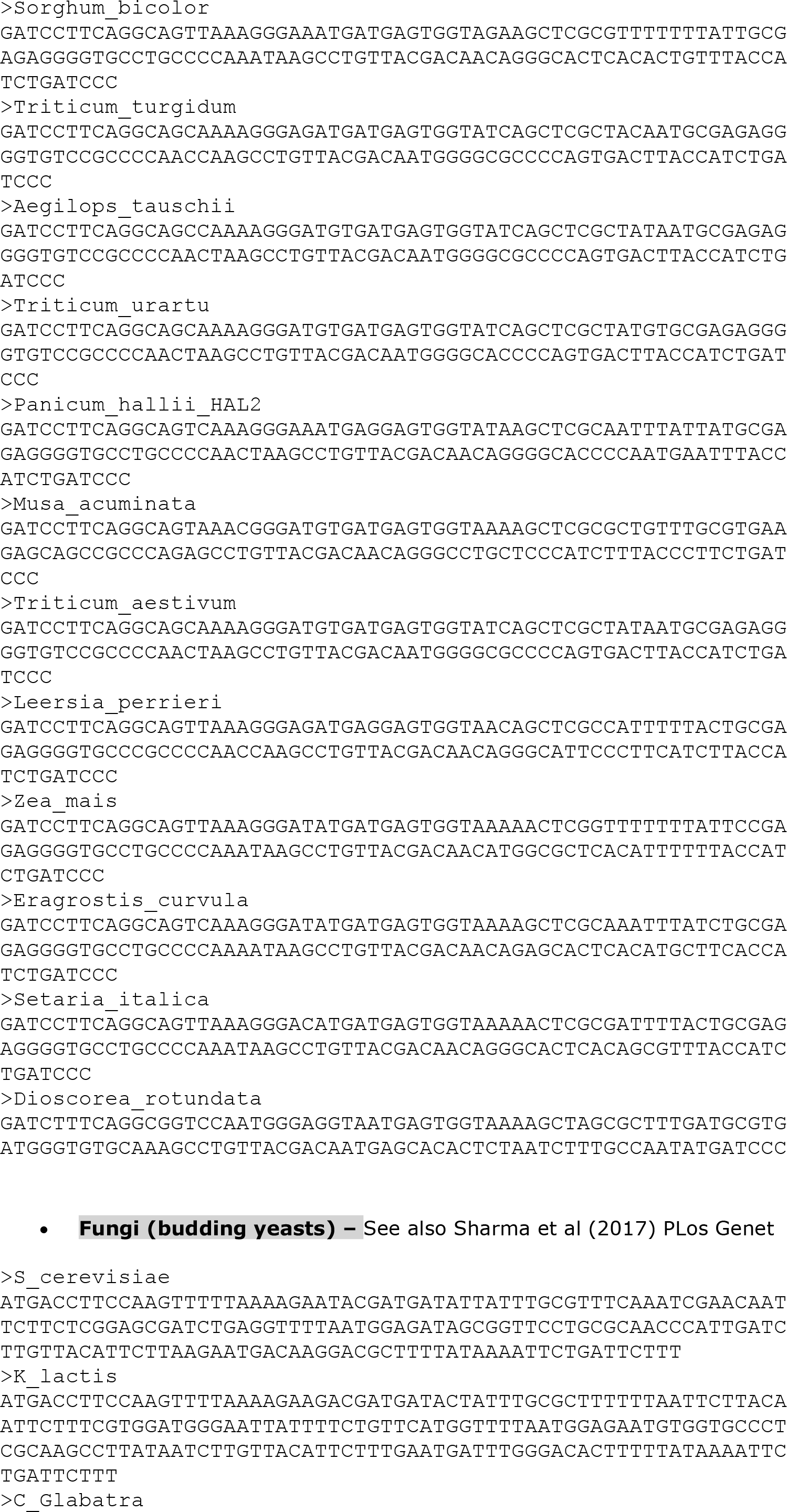
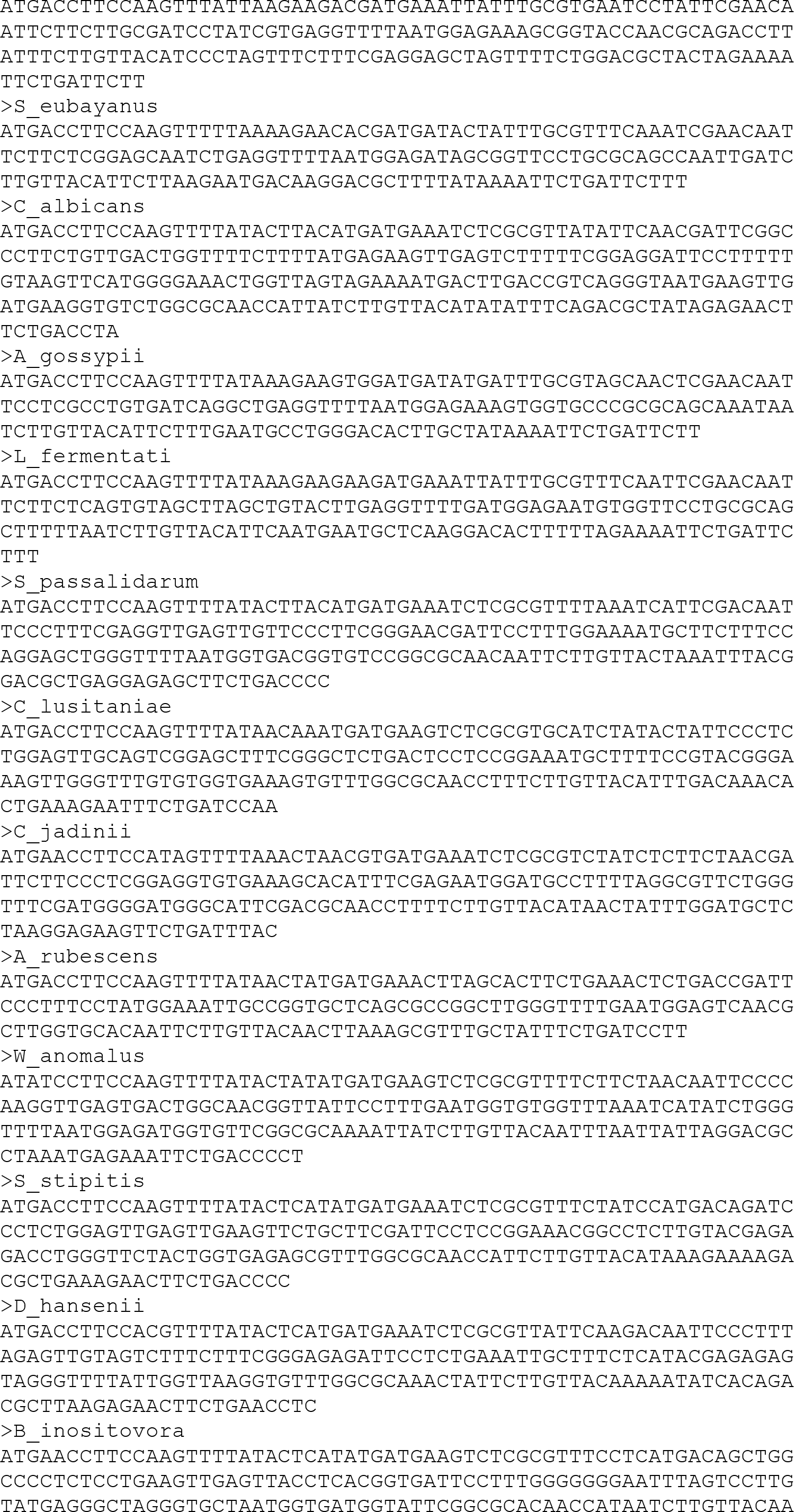
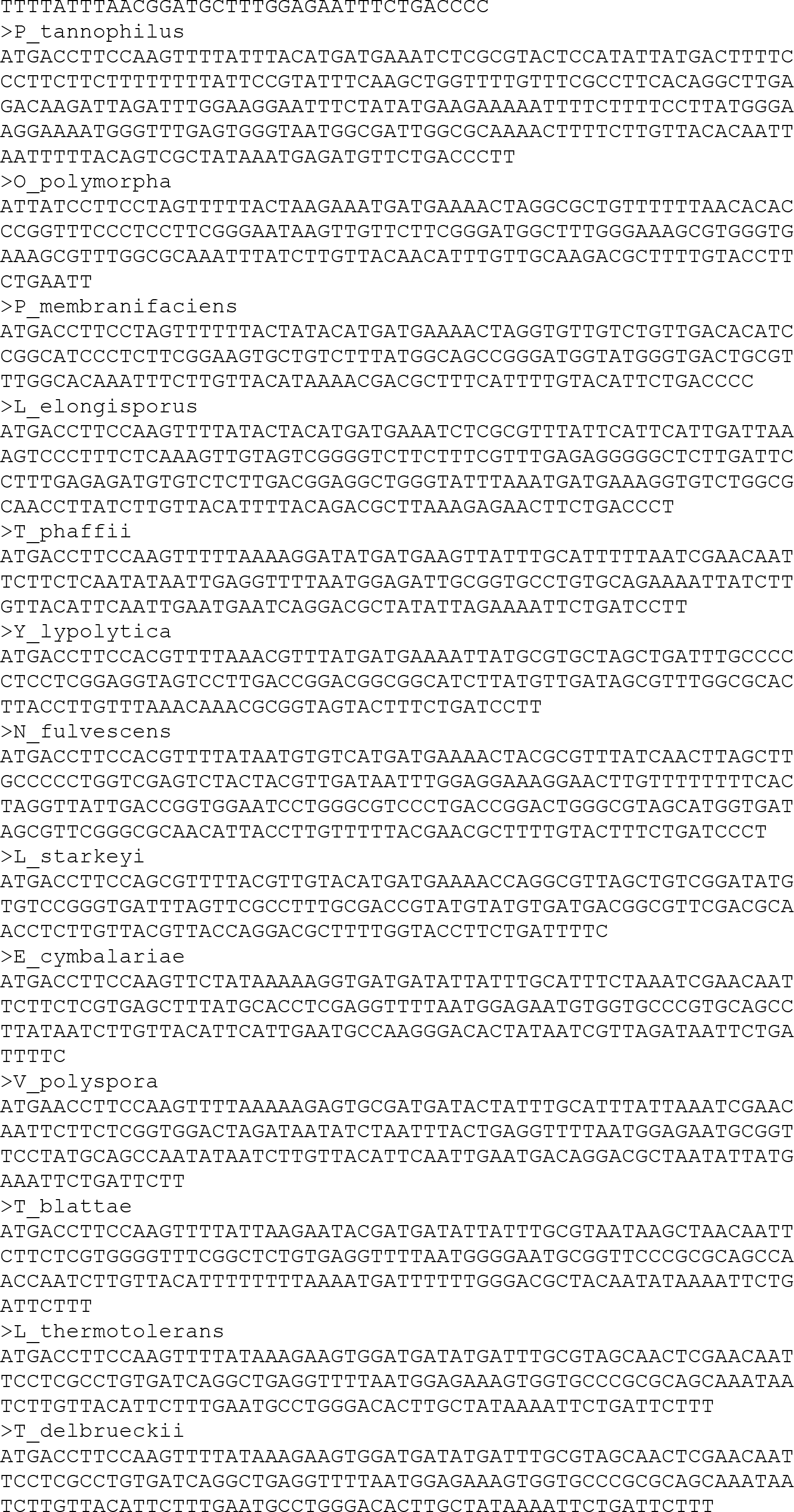
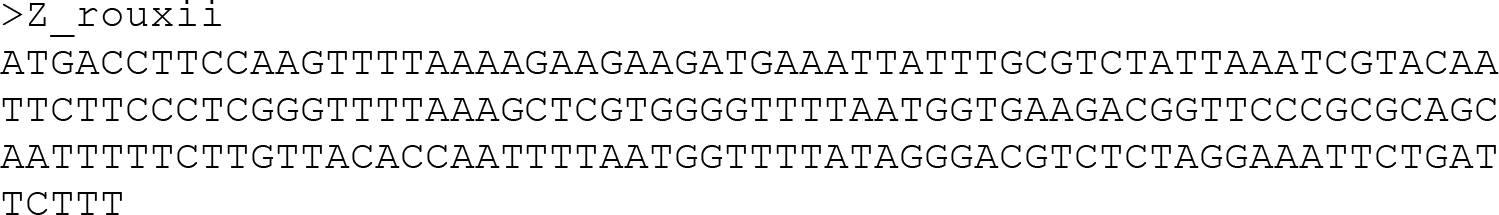
Examples of newly-identified SNORD13 sequences.

**Supplementary data S2.**
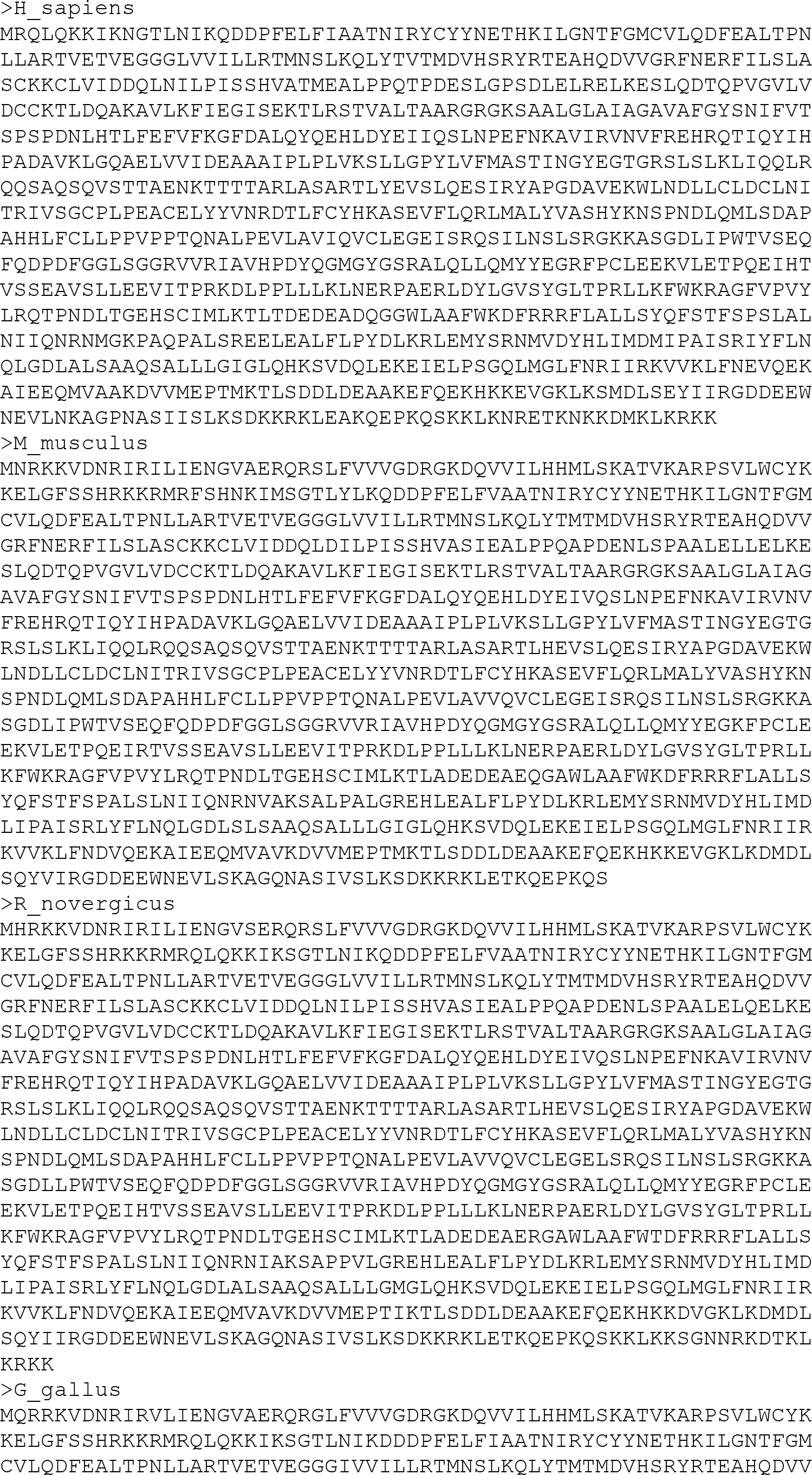
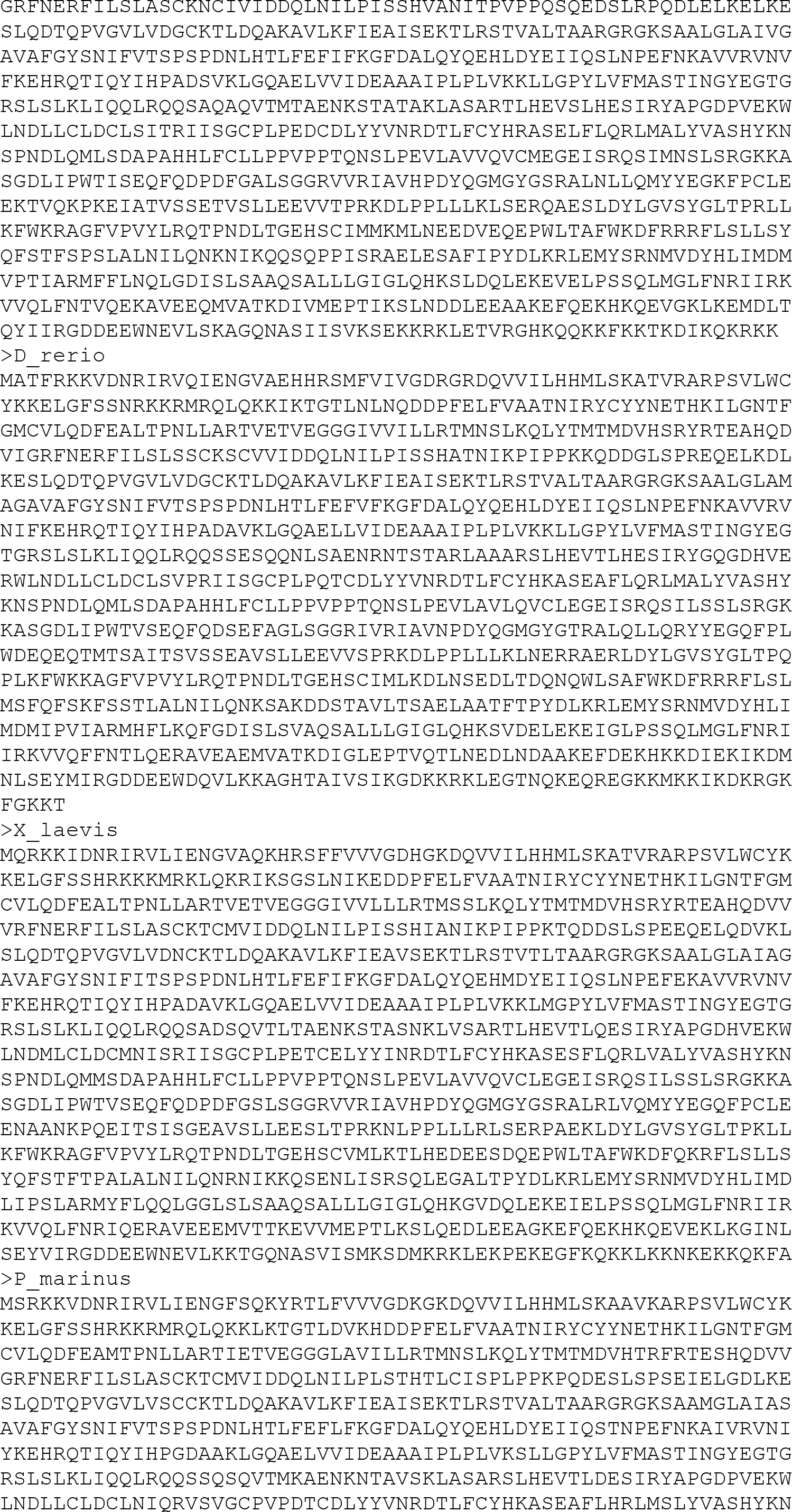
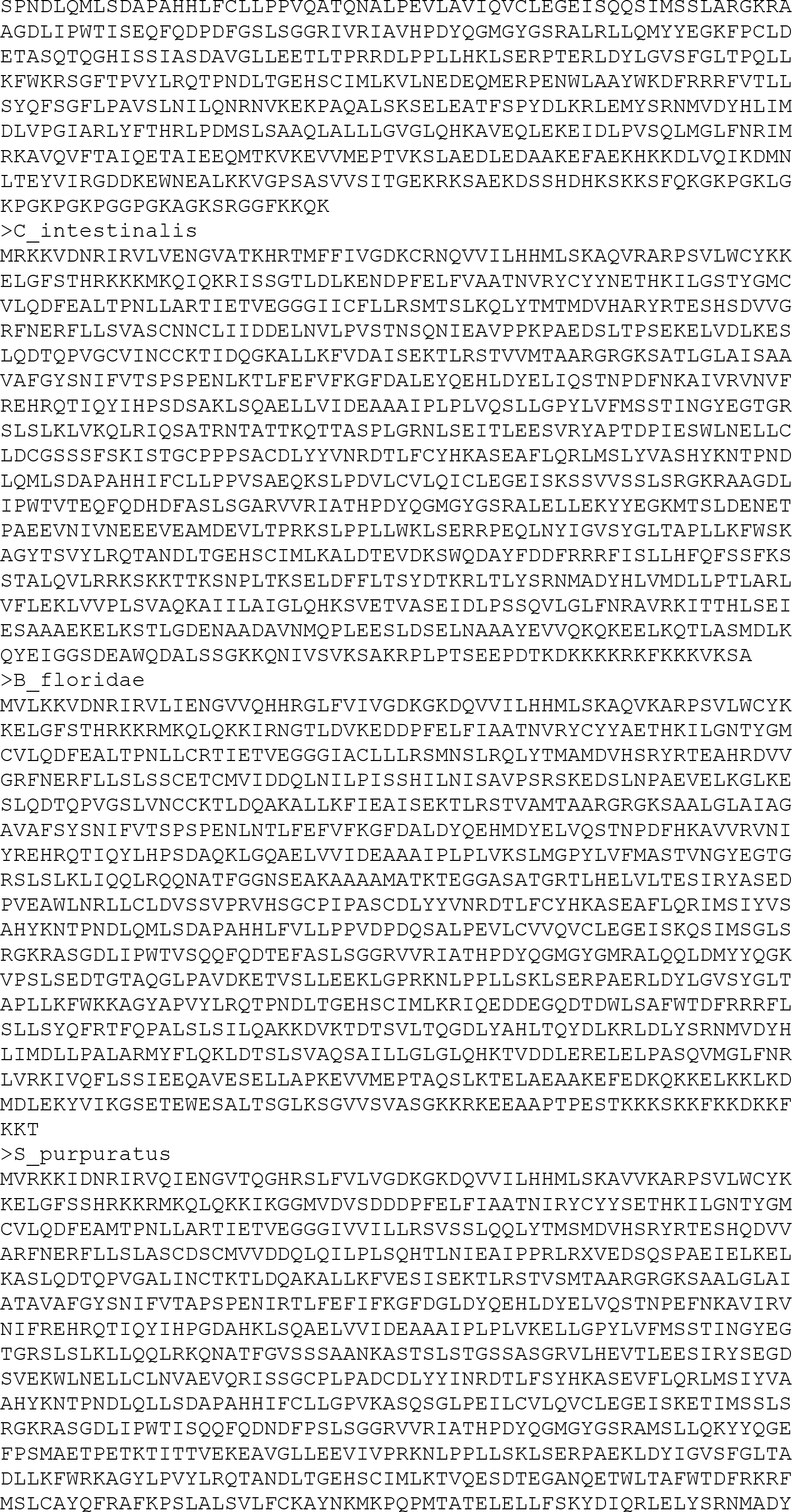
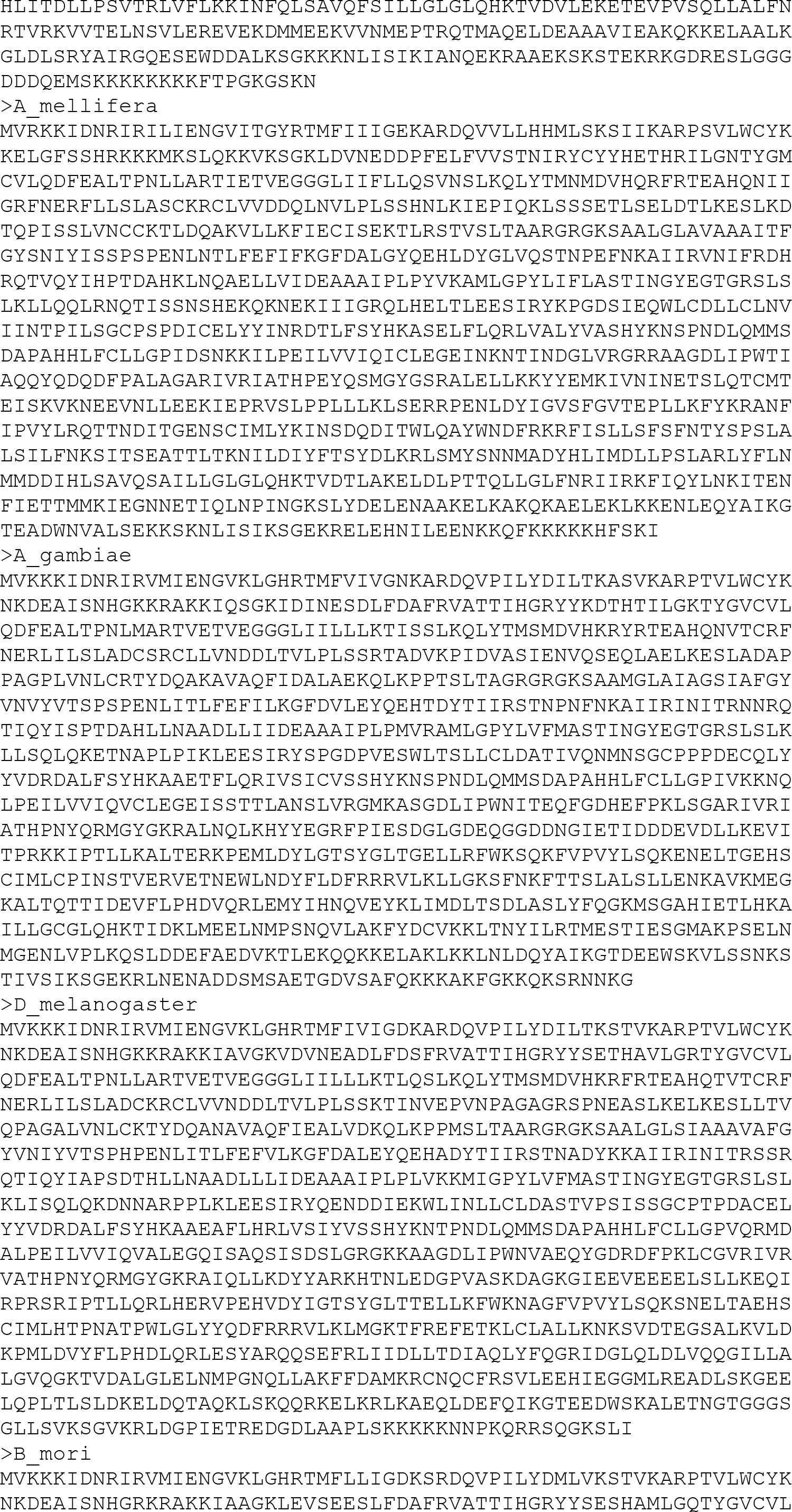
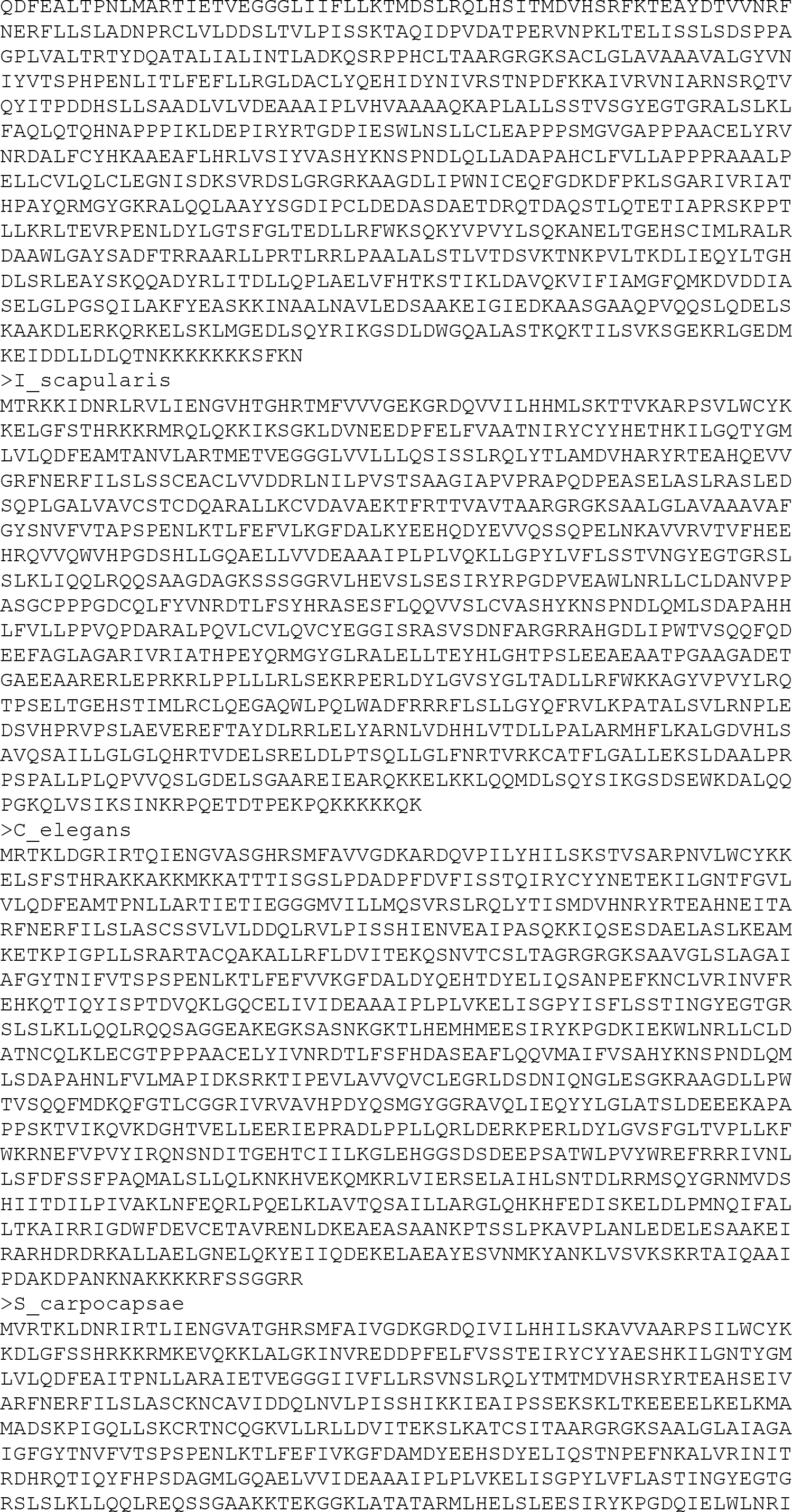
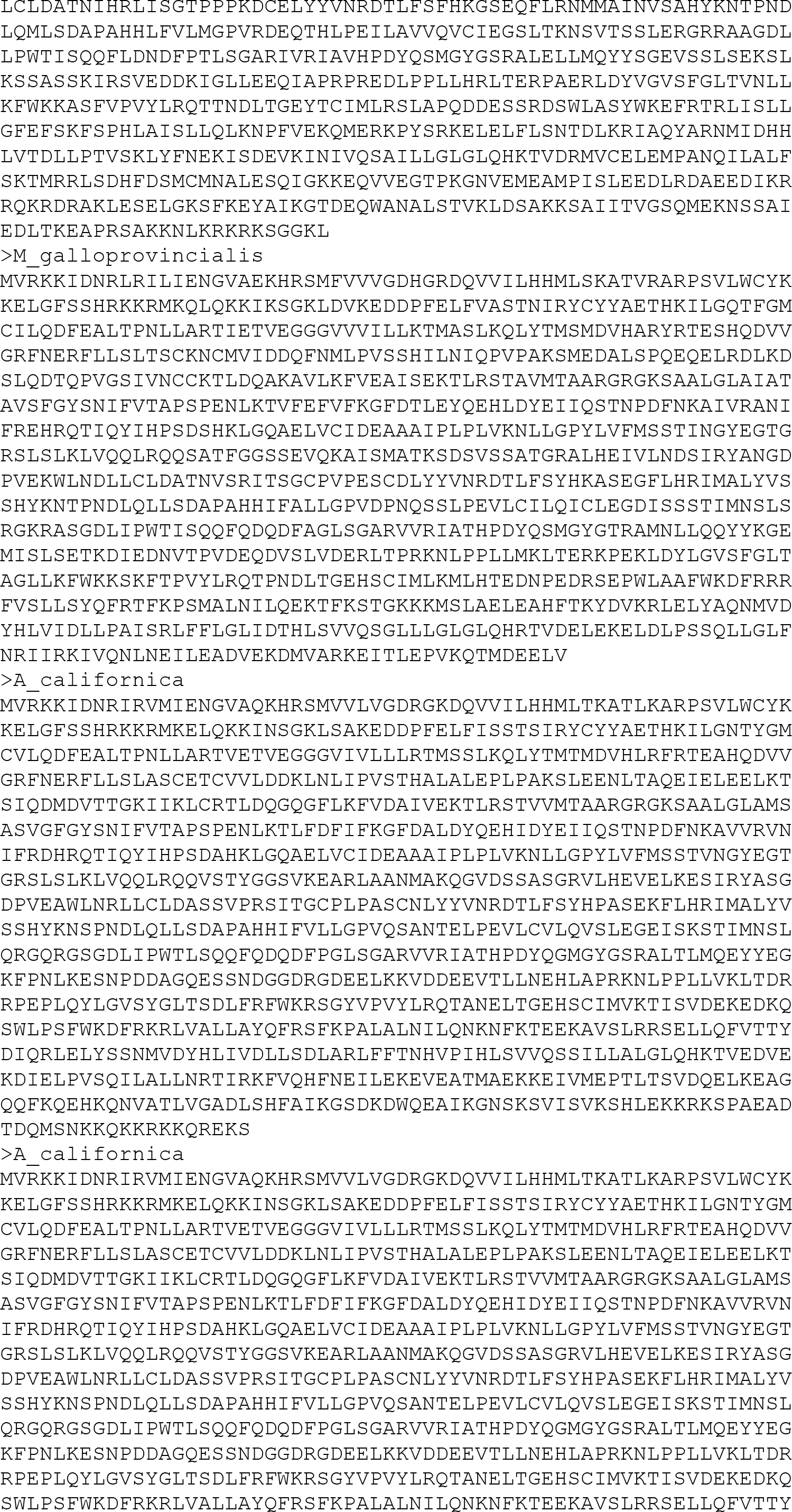
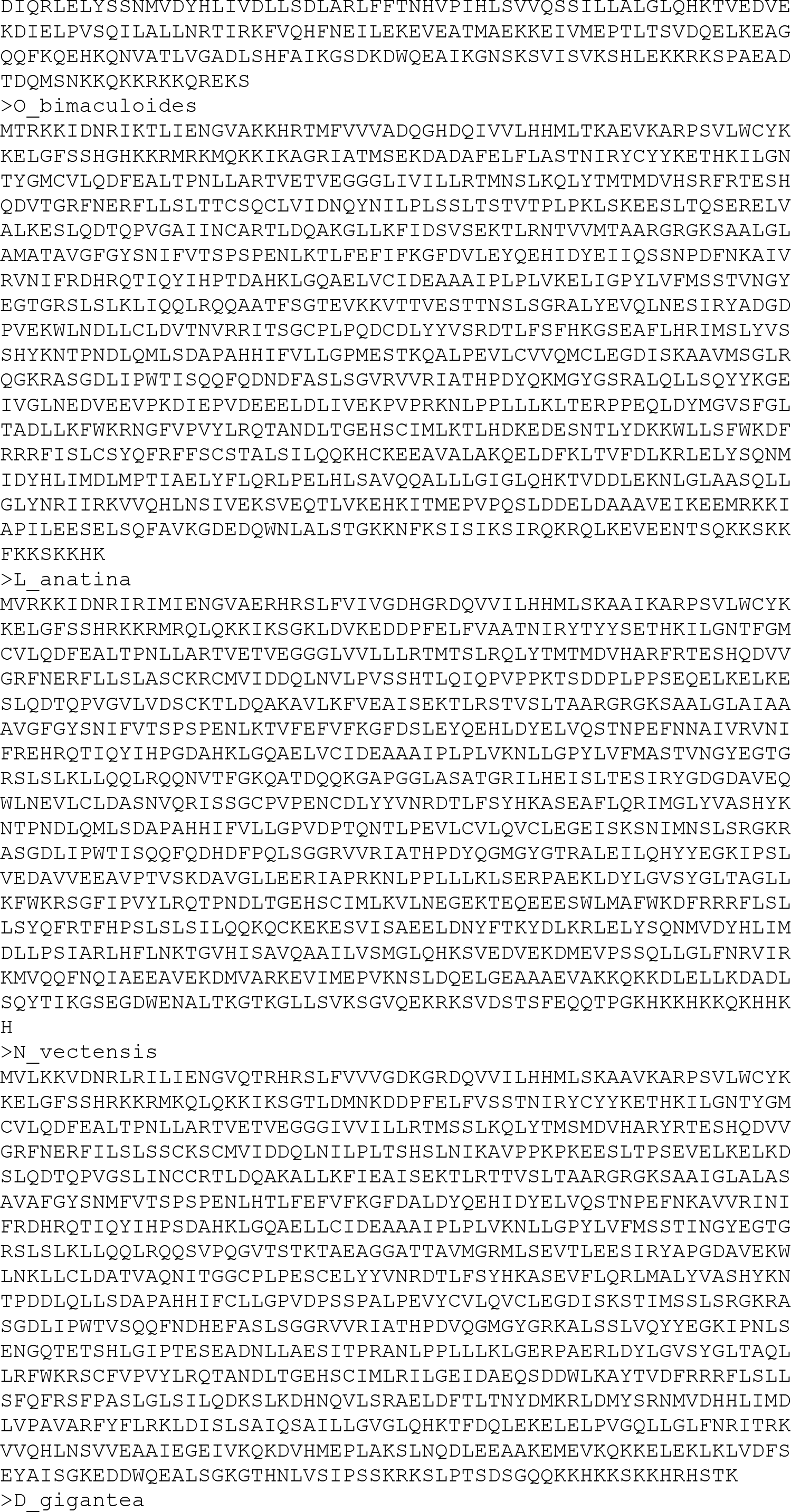
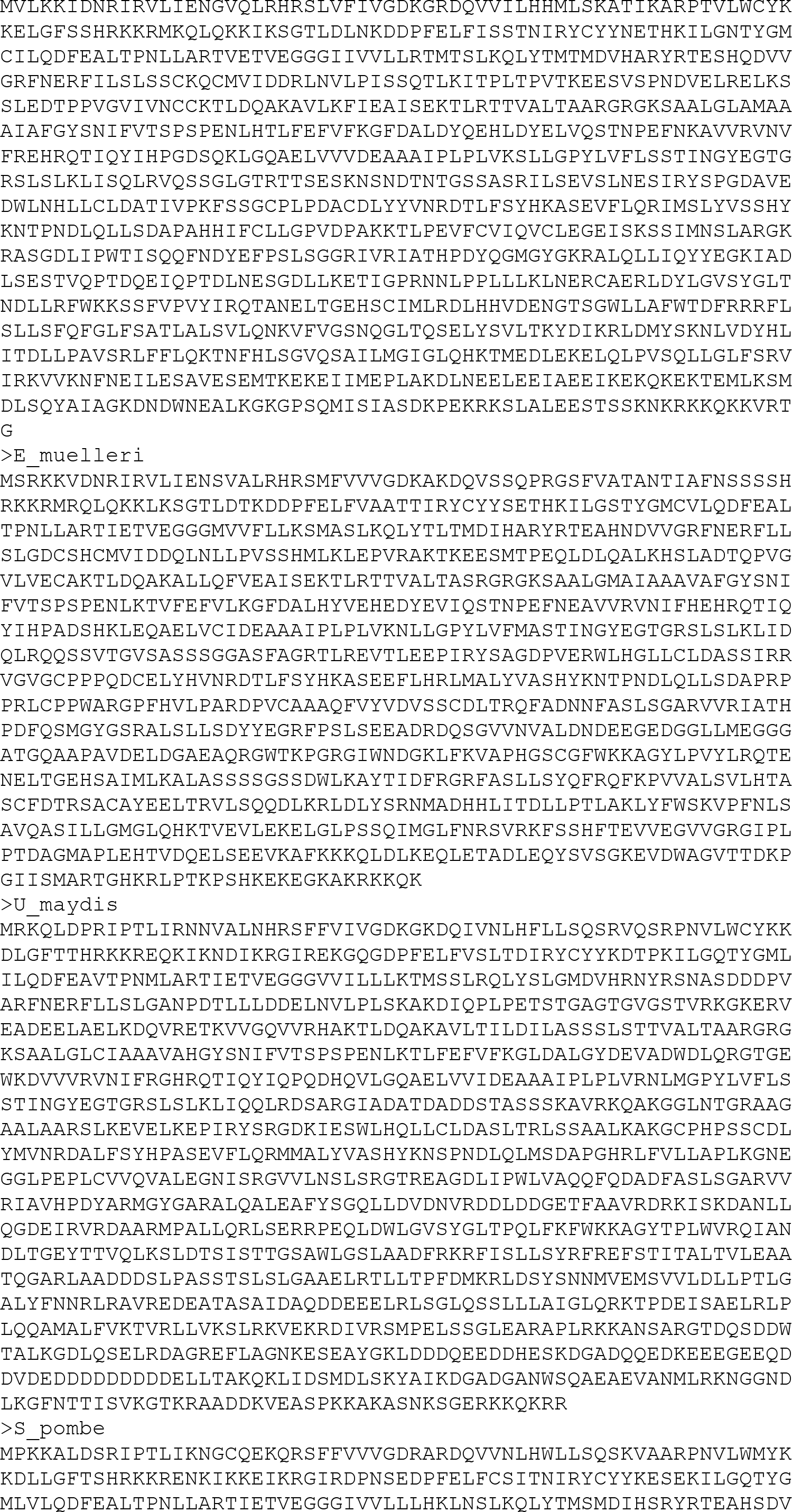
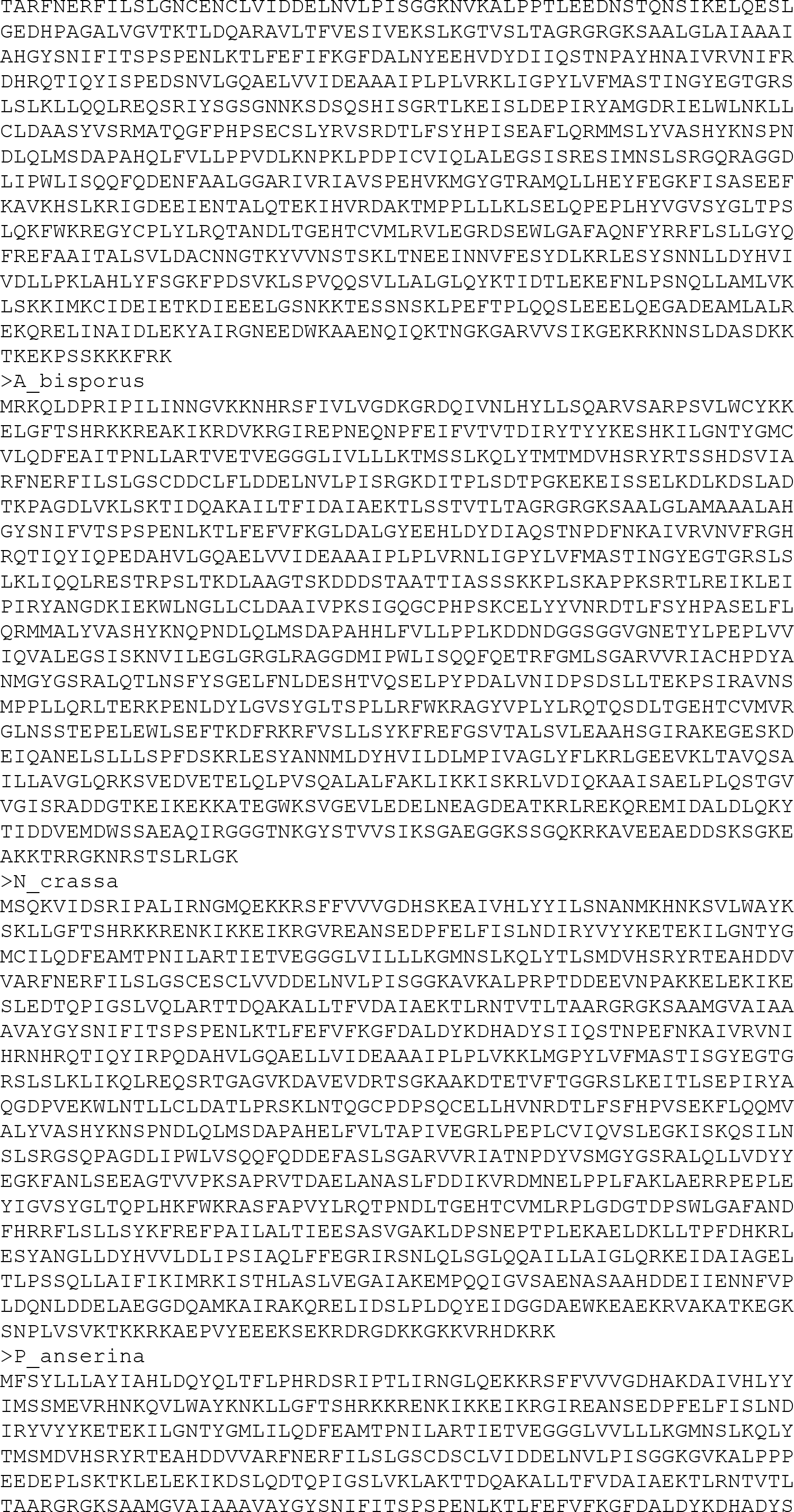
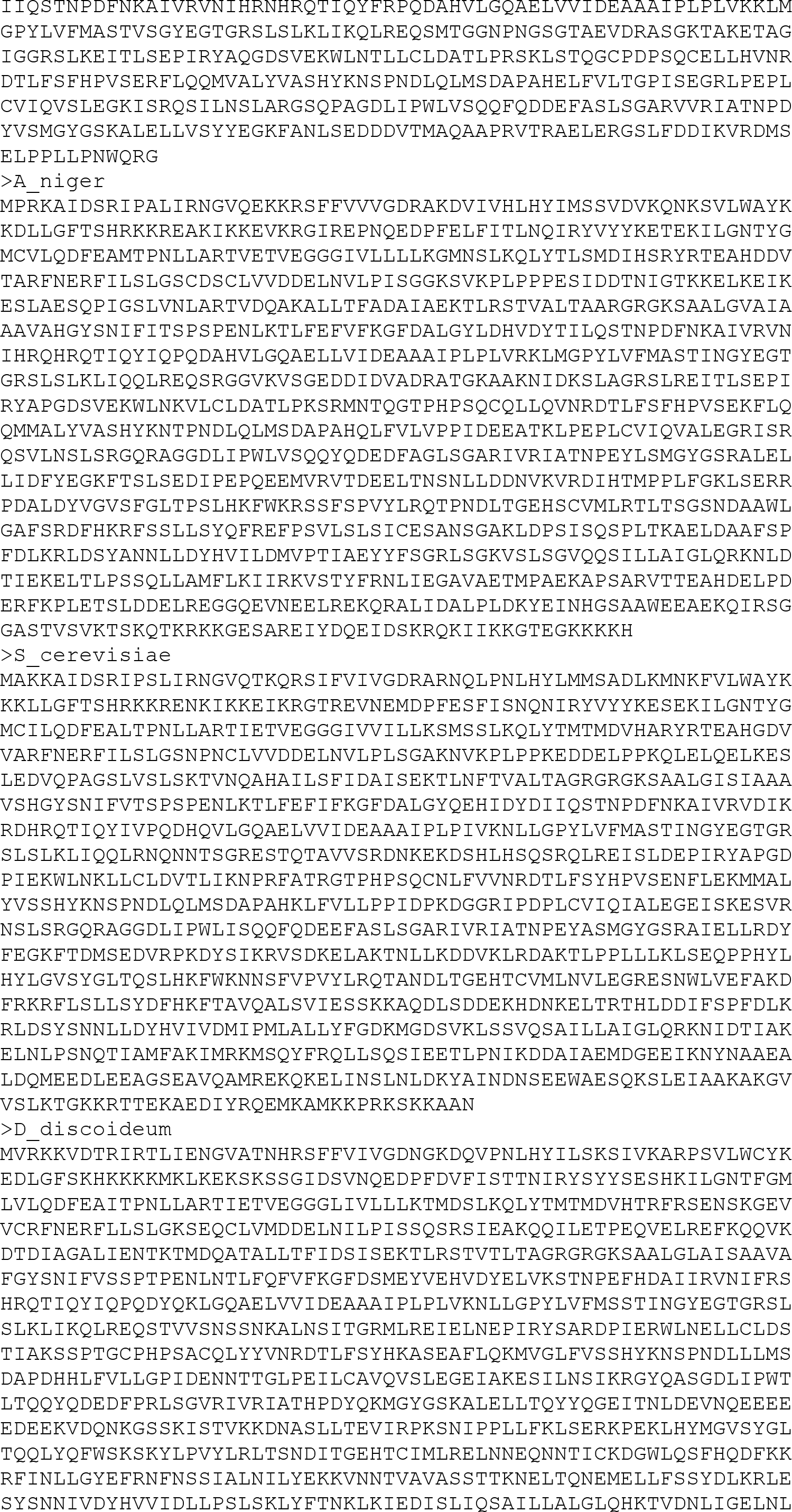
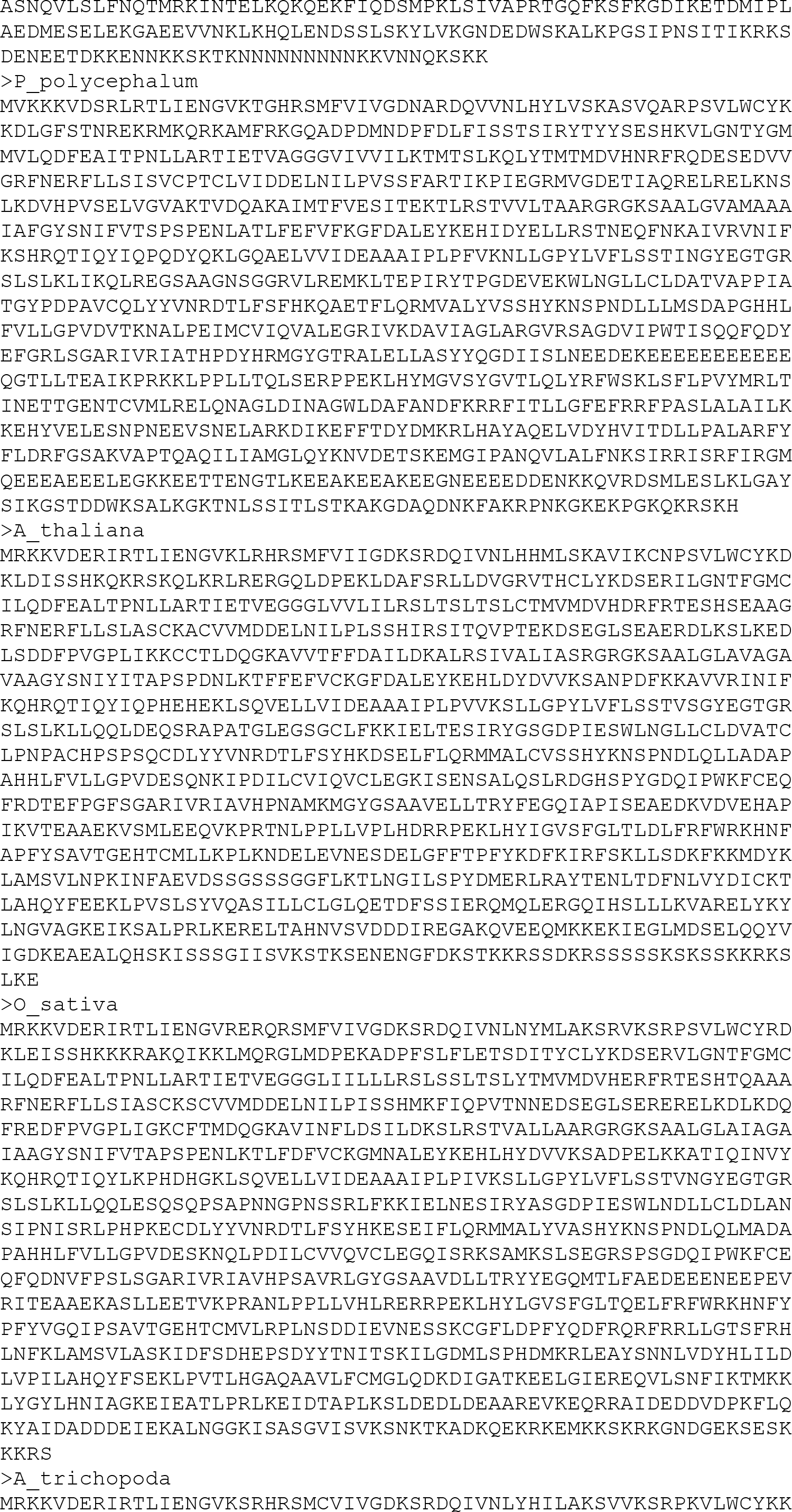
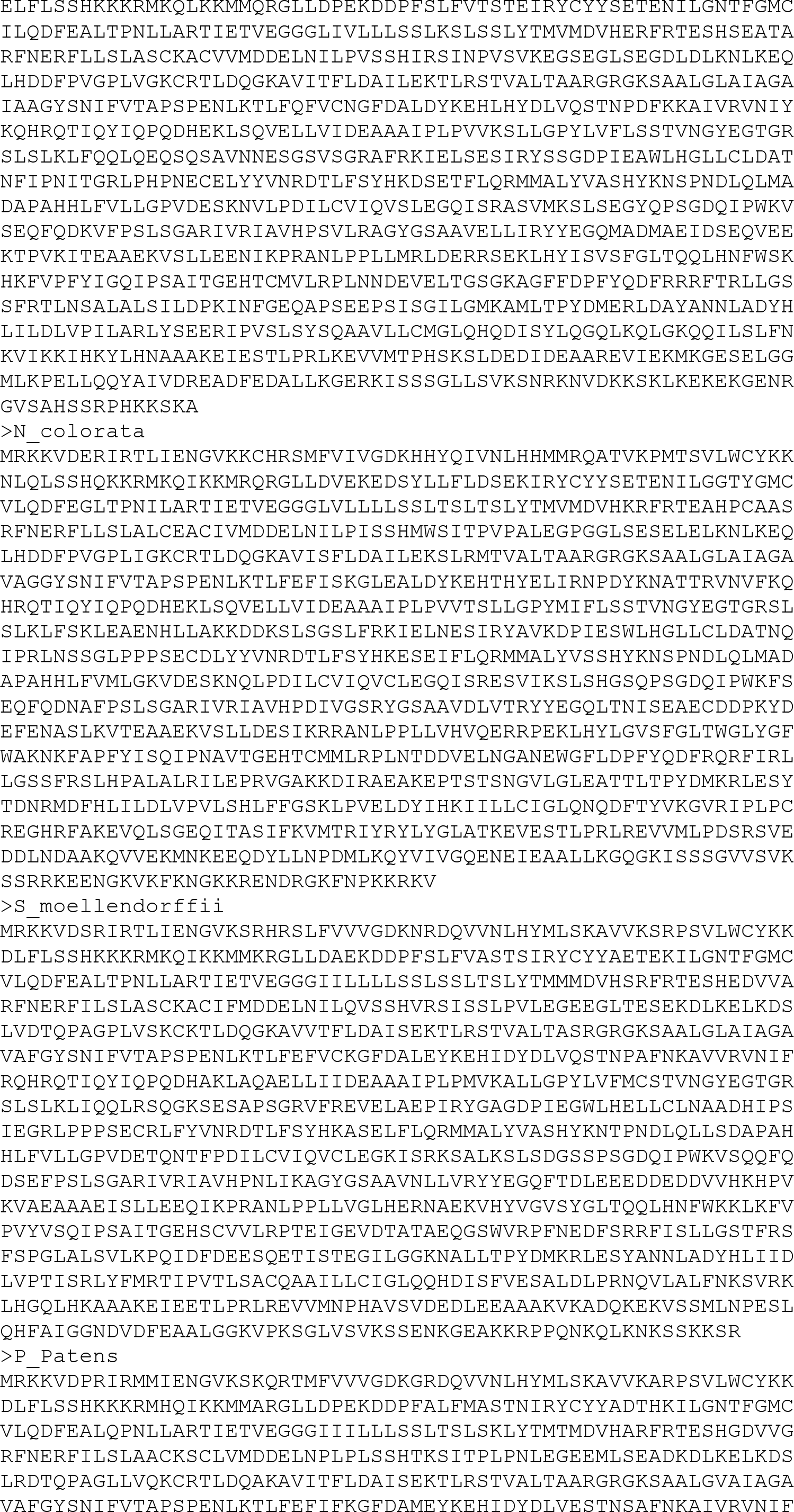
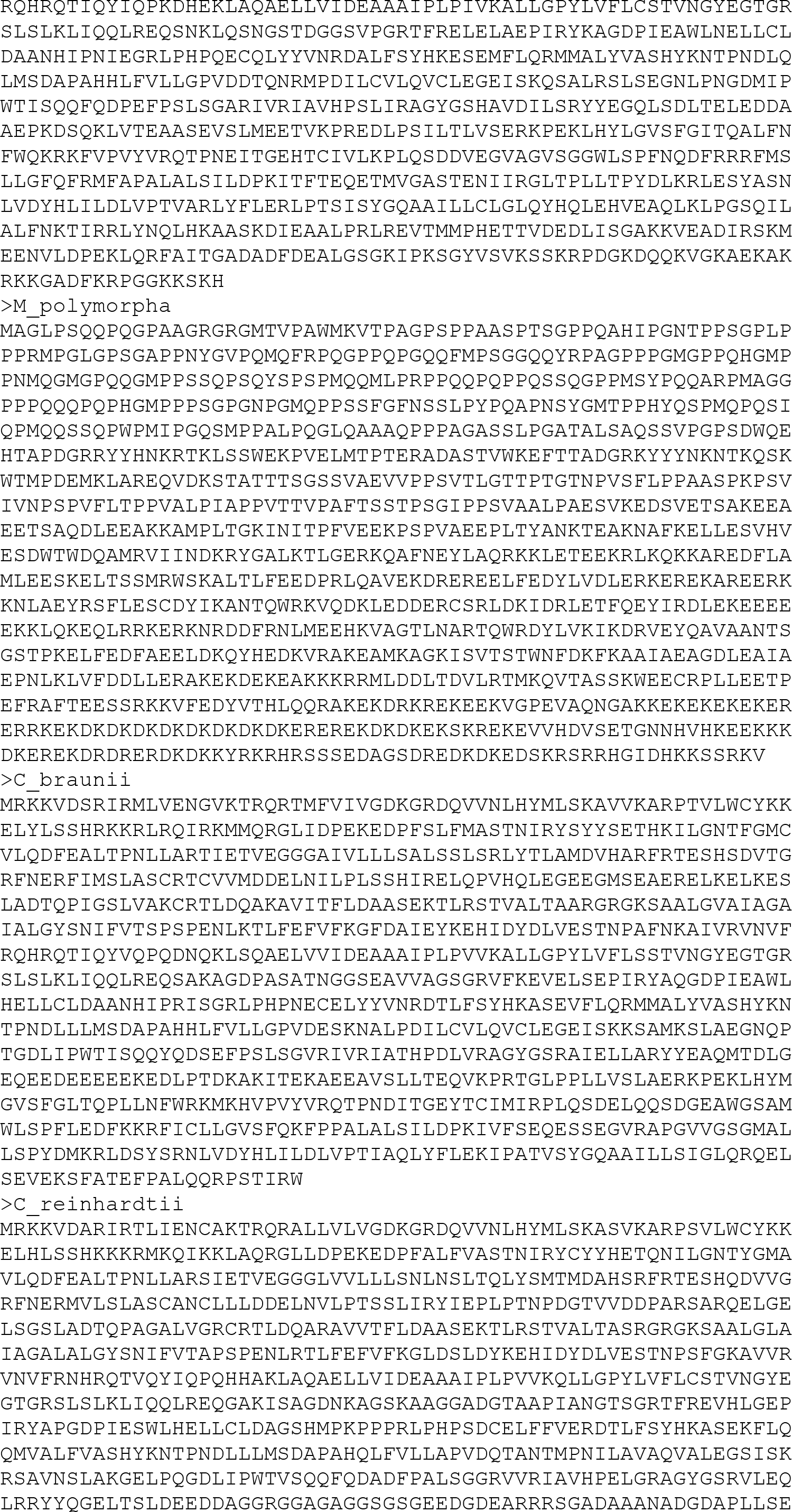
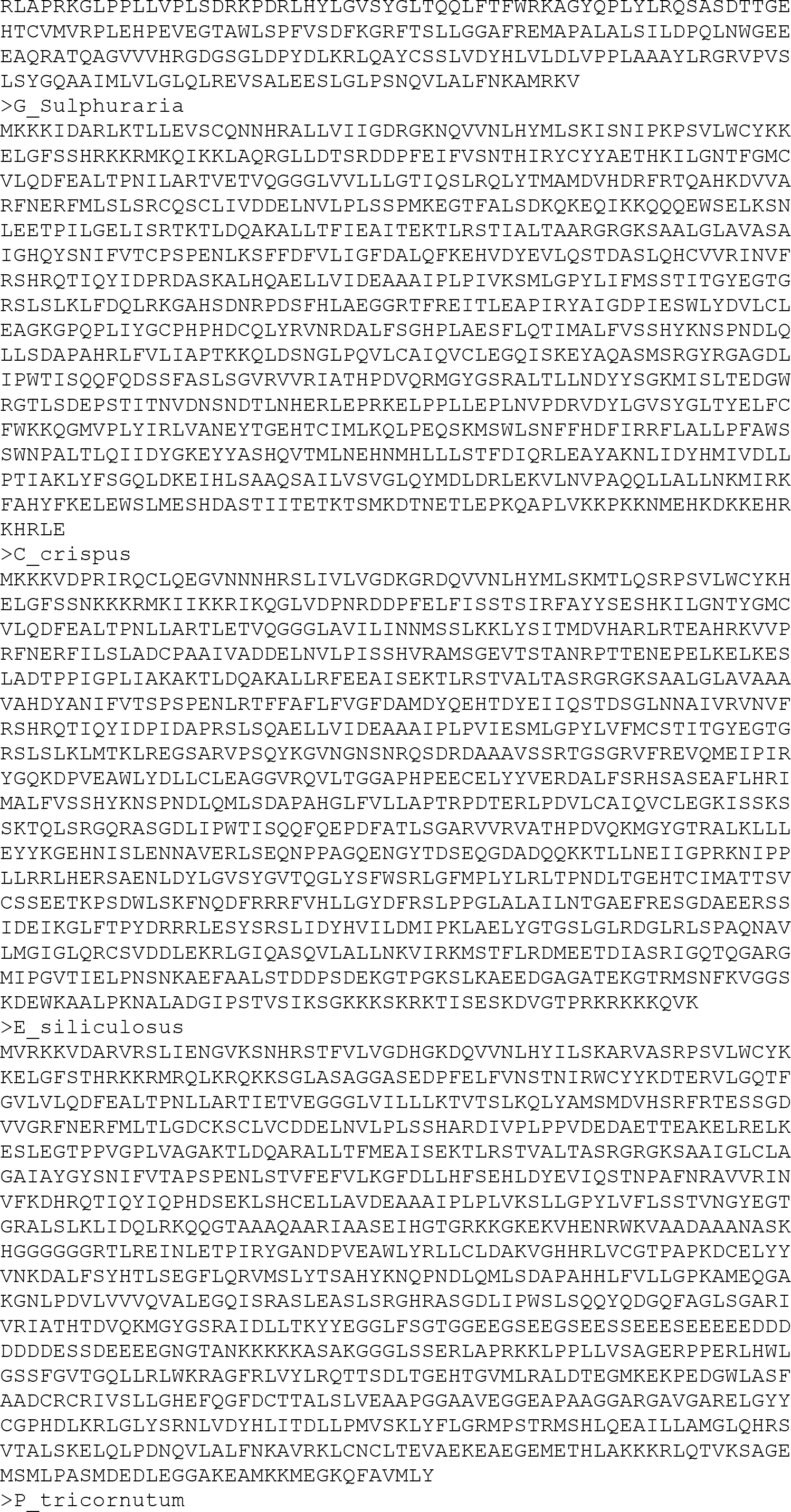
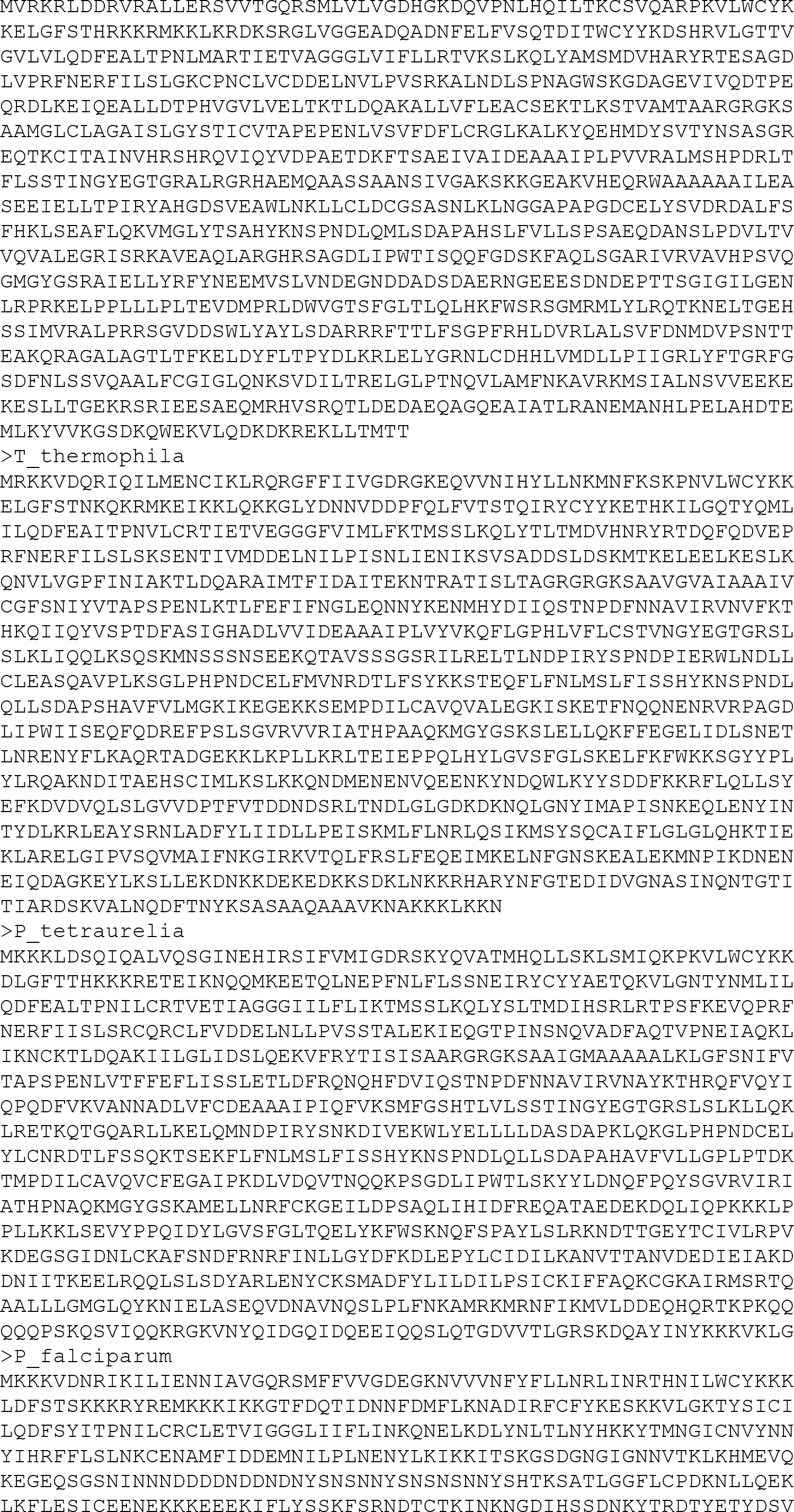
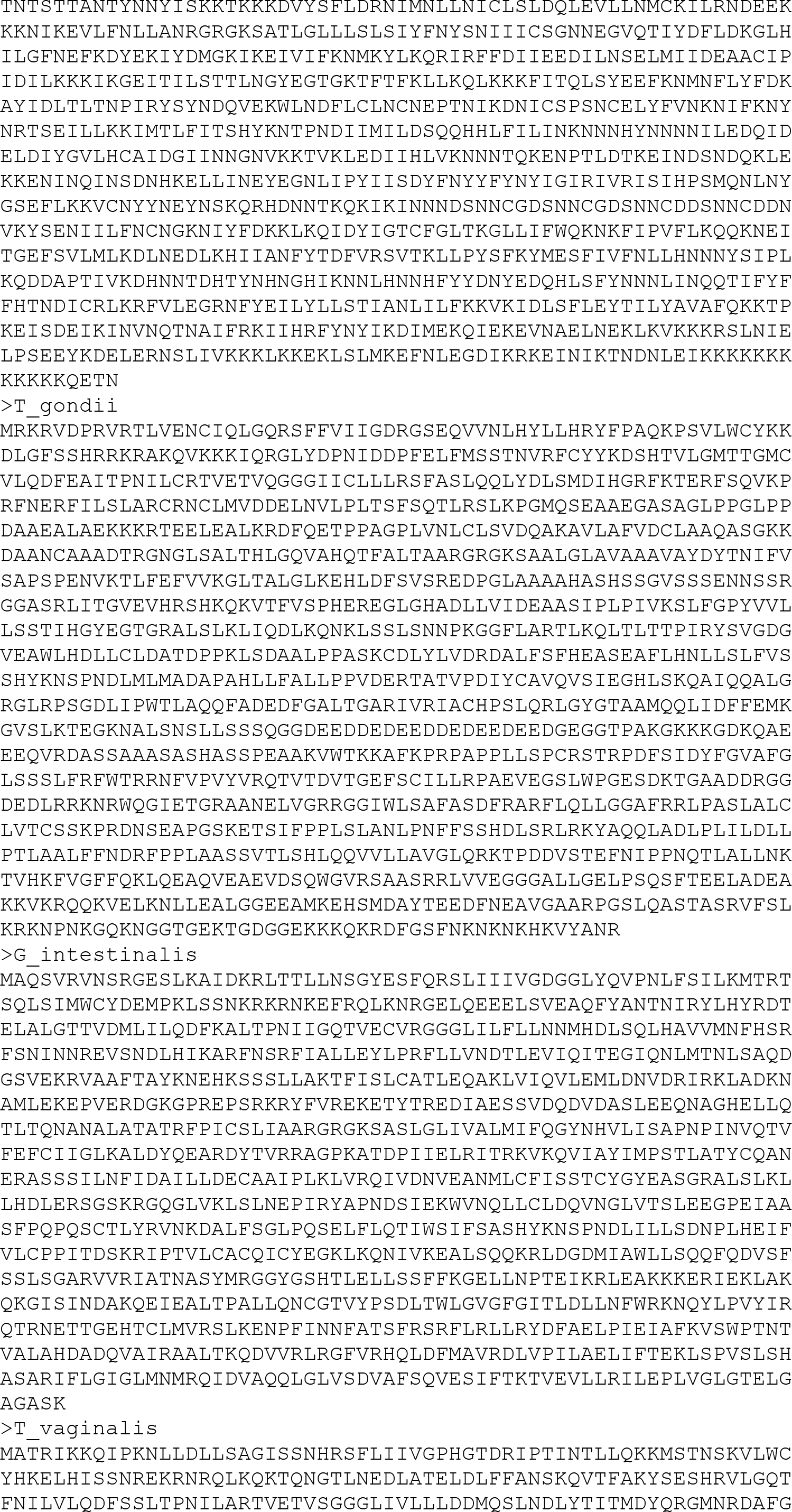
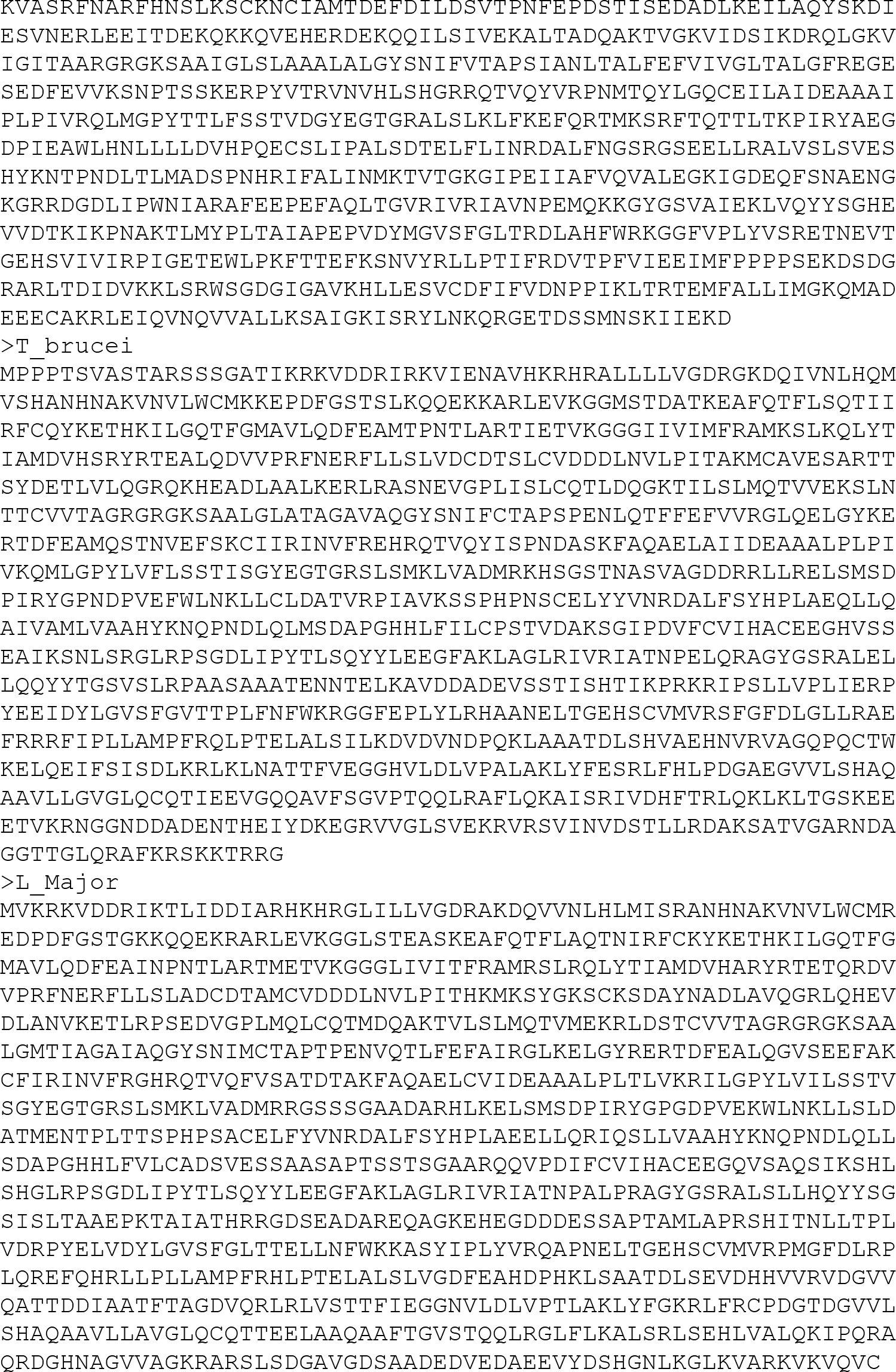
The primary structure of NAT10 across evolution.

**Supplementary data S3.**
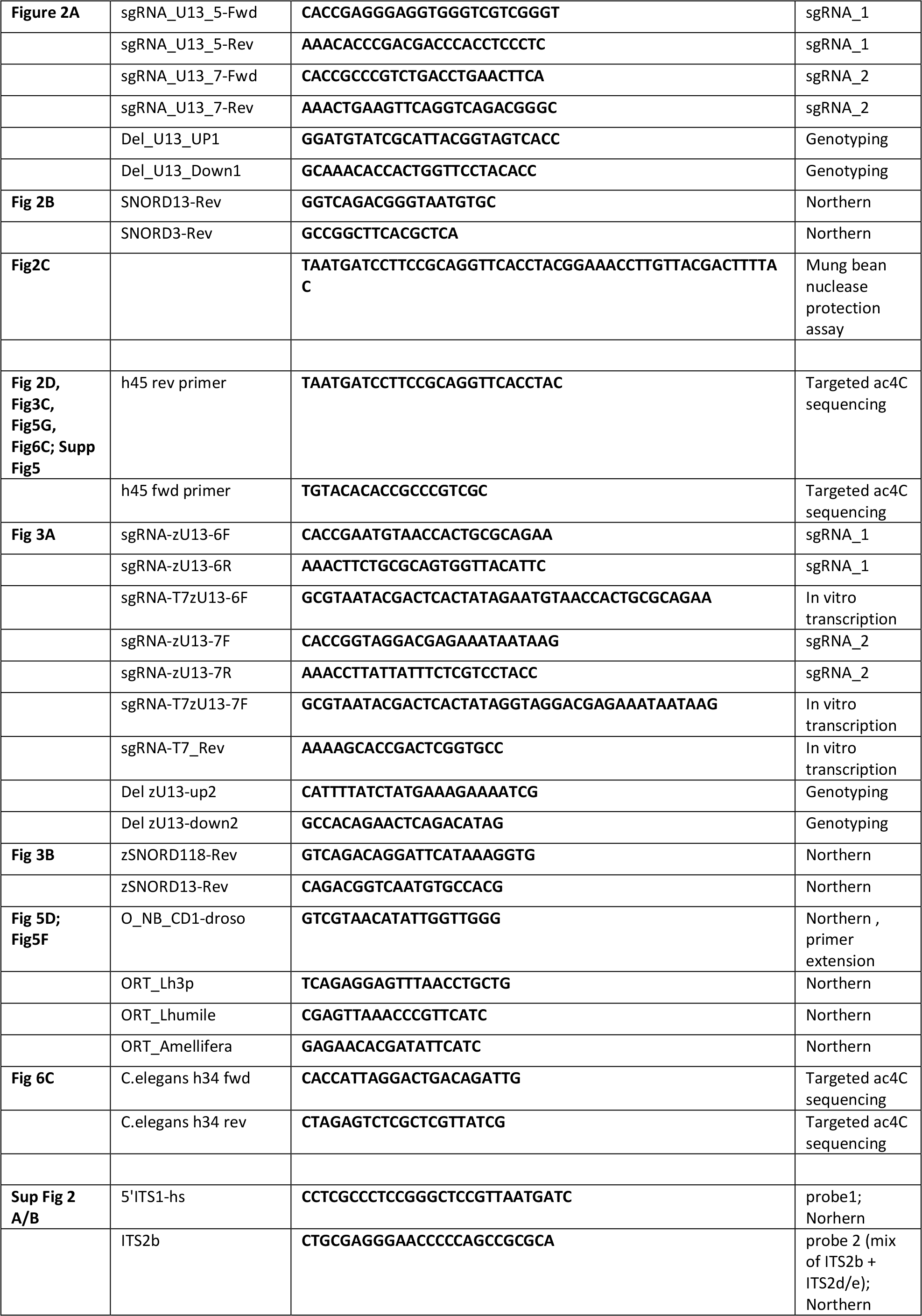
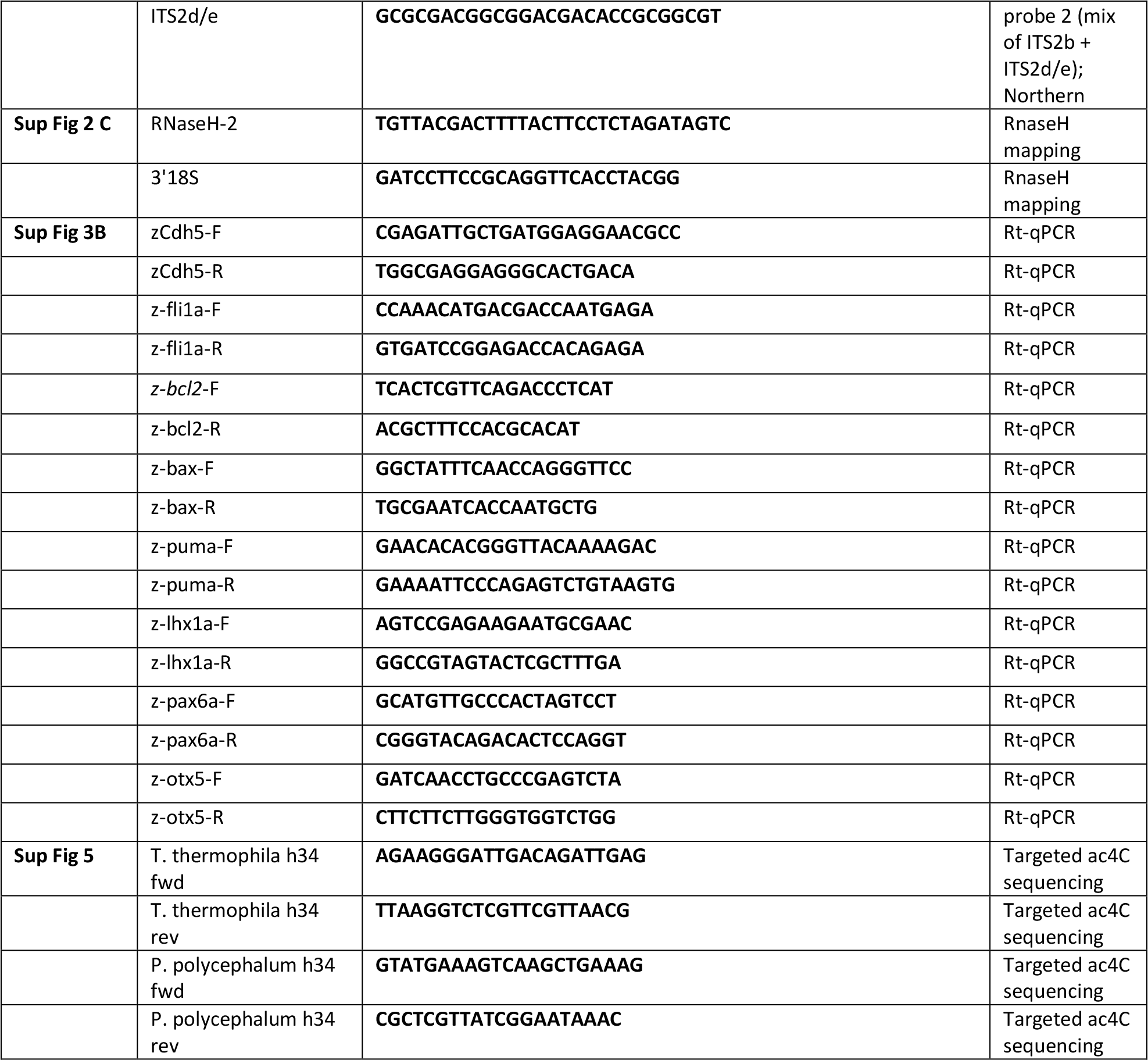
Sequences of primers.

